# Ecology and not phylogeny influences sensitivity to climate change in Muscicapidae Flycatchers in Eastern Himalayan and Indo-Burman hotspots

**DOI:** 10.1101/2023.10.05.561030

**Authors:** Aavika Dhanda, Michał T. Jezierski, Tim Coulson, Sonya Clegg

## Abstract

**Aims:** Species’ distributional responses to climate change can depend on ecology and phylogeny. The degree of habitat specialism is potentially important because habitat generalists with wider distributions are assumed to be less sensitive to environmental changes compared to narrowly-distributed habitat specialists. Additionally, predicting which aspects of the changing climate impacts species with different habitat associations may be particularly challenging in complex environments such as those of tropical mountains. We explore the effects of climate change on potential distributions of Muscicapidae Flycatchers in parts of the Eastern Himalayan Mountains and the Indo-Burman range, and relate predictions to the level of ecological habitat specialism, while accounting for phylogenetic relatedness.

**Location:** Bhutan and Northeast India

**Methods:** We used Maxent to develop ecological niche models of 60 Muscicapidae Flycatchers under different climatic scenarios by collating presences from Global Biodiversity Information Facility, Bioclimatic variables from WorldClim, and elevation from ASTER. Species were scored as habitat specialists or generalists using the Species Specialism Index. Variables with high contributions to Maxent models were extracted to explore sensitivity to climate change based on habitat specialism while testing for phylogenetic signal using Phylogenetic Generalised Least Squares.

**Results:** Maxent models had the highest contributions from variable bio8 (mean temperature of wettest quarter) under present climate, and tmax (maximum temperature) under future climatic scenarios. Phylogenetic Generalised Least Squares revealed that habitat generalists had higher sensitivity to climate change than specialists. We did not detect strong phylogenetic signal in sensitivity to abiotic variables under all climate scenarios.

**Main conclusions:** Potential distributions of Muscicapidae Flycatchers were sensitive to temperature variable in the month of the highest precipitation, and to maximum temperature. Potential distributions of habitat generalists were particularly sensitive to these abiotic variables, and those of habitat specialists less so. Sensitivity to abiotic variables did not show a pattern of phylogenetic niche conservatism. climate change, Muscicapidae Flycatchers, phylogenetic niche conservatism, habitat specialism, tropical mountains, Maxent

## INTRODUCTION

Climate change has implicated in biological responses in distributions, seasonal activities and migration patterns of approximately half of the species globally (IPCC, 2022), resulting in range shifts, adaptation, phenological tracking, or extinction in some species (Chen et al., 2011; Radchuk et al., 2019; Cleland et al., 2012; Maclean and Wilson, 2011, respectively). Out of these responses, latitudinal, altitudinal range shifts, and changes in species’ local distributions remain most widely studied and predicted (reviewed in Elith and Franklin, 2017). There are several factors such as dispersal ability, biotic interactions, phenology, available environmental space, and habitat specialisation which shape species’ distributions, and thus their ability to respond to climate change (Urban et al., 2013; Ponti and Sannolo, 2022; Enriquez-Urzelai et al., 2019, respectively). Although data on a majority of such ecological factors remain unavailable, habitat specialisation can be often quantified and used to make predictions related to distribution patterns (e.g., Lira et al., 2020). It is expected that habitat specialists, including the uncommon and high-elevation endemics with narrow ecological requirements are more likely to experience a decline in their distributions, whereas generalists capable of occupying several habitat types may not be severely impacted or could even be benefitting from climate change (review in Şekercioğlu et al., 2012).

Predicting responses to climate change may not be as straightforward for species occupying complex environments such as the tropical mountains. Tropical mountains are biodiversity hotspots (Myers et al., 2000) and are highly diverse ecosystems comprising of complex topographies and habitats contributing to several ecological and evolutionary patterns and processes (reviewed in Rahbek et al., 2019), thereby mediating heterogeneous responses to climate change in species (Tovar et al., 2013). Even though tropical mountains have long provided stable refuge to species (Fjeldså et al., 1999), these systems are now fast changing which is expected to result in high rates of range shifts and biotic attrition (Colwell et al., 2008).

Expectedly, habitat specialists tend to be more sensitive to climate change but there are variations in predictions from different climatic components affecting species’ distributions in tropical mountain systems. For example, forest specialist and insectivore bird species are more sensitive to increasing minimum temperatures than generalists with omnivore and herbivore guilds in tropical mountains of Africa (Dulle et al., 2016), whereas in the mountains of Costa Rica, habitat specialist bird species are more sensitive to both temperature and precipitation changes (Gasner et al., 2010). The sensitivity to abiotic environments can aid in an informed understanding of species’ niches and in identifying areas of stable refugia for spatial conservation prioritisation in the light of climate change (Keppel et al., 2012).

The patterns in predictions can be further explained by phylogenetically-explicit frameworks. Since closely related species are expected to have similar niches, commonly referred to as ‘phylogenetic niche conservatism’ (Losos, 2008), they may exhibit similar behavioural, ecological and life history traits (Kamilar and Cooper, 2013; but see Schuetz et al., 2019). Phylogenetic niche conservatism may explain various patterns in closely-related species such as divergence of foraging niches to avoid competition (Lovette and Hochachka, 2006), maintaining ancestral adaptation to certain environmental conditions (e.g., Morinière et al., 2016), and similarity in diversity-climate relationships (e.g., Wang et al., 2021).

Thus, phylogenetic relatedness needs to be accounted for to facilitate an improved understanding of species distributions, especially at regional scales (e.g., Pashirzad et al., 2018). Assessing phylogenetic relatedness can also have conservation implications as the climate-vulnerable traits, once identified in a species, can be extended to make assessments on its closely-related species (Miller-Rushing et al., 2010). However, factors like high topographical variation and altitude may lead to distinct phylogenetic signal in closely related species such as those occupying highlands and lowlands of tropical mountains (e.g., Schwallier et al., 2016).

We explore the potentially complex responses to climate change in Muscicapidae Flycatcher birds occupying tropical mountain environments. We focused on the Muscicapidae family of passerines which includes flycatchers, thrushes, and chats having variations in habitat specialism, high phylogenetic diversity (Sangster et al., 2010), and recent well-resolved phylogeny (Zhao et al., 2023). This provides us an opportunity to test for phylogenetic niche conservatism in closely-related species at a large spatial scale in a complex tropical mountainous environment. We characterised species-specific potential distributions of 60 species by developing Ecological Niche Models (ENMs) under both present and future climatic conditions, in parts of two global biodiversity hotspots-the Eastern Himalayan Mountains and the Indo-Burman Hills.

Parts of the Eastern Himalayan Mountain and Indo-Burman Hills-two global biodiversity hotspots (Mittermeier et al., 2011) encompass Bhutan and Northeast India which are located directly above the Tropic of Cancer at 21.94º - 29.46ºN and 97.41º - 88.01ºE (Figure 1). The Indo-Burma extends across southern parts of Northeast India, with plains of Brahmaputra in between-one of the largest rivers of Asia. The region provides highly heterogeneous environments with several altitudinal gradients (roughly from 10 metres to 7000 metres above sea level), varying climatic conditions and multiple habitat types, thereby facilitating high diversity in species such as of birds (Acharya et al., 2011), plants (Saikia et al., 2017), mammals (Dhendup et al., 2019), and butterflies (Singh and Chib, 2014). Bhutan and Northeast India is also faced with severe effects of climate change resulting in faster melting of glaciers, increased frequency of floods, and decreased seasonal rainfall in some parts (Borah et al., 2022; Das, 2016; Choudhury et al., 2019, respectively), which is expected to have severe consequences on the biodiversity of the region.

**Figure 1:**
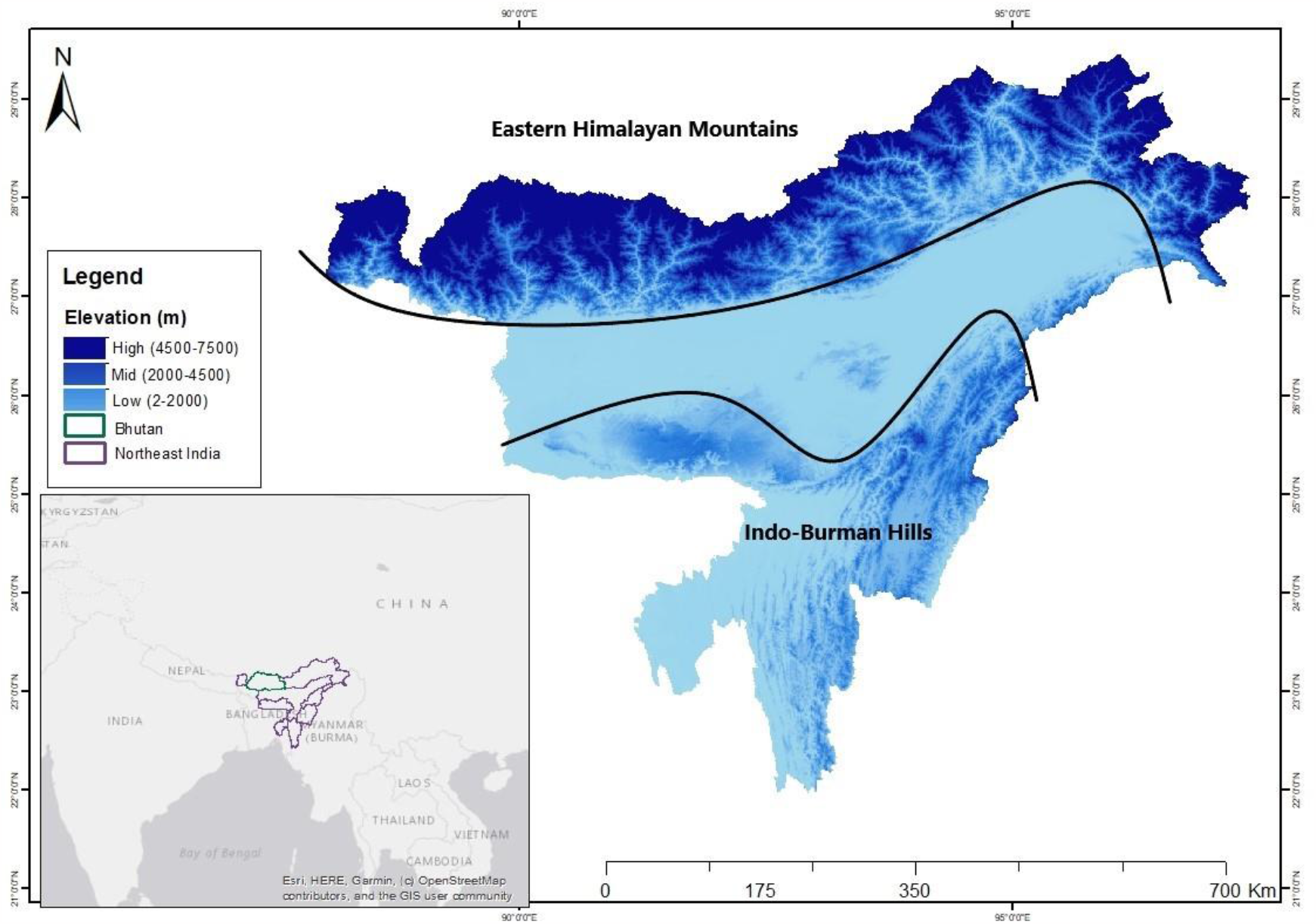
Bhutan and Northeast India with high elevation Eastern Himalayan mountains in the north, low to mid elevation Indo-Burman Hills in the south, and Brahamputra river plains in the middle

Our first aim was to determine potential distributions of the Muscicapidae Flycatchers in the region. Species’ potential distribution or the fundamental niche reflects the abiotic conditions best suited for species (reviewed in Peterson and Soberón, 2012). We used Maxent, a widely used machine-learning program that predicts species’ potential distributions based on presence-points to develop ENMs representing habitat suitability (Phillips et al., 2006). We then extracted variables with the highest contributions to the ENMs which explained abiotic sensitivity for all species. We expected that the potential distributions of species will be most affected by not only temperature, but by a combination of temperature and precipitation variables under both present and future climatic conditions, since the region of Eastern Himalaya and Indo-Burma offers a complex suite of abiotic environments.

We then investigated the role of species’ habitat specialism and evolutionary histories in explaining the patterns in potential distributions. We applied the Phylogenetic Generalised Least Squares method (PGLS) (Grafen and Hamilton, 1997) which explicitly incorporates phylogenetic non-independence between species with shared evolutionary histories (Symonds and Blomberg, 2014). We used the extracted abiotic variables with the highest contribution as a response variable and habitat specialism as the predictor variable while controlling for phylogeny in PGLS models. We then explored phylogenetic signal to assess if closely-related species are likely to show similar sensitivity to abiotic variables under both present and future climatic scenarios in highly complex environments in parts of the Eastern Himalayan and the Indo-Burman hotspots. We expected that the habitat specialists will be more sensitive to these abiotic variables.

## METHODS

### Presence records

A list of Muscicapidae Flycatchers residents and migrants in our study region, excluding vagrant species found outside their home ranges, was collated from eBird (Sullivan, et al., 2009), and Birds of the Indian Subcontinent: India, Pakistan, Sri Lanka, Nepal, Bhutan, Bangladesh and the Maldives (Grimmett et al., 2012) resulting in a list of 79 species from 26 genera. We collated presence records from the Global Biodiversity Information Facility (GBIF; https://www.gbif.org/), a biodiversity portal offering free and open-access geo-tagged species records (Telenius, 2011). Three categories of ‘basis of records’ from GBIF database were extracted: human observation, observation, and living sample, as these represent direct field observation records. Data wrangling and analyses was performed in RStudio Desktop version 2023.03.0+386 using R base software version 4.0.4 (R Core Team, 2020). Data extraction and cleaning was conducted using `dismo` package (Hijmans et al., 2020), and missing and duplicate coordinates and those outside the geographic extent of our study area were removed using `tidyverse` (Wickham, 2021). Species with less than 10 presence records and those with taxonomic uncertainties were eliminated at this stage. We thus proceeded with 60 species for developing ENMs and performing subsequent PGLS analysis.

### Bioclimatic variables

Gridded 19 bioclimatic variables for historical or near current climatic conditions were downloaded from WorldClim (https://worldclim.org/) at a spatial resolution of 2.5 minutes (Fick and Hijmans, 2017). To check for, and remove bioclimatic variables with high collinearity, we performed Variance Inflation Factor (VIF) analysis (Thompson et al., 2017) using `usdm` package (Naimi, 2017). Variables with VIF values greater than 10 were removed (Yoo et al., 2014), yielding seven variables: bio2, bio3, bio8, bio14, bio15, bio18 and bio19 (Table 1).

**Table 1.** Seven Bioclimatic predictor variables for current climate scenario retained after performing Variance Inflation Factor (VIF) test of collinearity.

| Codes | Description | VIF values |
| --- | --- | --- |
| `bio2` | Mean diurnal range (mean of monthly (max temp- min temp)) | 3.466 |
| `bio3` | Isothermality x 100 | 1.711 |
| `bio8` | Mean temperature of wettest quarter | 2.160 |
| `bio14` | Precipitation of driest month | 6.290 |
| `bio15` | Precipitation seasonality (coefficient of variance) | 1.738 |
| `bio18` | Precipitation of warmest quarter | 3.077 |
| `bio19` | Precipitation of coldest quarter | 7.509 |

ASTER Global Digital Elevation Model was obtained for the study area from United States Geological Survey AppEEARS website (https://lpdaacsvc.cr.usgs.gov/appeears/) at a spatial resolution of 30 metres and resampled in ArcGIS Desktop 10.6.1 Advanced (ESRI, 2018). Elevation was used as the eighth predictor variable in addition to those in Table 1.

Future climatic gridded data for the next two decades (2021-2040) was obtained from WorldClim at a spatial resolution of 2.5 minutes for which we chose Beijing Climate Center Climate System Model BCC-CSM2-HR, a global circulation model downscaled from CMIP6 and widely used for Asia (Eyring et al., 2016). Four climate change scenarios described by Shared Socio-economic pathways (SSPs) representing different possible future greenhouse gas emissions were SSP1-2.6 (low emission scenario), SSP2-4.5 (intermediate emission scenario), SSP4-6.0 (medium emission scenario), and SSP5-8.5 (high emission scenario) (van Vuuren et al., 2011; Thomson et al., 2011; Masui et al., 2011; Riahi et al., 2011, respectively) in which the global mean temperature is projected to increase by 0.9°C, 1.1°C, 1.0°C and 1.3°C for 2021-2041, respectively (IPCC, 2019). We used four variables for each of the four SSPs - maximum temperature (tmax), minimum temperature (tmin), and precipitation (prec), also obtained from WorldClim, along with elevation from ASTER.

### Maxent settings for ENMs

Maxent is one of the best performing models for presence-only data (Elith et al., 2006) and is frequently used to predict species distribution changes due to climate change (Zhang et al., 2018). Maxent compares environmental conditions where a species has been recorded to those of the auto-generated random ‘background points’ following the principle of maximum entropy using a set of restraints provided as ‘settings’ by the modeler (Phillips et al., 2017). We used Maxent.jar software to produce potential distribution maps for each species (https://www.cs.princeton.edu/). Maxent.jar tends to develop complex maps under default settings which can be avoided by manually selecting settings to furnish simpler maps (Halvorsen et al., 2016). Thus, we selected quadratic features to reflect species’ physiological response curve which generally represents a Gaussian-shaped unimodal curve (Austin, 2007), along with linear features to ensure Maxent does not choose background points for species from areas they are not likely to inhabit, such as those in freezing temperatures (Merow et al., 2013). This particularly applies to tropical species as they can have truncated niches and do not occupy full range of available climatic conditions (Smith et al., 2020).

We used 10 model replicates and set bootstrapping as the replicated run type since it assesses the likelihood of predictor variable being independent and also generates parsimonious predictive models (Austin and Tu, 2004). Random test percentage was set to 25% for model validation (Xu and Goodacre, 2018). We then explored the effect of varying maximum number of background points using 2,000, 5,000 and 10,000 points as suggested by Merow et al. (2013), but found no differences in present climate models. Hence, we proceeded with 10,000 background points for all present and future climate change models to maintain consistency.

Raw output was selected for all models which corresponds to relative occurrence rate (ROR) in Maxent (Merow et al., 2013), since presence only data can at best estimate ROR and not absolute occurrence rates (Fithian and Hastie, 2013; Guillera-Arroita et al., 2014). The maximum number of iterations was set to 500 and jackknife method was used to assess variable importance of all ENMs. Receiver operating characteristics analysis or area under the curve (AUC) was used to estimate the accuracy of the models, which is considered a relatively better measure of predictive accuracy in comparison to other criteria such as Naive Bayes Classifier and Decision Trees for evaluating and comparing machine learning algorithms (Huang et al., 2003). We selected a mean model of ten replicated models to compute AUC scores. AUC scores range from 0 to 1 to describe how well a model has performed by discriminating between sensitivity (presence correctly predicted as presence) and specificity (absence correctly predicted as absence) such that a score above 0.5 is considered well-performing and those below 0.5 are poor-performing (Jiménez-Valverde, 2012). Furthermore, previous studies have highlighted strong relationship between AUC scores and species rarity, indicating species with fewer presence points are likely to have a high AUC scores and thus have better performing models (Connor et al., 2018; Veloz, 2009). We explored if such a relationship exists between AUC scores and total number of presence points in our focal species by fitting a Generalised Linear Model.

### Phylogenetic analysis

#### Percentage Contribution of Variables-response variable

We used Percentage Contribution of Variables (PCVs) from ENMs of each species as a proxy for sensitivity to climate change under both present and future climatic scenarios. PCVs are continuous percentage measures of variable importance. We extracted means of PCVs of 10 bootstrapped models for each of the species from Maxent output file called ‘maxentresults’. Out of the eight variables used under present climate, i.e., bio2, bio3, bio8, bio14, bio15, bio18, bio19 (see Methods Table 1), and elevation; bio8 (mean temperature of wettest quarter) and elevation had the highest percentage contribution for most species. From the variables used for four future climate scenarios, i.e., tmax, tmin, prec, and elevation, the variables tmin and tmax had the highest percentage contribution in ENMs of most species under each of the four scenarios (for detailed PCVs, see Supplementary material Table S1 and Table S2). Thus, percentage contribution of bio8 and of elevation under present climate, and tmax and tmin under future climate scenarios were selected for subsequent PGLS modelling.

#### Species Specialisation Index - explanatory variable

There are several habitat specialisation metrics used for birds, however, most are based on abundances, such as the Rank occupancy-abundance Profile (Collins et al., 2009) or Marsh

Specialisation Index (Correll et al., 2016), and on expert opinion (e.g., Gardali et al., 2012a); all frequently limited for the understudied systems. We used Species Specialisation Index (SSI) which orders species from specialists to generalists based on the variances of average densities of species in the total number of habitats without having to rely on expert knowledge (Julliard et al., 2006). In order to score each species according to its habitat preferences, we extracted information from Birds of the World (2023) and categorised all the habitat types in 10 broad classes that the Muscicapidae Flycatchers in the study region are found inforests, secondary forest, bamboo, scrub, reeds, riverain forests, rivers and streams, alpine, rocky slopes, and open. We then calculated SSI for 60 species across 10 habitats, where *h* represents number of habitat classes a species is found in, and *H* represents total habitat classes. (1).

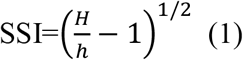

The index was then normalised on the scale of 0 to 1, termed as *S’*, such that (*S’*→ 0) signifies species is an extreme habitat generalist, e.g., *Copsychus saularis* (Oriental Magpie-Robin) (found in 6 out of 10 habitats) while (*S’*→ 1) signifies species is an extreme habitat specialist, e.g., *Niltava grandis* (Large Niltava) (found in 1 out of 10 habitats).

### Phylogenetic modelling

We obtained complete phylogeny of 60 species for Muscicapidae Flycatchers family built using mitochondrial and nuclear data from Zhao et al. (2023). We used the `ape` package to read, plot and manipulate the phylogenetic tree (Paradis and Schliep, 2019), and gls function in `nlme` package (Pinheiro et al., 2017) to fit multiple PGLS models. First, linear model residuals using PCVs and *S’* were tested for normality. Model residuals of bio8, elevation, and tmax under low, intermediate, medium, and high emission scenarios were normal. The remaining variable, i.e., tmin was log-transformed but did not conform to normality under any future emission scenario, hence was not used in the PGLS models.

To investigate the role of species habitat specialism in explaining the highest percentage contribution of abiotic variables in ENMs, we fit two PGLS models for present climatic conditions; one, using percentage contribution of bio8, and another using percentage contribution of elevation as response variable, and *S’* as a predictor variable in both the models. We repeated the same procedure with tmax for future climatic conditions. We used phylogeny as a fixed effect and the fixed variances in distances from root to tip of each tree as weights. The PGLS analysis was performed based on Brownian motion model (Felsenstein, 1985).

We measured the phylogenetic signal in contributions from abiotic variables based on the significant PGLS models using `phylosignal` package (Keck et al., 2016). Blomberg’s *K* was used to estimate the strength of phylogenetic signal such that *K*<1.0 represents less phylogenetic signal than expected and *K*>1.0 represents greater phylogenetic signal than expected under Brownian motion (Blomberg et al., 2003). We then estimated local Moran’s *I* score to detect for phylogenetic autocorrelation, where the values of *I* denote degree of autocorrelation between closely related species (Gittleman and Kot, 1990). Phylogenetic trees with annotations of PCVs resulting in significant PGLS models constructed using `ggtree` package (Yu et al., 2017).

## RESULTS

### Model performance and variable contributions

Model AUC scores ranged from 0.66 to 0.98 for present climatic conditions, and from 0.63 to 0.97 under four future climate change scenarios (Supplementary material Table S3). These values (all above 0.5) denote that the models performed well and were either sufficient, good or very good at discriminating between sensitivity and specificity (Šimundić, 2009). While all models performed well, generalised linear models predicted a significant negative relationship between AUC scores and the total number of presence points under present climatic conditions (*P* > |t| =3.36e-0.7), and all future climatic conditions - SSP 1-2.6 (*P* > |t| =5.45e-05), SSP 2-4.5 (*P* > |t| =5.13e-05), SSP 4-6.0 (*P* > |t| =4.21e-05), SSP 5-8.5 (*P* > |t| =5.21e-05). This relationship could be attributed to Maxent’s tendency to better discriminate between specificity and sensitivity for species with fewer presences as compared to those with several presence points (Stokland et al., 2011), since species with more presence points and thus wider distributions can make it challenging for Maxent to distinguish between suitable and unsuitable areas (Franklin et al., 2009).

The percentage contributions of bio8 and elevation varied amongst the species. Bio8 contributed the most to potential distribution of *Copsychus saularis*, and least to *Ficedula parva*, while elevation contributed the most to *Phoenicurus hodgsoni* and least to *Copsychus saularis*-all of which are habitat generalists (Figure 2). Under the four future climate scenarios, tmax consistently retained the most information in potential distributions of *Copsychus saularis*, a habitat generalist, and *Niltava grandis*, a habitat specialist (see Supplementary Figure S59 to S62).

**Figure 2:**
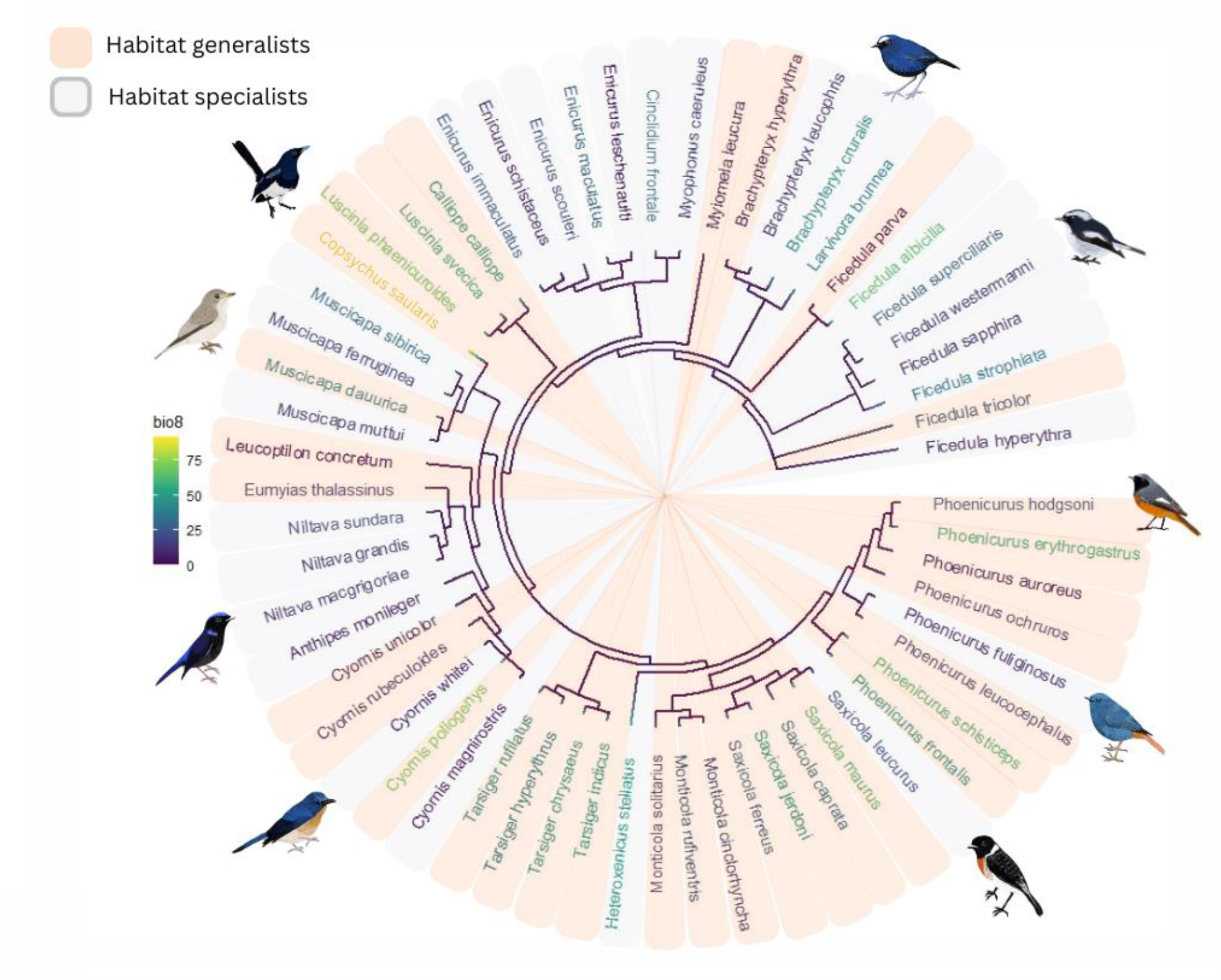
Percentage contribution of bio8 (Mean temperature of wettest quarter) of the under the present climate to Ecological Niche Models of Muscicapidae Flycatcher species categorised as habitat specialists (light grey) or generalists (light orange) based on Species Specialisation Index

### Potential distribution maps

The predicted potential distribution maps of an extreme generalist and an extreme specialist species, *Copsychus saularis* and *Niltava grandis*, respectively, are presented as exemplars. Under the present climate, *C. saularis* has high suitability along the plains and the Indo-Burman Hills, and low suitability in parts of Eastern Himalayan Mountains (Figure 3). *N. grandis* has low suitability in the plains, but medium to high suitability in parts of Eastern Himalaya (Figure 4). Under future climatic scenarios, the potential distribution of *C. saularis* remains largely unaffected, while that of *N. grandis* further decreases in parts of Indo-Burman Hills but remains roughly the same in the Eastern Himalaya.

**Figure 3:**
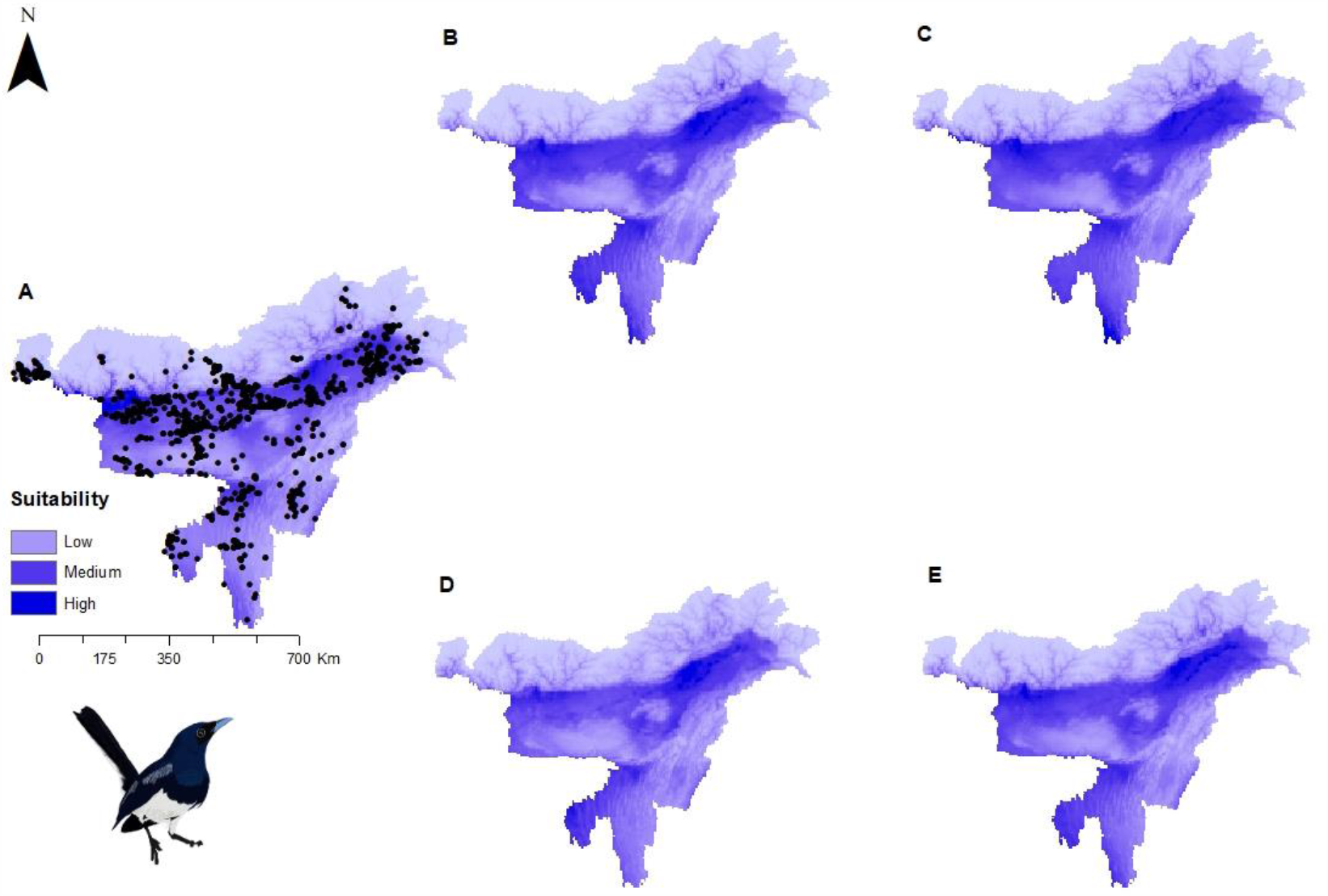
Potential distributions of *Copsychus saularis* (Oriental Magpie-Robin), a habitat generalist, under present climate with presence points (A) and under four future climatic scenarios- (B) SSP1-2.6 (low), (C) SSP2-4.5 (intermediate), (D) SSP 4-6.0 (medium) and (E) SSP5-8.5 (high)

**Figure 4:**
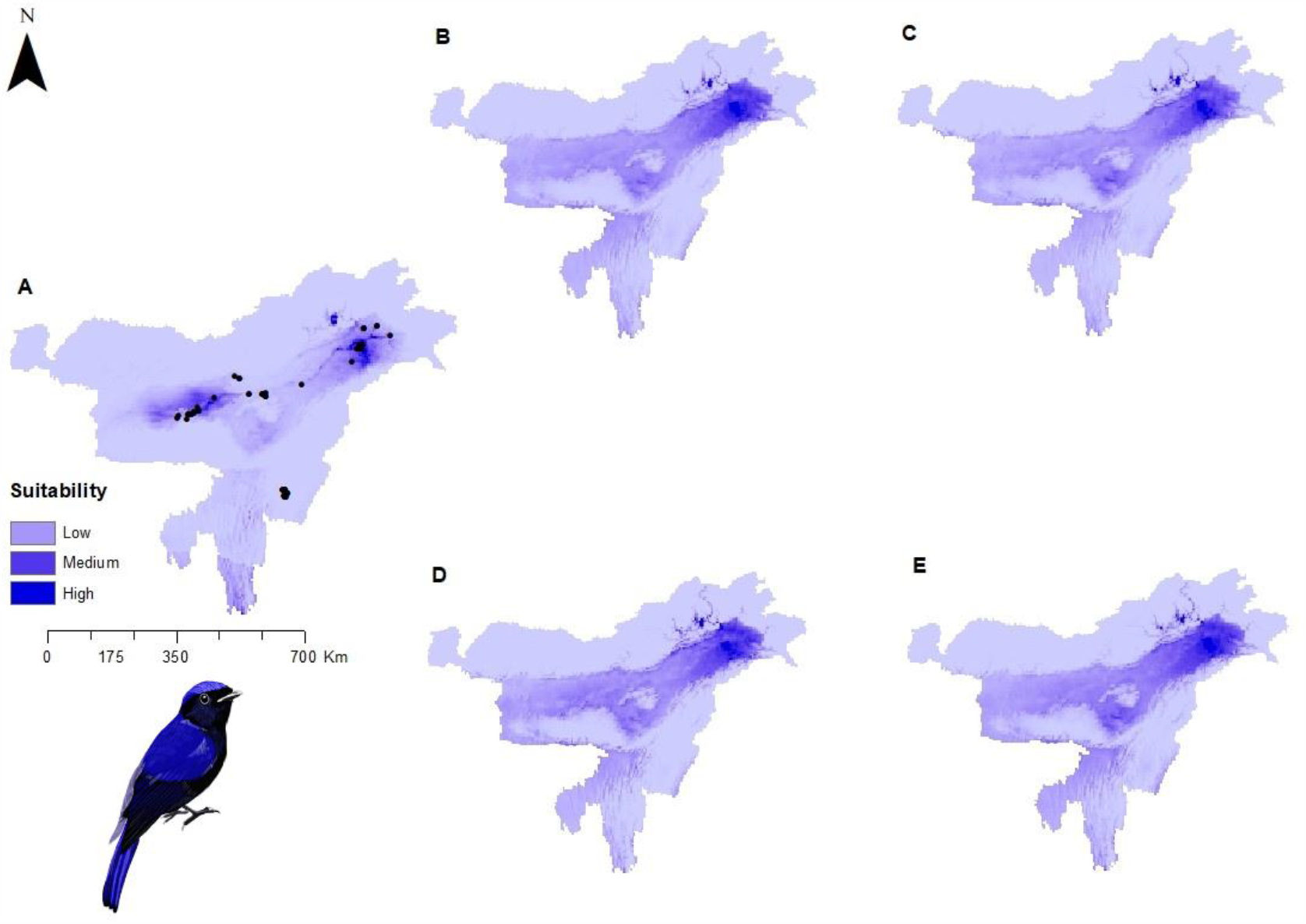
Potential distributions of *Niltava grandis* (Large Niltava), a habitat specialist, under present climate with presence points (A) and under four future climatic scenarios- (B) SSP1-2.6 (low), (C) SSP2-4.5 (intermediate), (D) SSP 4-6.0 (medium) and (E) SSP5-8.5 (high)

For the other 58 species (see Supplementary Figures S1 to S58), Maxent predicted restricted potential distributions for *Phoenicurus erythrogastrus* (White-winged Redstart) (Figure S43), *Brachypteryx hyperythra* (Rusty-bellied Shortwing) (Figure S3), *Monticola cinclorhyncha* (Blue-capped Rock Thrush) (Figure S31), and *P. schisticeps* (White-throated Redstart) (Figure S49), and wider potential distributions for *Saxicola maurus* (Siberian Stonechat) (Figure S54), *P. auroreus* (Daurian Redstart) (Figure S42), *Ficedula parva* (Red-breasted Flycatcher) (Figure S20), and *Muscicapa muttui* (Brown-breasted Flycatcher) (Figure S36).

### Phylogenetic analysis

*S’*, a measure of habitat specialism, was negatively correlated with percentage contribution of bio8 under the present climate, and with the percentage contribution of tmax under low, intermediate, medium, and high emission scenarios (Table 2). However, there was no correlation between *S’* and elevation under present climate.

**Table 2:** Results from Phylogenetic Generalised Least Squares (PGLS) models using Percentage Contribution of Variables (PCVs) for present and future climatic conditions (*P-values* of significant models are in **bold**)

| PCVs | Value | Standard error | t-value | P-values |
| --- | --- | --- | --- | --- |
| **Present climate** |  |  |  |  |
| bio8 | -27.36 | 13.52 | -2.02 | **<0.05** |
| elevation | 10.15 | 8.49 | 1.19 | 0.23 |
| **Future climate** |  |  |  |  |
| SSP1-2.6 (low emission scenario) |  |  |  |  |
| tmax | -37.26 | 013.62 | -2.73 | **<0.01** |
| SSP2-4.5 (intermediate emission scenario) |  |  |  |  |
| tmax | -32.22 | 12.44 | -2.58 | **<0.05** |
| SSP4-6.0 (medium emission scenario) |  |  |  |  |
| tmax | -41.57 | 12.52 | -3.31 | **<0.01** |
| SSP5-8.5 (high emission scenario) |  |  |  |  |
| tmax | -2.05 | 0.57 | -3.55 | **<0.001** |

Moran’s *I* dot plot demonstrated presence of autocorrelation amongst variables with the highest percentage contributions and tips of the phylogenetic tree representing species. Although there was positive correlation in contribution of bio8 amongst *Niltava macgrigoriae, N. grandis, Myophonus caeruleus, Myiomela leucura, Muscicapa sibirica*, and amongst *Muscicapa dauurica, Monticola solitarius, Monticol rufiventris* (Figure 5), the phylogenetic signal had overall low strength (*K*= 0.40) generally implying that closely related species of Muscicapidae Flycatchers did not have an overlap in contributions of bio8 to their niche models.

**Figure 5:**
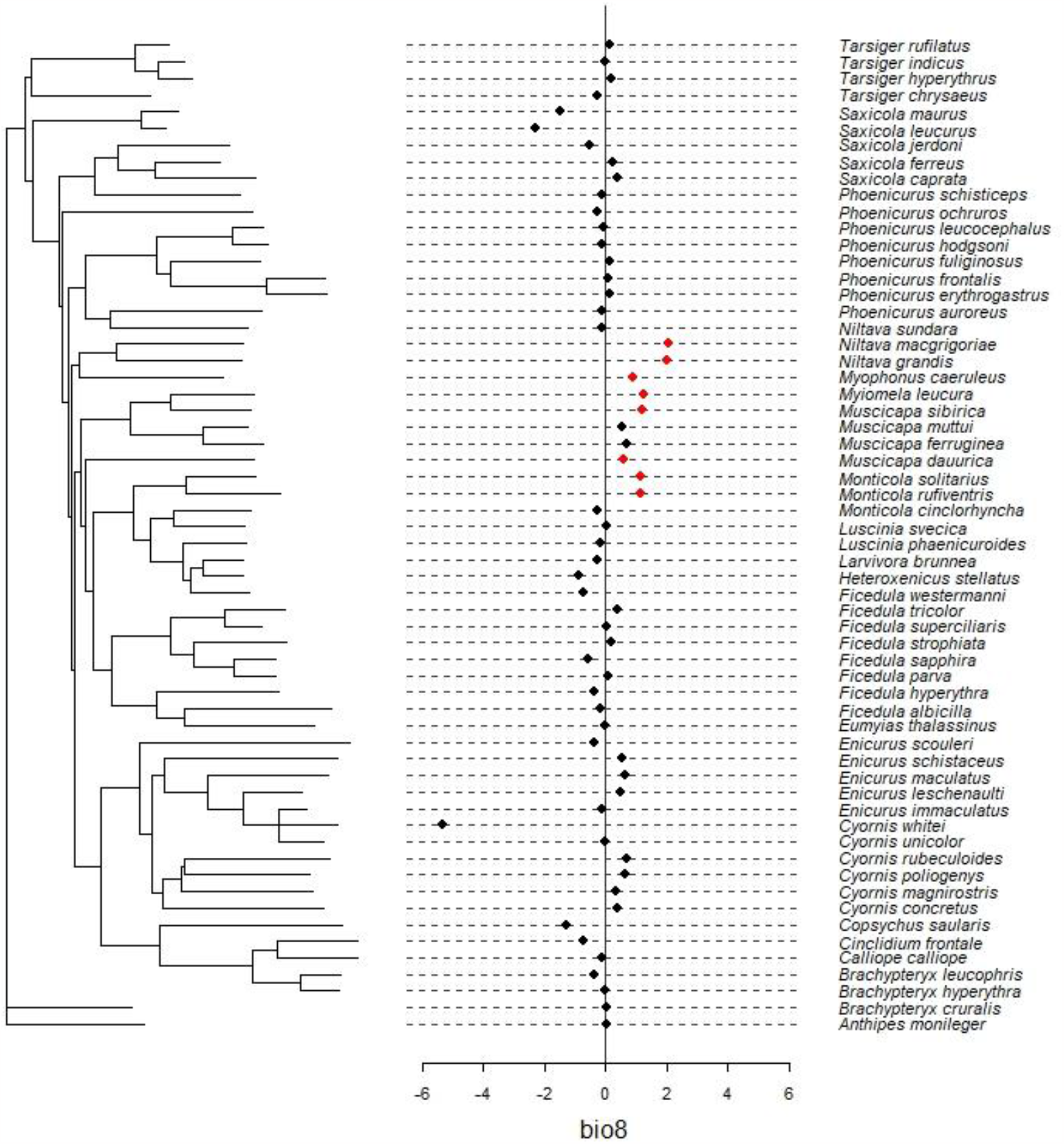
Local Moran’s *I* plot of phylogenetic signal in percentage contribution of bio8. The red dots falling on and side of zero line correspond to significant relationships indicating positive correlation

Under future climatic conditions, positive correlation continues in contribution of tmax amongst species such as *N. sundara, N. macgrigoriae, M. caeruleus*, and amongst *M. dauurica, M. rufiventris, M. solitaries* (see Supplementary Figure S63). However, there was low strength in phylogenetic signal in response to tmax under low emission scenario (*K*=0.41), intermediate emission scenario (*K*=0.46), medium emission scenario (*K*=0.41), and high emission scenario (*K*=0.34).

## DISCUSSION

The region of Northeast India and Bhutan is highly susceptible to climate change (Bajracharya et al., 2007; Ravindranath et al., 2011), but its effects on species inhabiting the region are largely understudied. The predicted distributions effects of climate change on the Muscicapidae Flycatchers, one of the largest terrestrial bird families in this region, showed varied responses across the family with little evidence of phylogenetic niche conservatism in species’ sensitivity to abiotic variables under all climate scenarios. Rather, habitat specialism explained sensitivity to bioclimatic variables under both present and future climate scenarios. However, in contrast to expectations, we found evidence that the distributions of habitat generalists rather than specialists, were most responsive to bioclimatic variables.

At a broad level, the variable contribution to potential distributions followed our general expectations under present climatic conditions. Most birds had high contributions from the mean temperature of the wettest quarter (bio8), implying that the potential distributions, and thus the abiotic niches of our species, are impacted by a combination of temperature and precipitation conditions. Bio8 has previously shown high contributions to potential distributions of many Himalayan and Indo-Burman species such as the medicinal plants of Liliaceae family (Rana et al., 2017), the native pheasant-Himalayan Monal *Lophophorus impejanus* (Rai et al., 2020), the invasive *Lantana camara L*. (Bushi et al., 2022), the threatened orchid *Satyrium nepalense* (Kumar and Rawat, 2022), the Crayfish *Cherax quadricarinatus* and *Procambarus virginalis* (Zeng and Yeo, 2018), and the falcon *Falco jugger* (Sutton et al., 2020). Thus, several species of this region may be expected to track their thermal as well as hygric niches, which may lead to complex patterns of directions of range shifts or distribution changes in the future. We expected the trend to continue, i.e. high contributions from temperature and precipitation variables to the ENMs under future climatic scenarios as well, but we found the maximum temperature (tmax) had the highest contribution while precipitation (prec) had the lowest. The lower importance of precipitation to potential distributions of species under future climate scenario models is plausibly due to the large uncertainties in precipitation projections in the CMIP6 General Circulation Models (John et al., 2022), causing in an underestimation of contributions of the precipitation variable.

The second important finding is that the sensitivity to abiotic components bio8 under present climate, and tmax under future climate was explained by species specialisation index in PGLS models, and contrary to our expectation, we found that as habitat specialism increased, species’ sensitivity to abiotic variable declined. However, highly habitat specialist species such as *Brachypteryx cruralis* (Himalayan Shortwing), *E. leschenaulti* (White-crowned Forktail), and *Muscicapa muttui* (Brown-breasted Flycatcher) had sparse presence points in fewer grids to begin with which might have been insufficient in predicting precise degrees of sensitivity to abiotic variables across a large spatial scale. We expect such patterns could be because of uncertainties associated with gridded climatic data over the region, extrapolations from which may lead to added challenges for conservation planning and management under climate change.

The third major finding that is we did not find evidence of phylogenetic niche conservatism in the contributions of abiotic variables to potential distributions under climate change. Closely-related species are expected to have overlapping ecological niches (Losos, 2008) and therefore may have similar responses to climate change. A global assessment of thermal traits reported a strong signal of phylogenetic niche conservatism in pantropical birds and mammal species at a large geographical scale (Khaliq et al., 2015). Phylogenetic niche conservatism in closely-related species has also been reported in responses to climate change in regional examples such as diversity-climate relationships in Rosaceae species in China (Wang et al., 2021), and overlapping prediction ranges for some of the western *Plethodon* species in the Pacific Northwest (Nottingham and Pelletier, 2021). However, there are also empirical examples to the contrary-at regional and local scales, for instance, *Prionailurus* species show disparate spatial overlap resulting in heterogeneous responses to environmental responses in the Indian subcontinent (Silva et al., 2020), and closely-related Atlantic corals exhibiting random patterns of associations between functional traits and distributional shifts (Rodriguez et al., 2019). As more empirical examples accrue, we will gain a better understanding of when phylogenetic niche conservatism in response to climate change may be important, however we should not assume that phylogeny is predictive along all axes of responses.

### Caveats

While all our Maxent models performed well, it is important to highlight the role of spatial scales and data required to develop ecological niche models. We expect the results could vary for the Muscicapidae Flycatchers if undertaken over a larger scale or smaller scale which will lead to an increase or decrease in the number of habitat types available for species to occupy. Since most of the tropical regions are largely data deficient (Butchart and Bird, 2010), we were limited by data availability for these species in our region and hence proceeded with presence-only models. It is likely that the availability of information on dispersal ability and biotic interactions will improve findings (e.g., Heikkinen et al., 2007). However, it is also likely that even with additional data on biotic interactions, the predictions may continue to remain the same as these effects are difficult to capture at macro-scales (Araújo and Luoto, 2007). Lastly, the niche models we used are static and do not take into consideration the mechanisms of evolutionary adaptations which cannot be ruled out for complex mountainous environments offering opportunities for high speciation.

## Acknowledgments

Aavika Dhanda was supported by the Oxford Mary-de Zouche Graduate Scholarship (Oxford India Centre for Sustainable Development at Somerville College, University of Oxford). The authors would like to thank Shivangi Chowdhry for digitally illustrating 60 bird species as seen in figures and maps in the main text and supplementary file.

## Author contributions

Aavika Dhanda conceptualized the idea, designed the methodology, performed modeling and data analysis, and prepared the original draft. Michał T. Jezierski performed modeling and model interpretation. Tim Coulson contributed to methodology design and model interpretation. Sonya Clegg contributed to conceptualizing the idea and model interpretation. All authors contributed to giving critical feedback to drafts.

## Data and code availability

R code and the related files are available at https://github.com/aavikad/muscicapidae-flycatchers-_v1.0

