## Supplementary Figures S1-S63 for "Ecology and not phylogeny influences sensitivity to climate change in Muscicapidae Flycatchers in Eastern Himalayan and Indo-Burman hotspots"

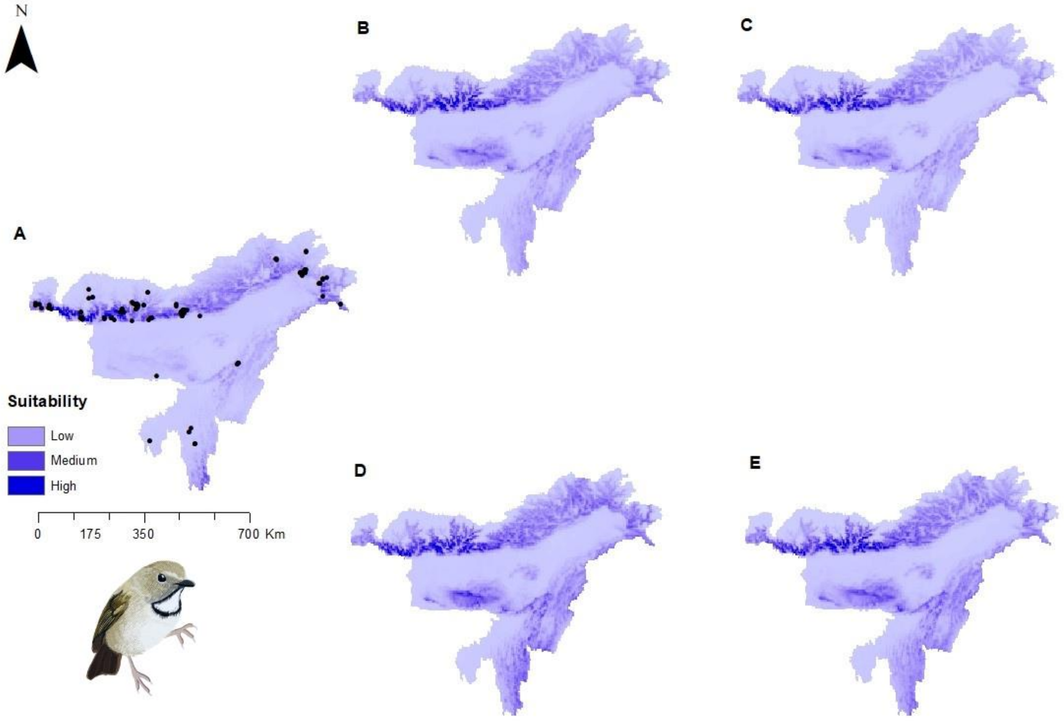


Figure S1: Potential distributions of *Anthipes monileger* (White-gorgeted Flycatcher) under present climate with presence points (A) and under four future climatic scenarios- (B) SSP1-2.6 (low), (C) SSP2-4.5 (intermediate), (D) SSP 4-6.0 (medium) and (E) SSP5-8.5 (high)

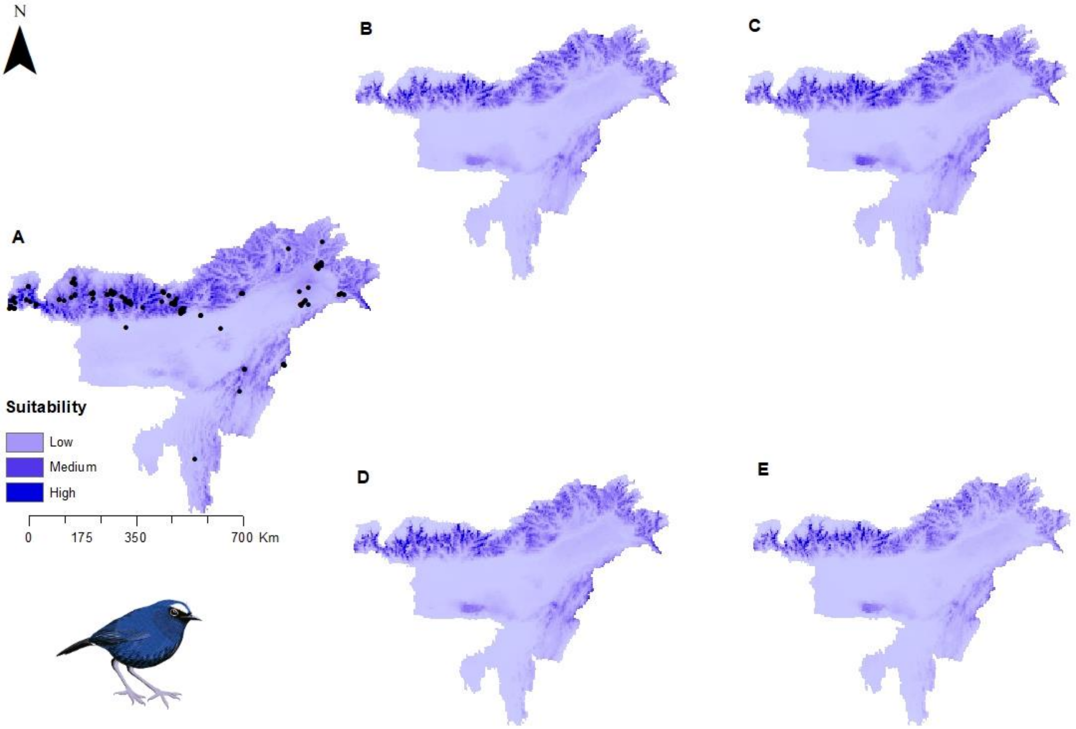


Figure S2: Potential distributions of *Brachypteryx cruralis* (Himalayan Shortwing) under present climate with presence points (A) and under four future climatic scenarios- (B) SSP1-2.6 (low), (C) SSP2-4.5 (intermediate), (D) SSP 4-6.0 (medium) and (E) SSP5-8.5 (high)

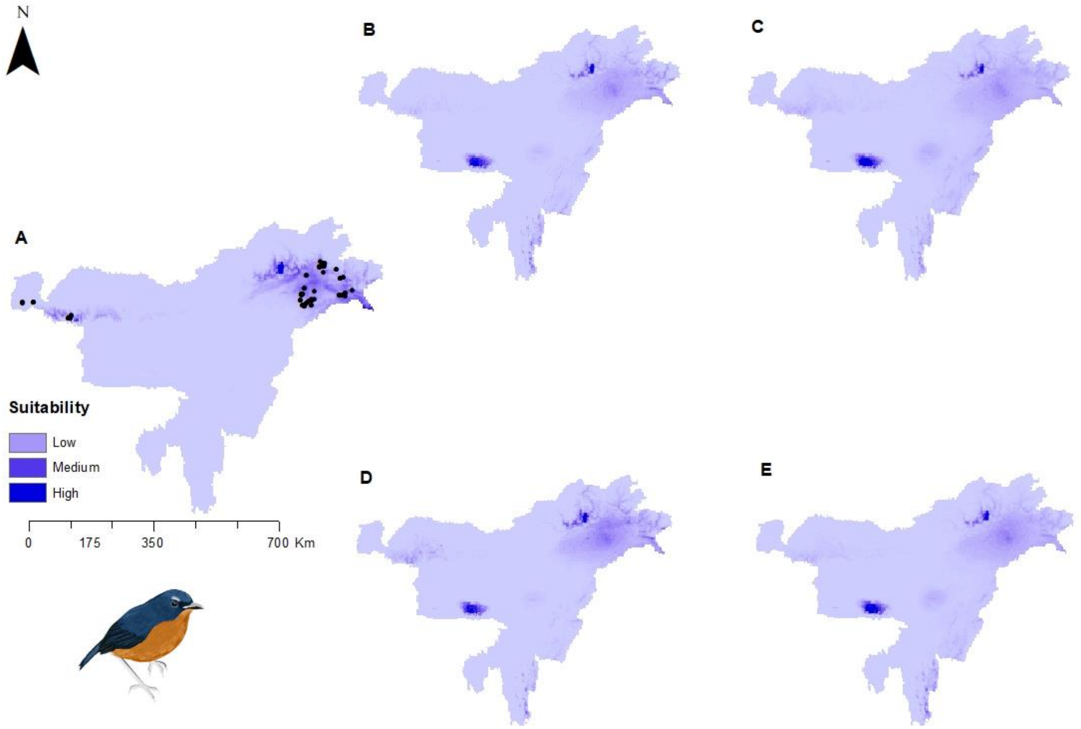


Figure S3: Potential distributions of *Brachypteryx hyperythra* (Rusty-bellied Shortwing) under present climate with presence points (A) and under four future climatic scenarios- (B) SSP1-2.6 (low), (C) SSP2-4.5 (intermediate), (D) SSP 4-6.0 (medium) and (E) SSP5-8.5 (high)

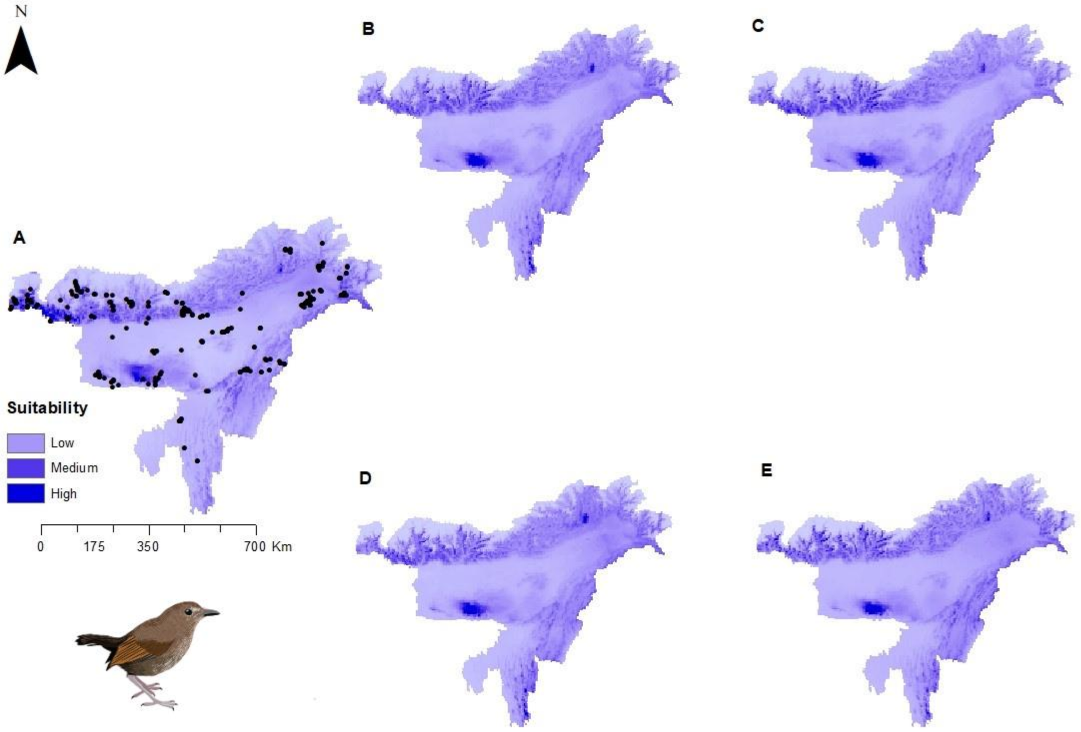


Figure S4: Potential distributions of *Brachypteryx leucophris* (Lesser Shortwing) under present climate with presence points (A) and under four future climatic scenarios- (B) SSP1-2.6 (low), (C) SSP2-4.5 (intermediate), (D) SSP 4-6.0 (medium) and (E) SSP5-8.5 (high)

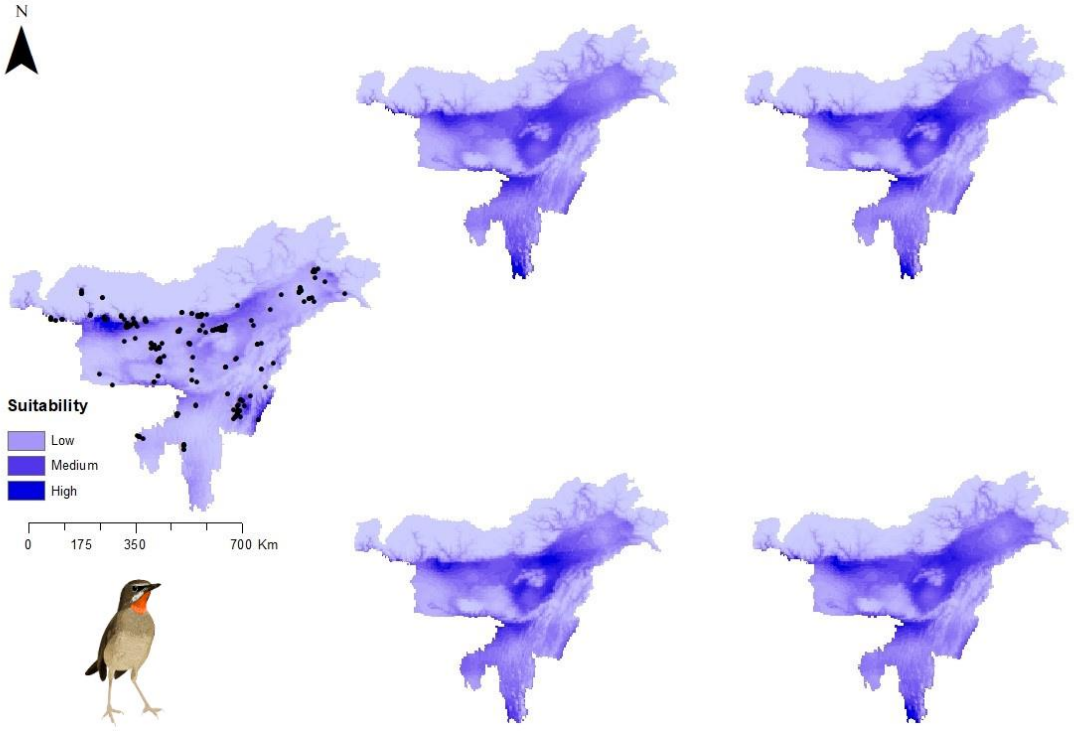


Figure S5: Potential distributions of *Calliope calliope* (Siberian Rubythroat ) under present climate with presence points (A) and under four future climatic scenarios- (B) SSP1-2.6 (low), (C) SSP2-4.5 (intermediate), (D) SSP 4-6.0 (medium) and (E) SSP5-8.5 (high)

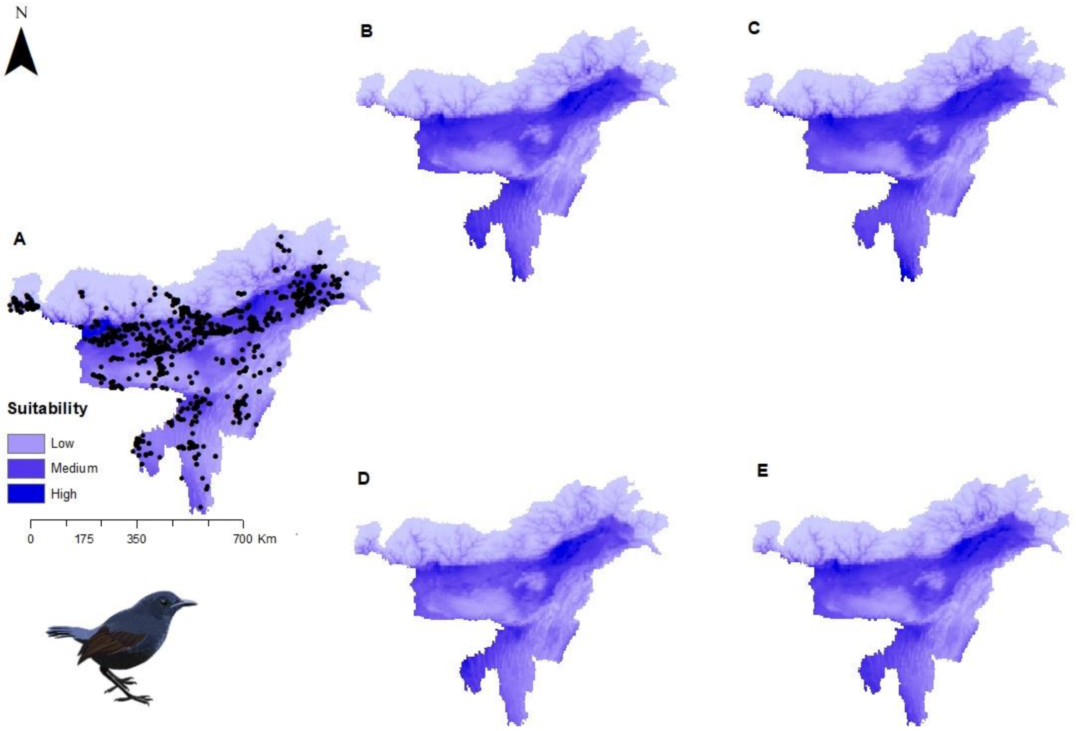


Figure S6: Potential distributions of *Cinclidium frontale* (Blue-fronted Robin) under present climate with presence points (A) and under four future climatic scenarios- (B) SSP1-2.6 (low), (C) SSP2-4.5 (intermediate), (D) SSP 4-6.0 (medium) and (E) SSP5-8.5 (high)

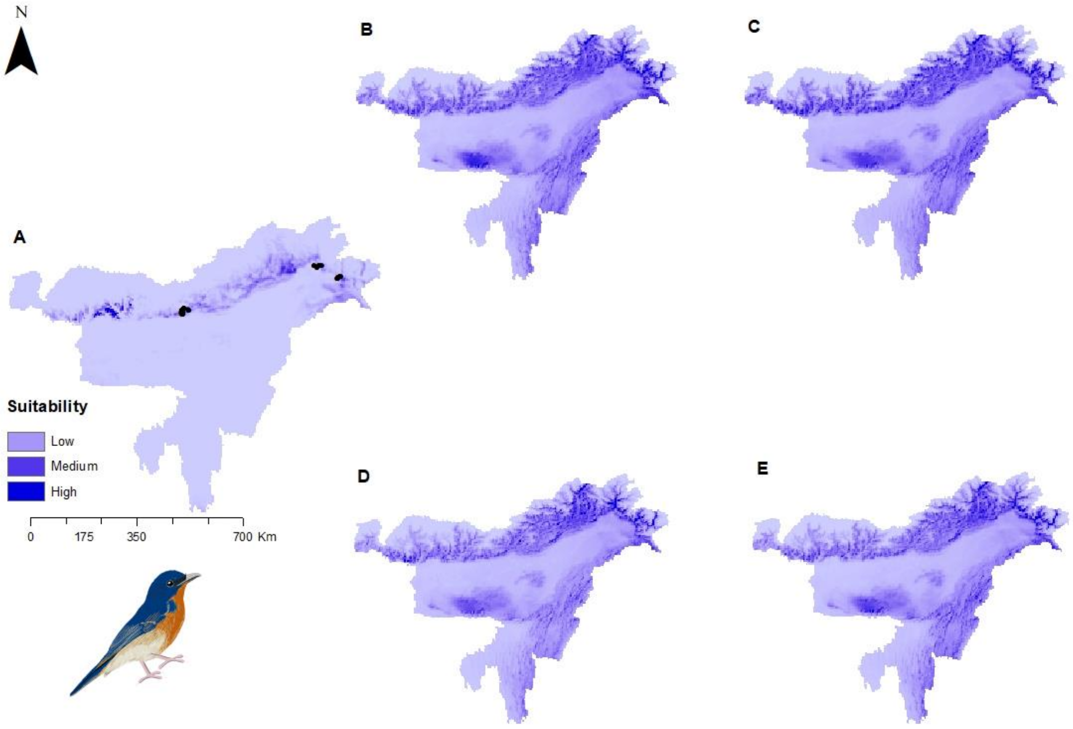


Figure S7: Potential distributions of *Cyornis magnirostris* (Large Blue Flycatcher) under present climate with presence points (A) and under four future climatic scenarios- (B) SSP1-2.6 (low), (C) SSP2-4.5 (intermediate), (D) SSP 4-6.0 (medium) and (E) SSP5-8.5 (high)

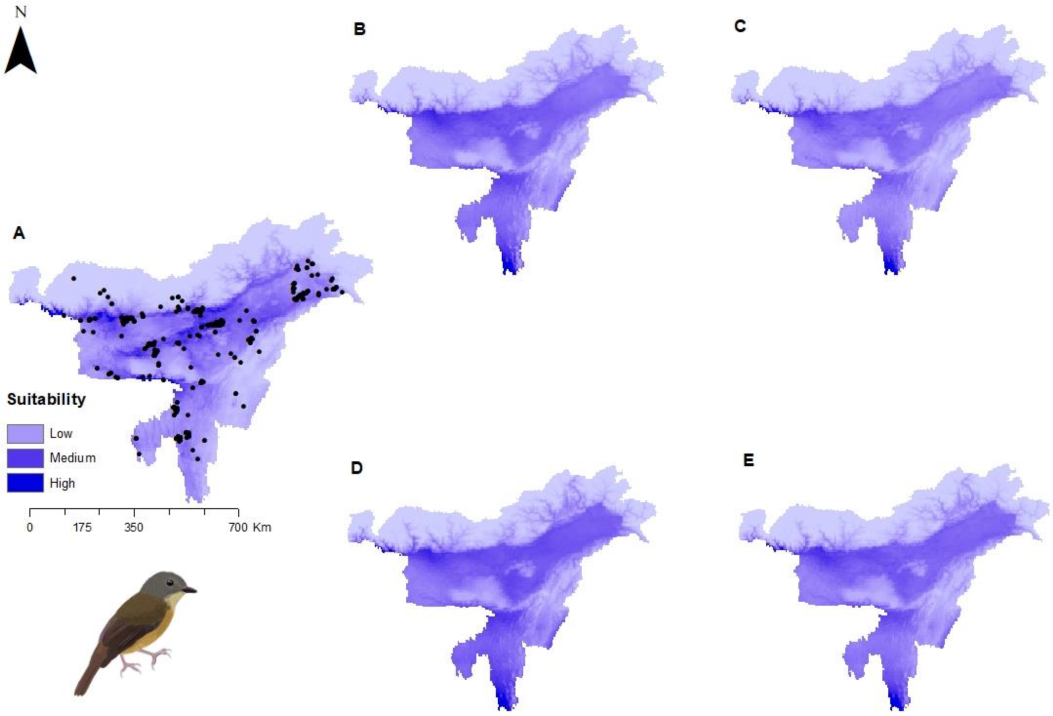


Figure S8: Potential distributions of *Cyornis poliogenys* (Pale-chinned Blue Flycatcher) under present climate with presence points (A) and under four future climatic scenarios- (B) SSP1-2.6 (low), (C) SSP2-4.5 (intermediate), (D) SSP 4-6.0 (medium) and (E) SSP5-8.5 (high)

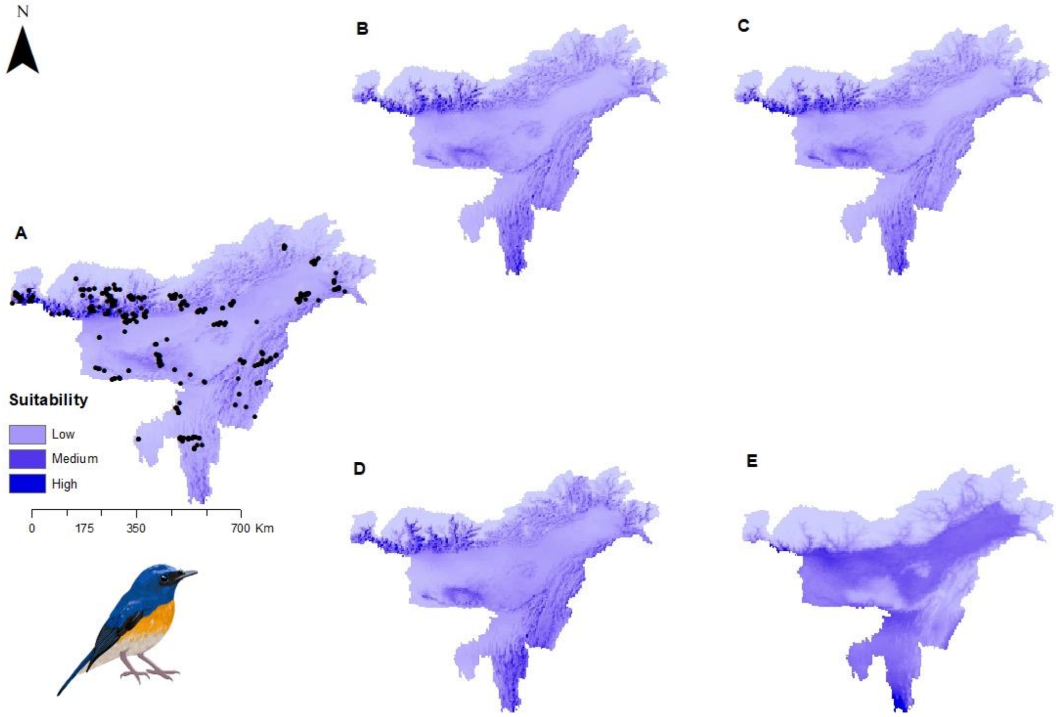


Figure S9: Potential distributions of *Cyornis rubeculoides* (Blue-throated Flycatcher) under present climate with presence points (A) and under four future climatic scenarios- (B) SSP1-2.6 (low), (C) SSP2-4.5 (intermediate), (D) SSP 4-6.0 (medium) and (E) SSP5-8.5 (high)

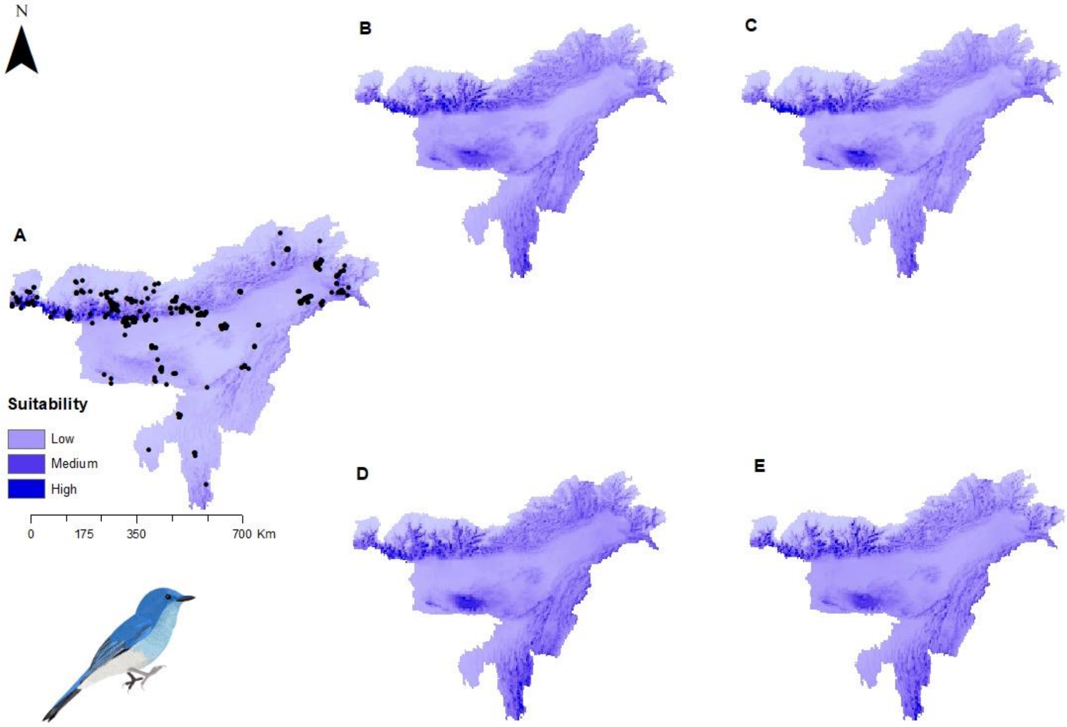


Figure S10: Potential distributions of *Cyornis unicolor* (Pale Blue Flycatcher) under present climate with presence points (A) and under four future climatic scenarios- (B) SSP1-2.6 (low), (C) SSP2-4.5 (intermediate), (D) SSP 4-6.0 (medium) and (E) SSP5-8.5 (high)

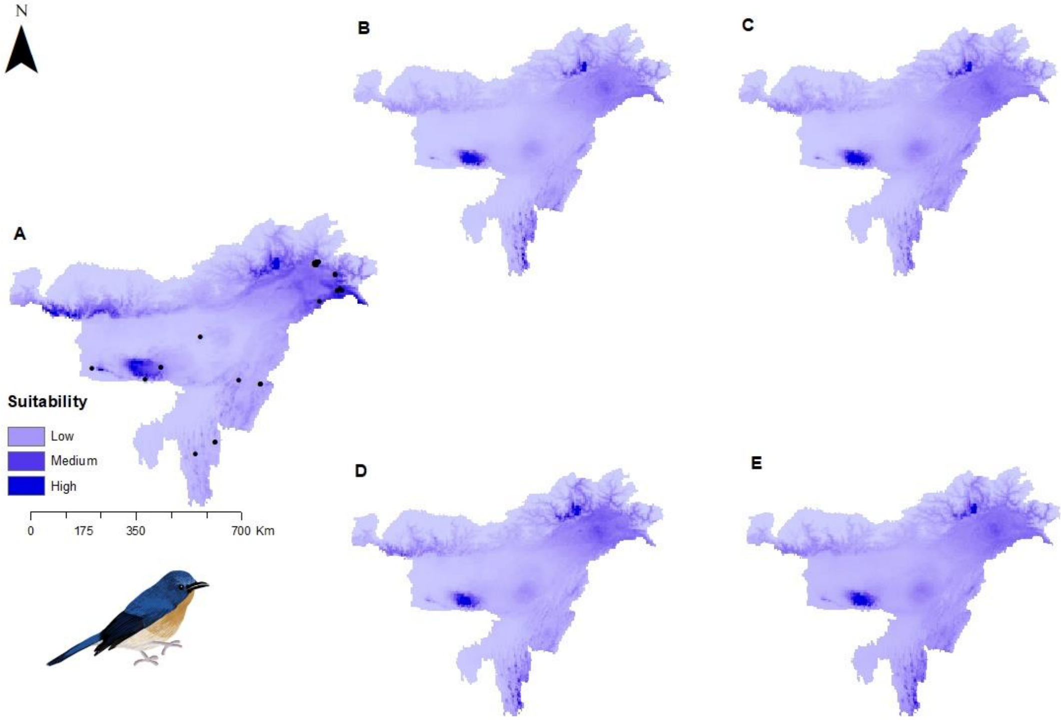


Figure S11: Potential distributions of *Cyornis whitei* (Hill Blue Flycatcher) under present climate with presence points (A) and under four future climatic scenarios- (B) SSP1-2.6 (low), (C) SSP2-4.5 (intermediate), (D) SSP 4-6.0 (medium) and (E) SSP5-8.5 (high)

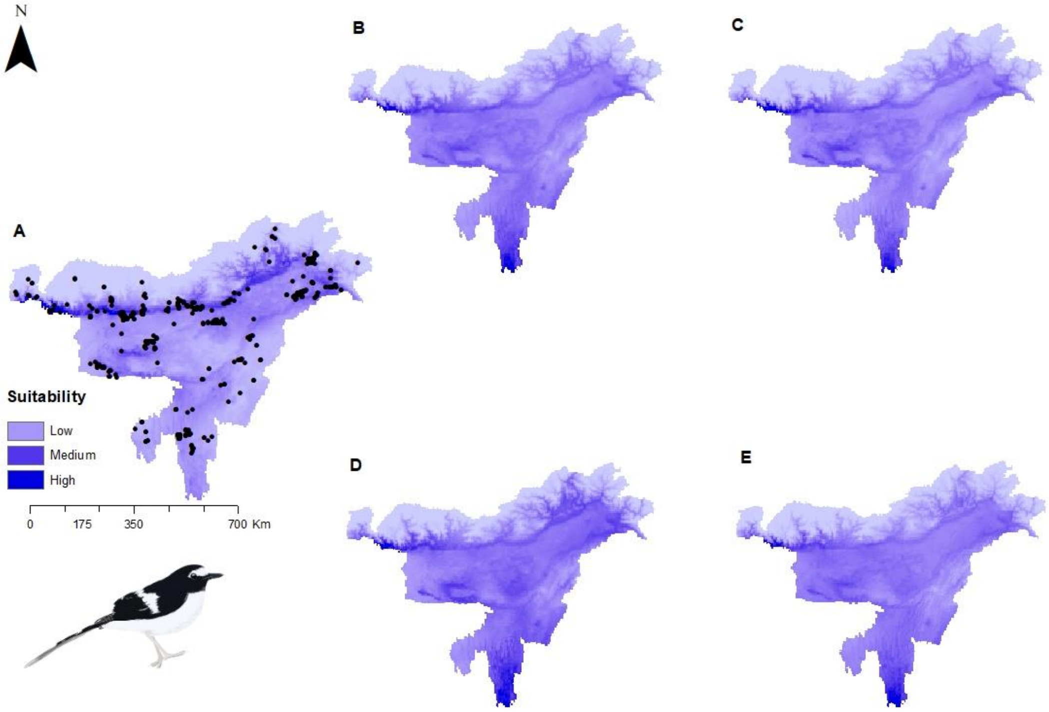


Figure S12: Potential distributions of *Enicurus immaculatus* (Black-backed Forktail) under present climate with presence points (A) and under four future climatic scenarios- (B) SSP1-2.6 (low), (C) SSP2-4.5 (intermediate), (D) SSP 4-6.0 (medium) and (E) SSP5-8.5 (high)

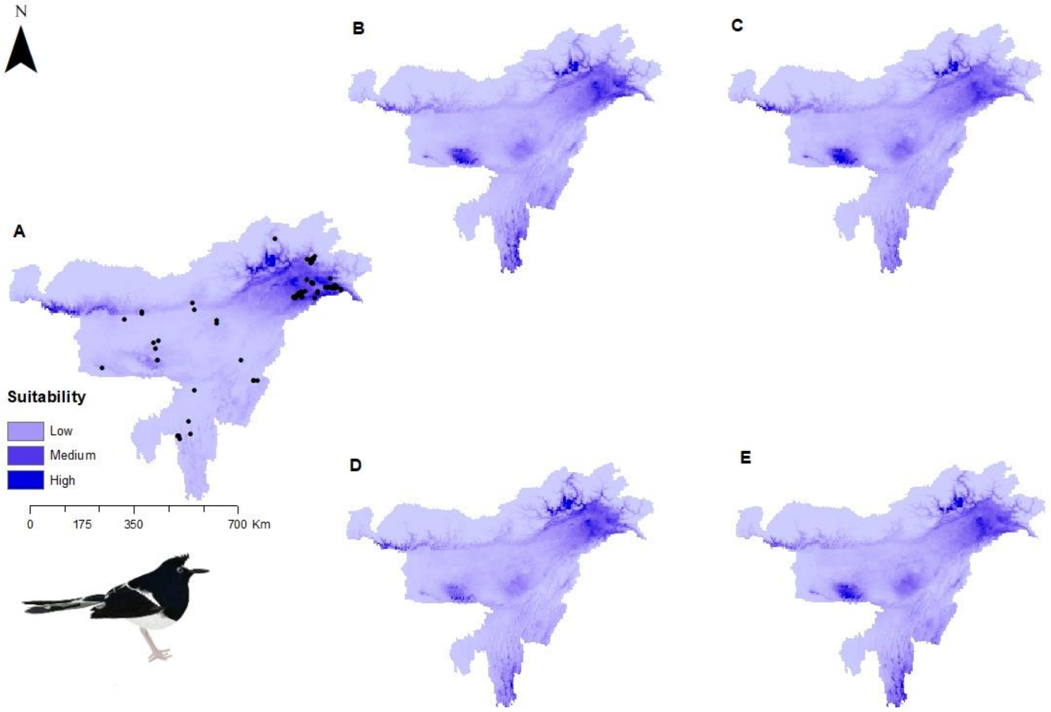


Figure S13: Potential distributions of *Enicurus leschenaulti* (White-crowned Forktail) under present climate with presence points (A) and under four future climatic scenarios- (B) SSP1-2.6 (low), (C) SSP2-4.5 (intermediate), (D) SSP 4-6.0 (medium) and (E) SSP5-8.5 (high)

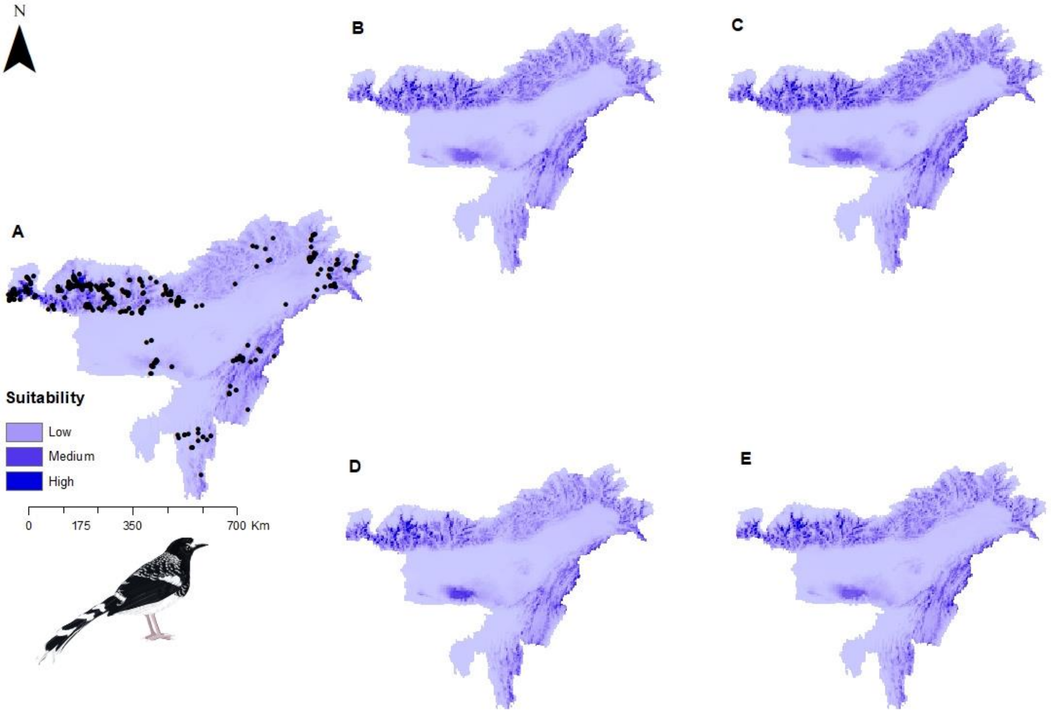


Figure S14: Potential distributions of *Enicurus maculatus* (Spotted Forktail) under present climate with presence points (A) and under four future climatic scenarios- (B) SSP1-2.6 (low), (C) SSP2-4.5 (intermediate), (D) SSP 4-6.0 (medium) and (E) SSP5-8.5 (high)

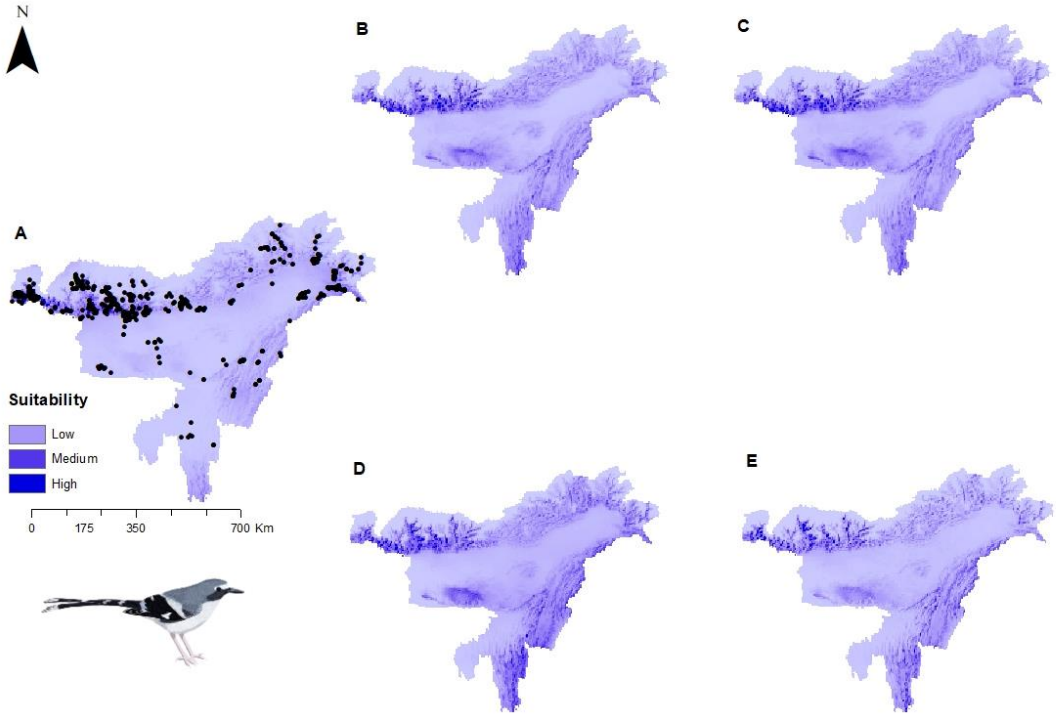


Figure S15: Potential distributions of *Enicurus schistaceus* (Slaty-backed Forktail) under present climate with presence points (A) and under four future climatic scenarios- (B) SSP1-2.6 (low), (C) SSP2-4.5 (intermediate), (D) SSP 4-6.0 (medium) and (E) SSP5-8.5 (high)

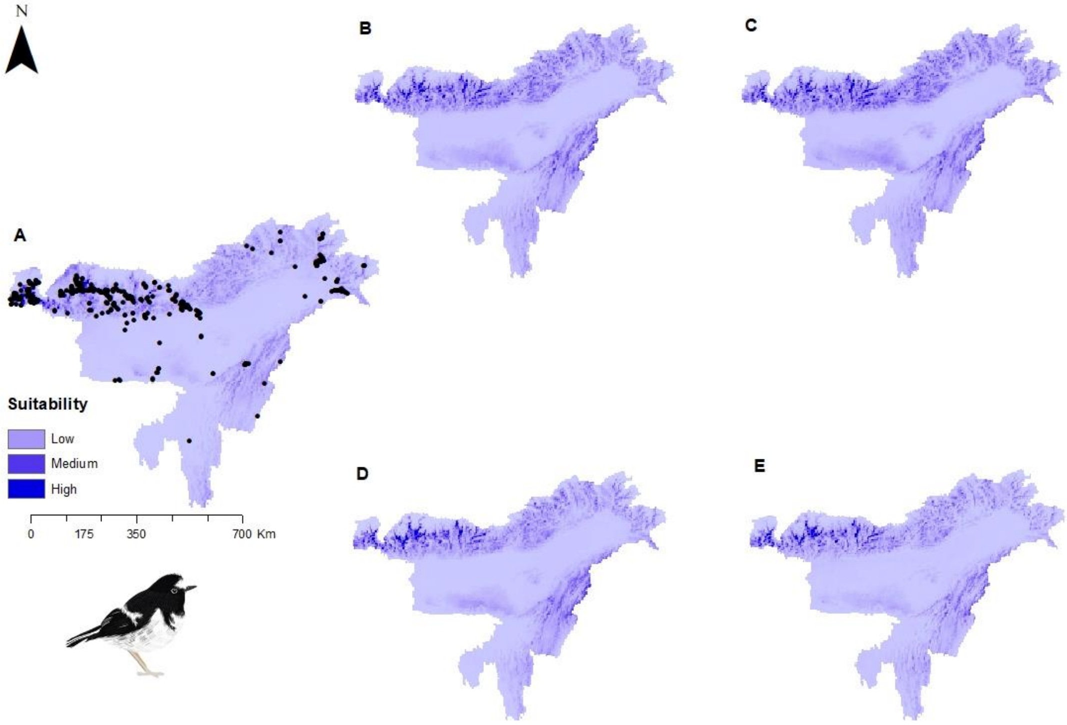


Figure S16: Potential distributions of *Enicurus scouleri* (Little Forktail) under present climate with presence points (A) and under four future climatic scenarios- (B) SSP1-2.6 (low), (C) SSP2-4.5 (intermediate), (D) SSP 4-6.0 (medium) and (E) SSP5-8.5 (high)

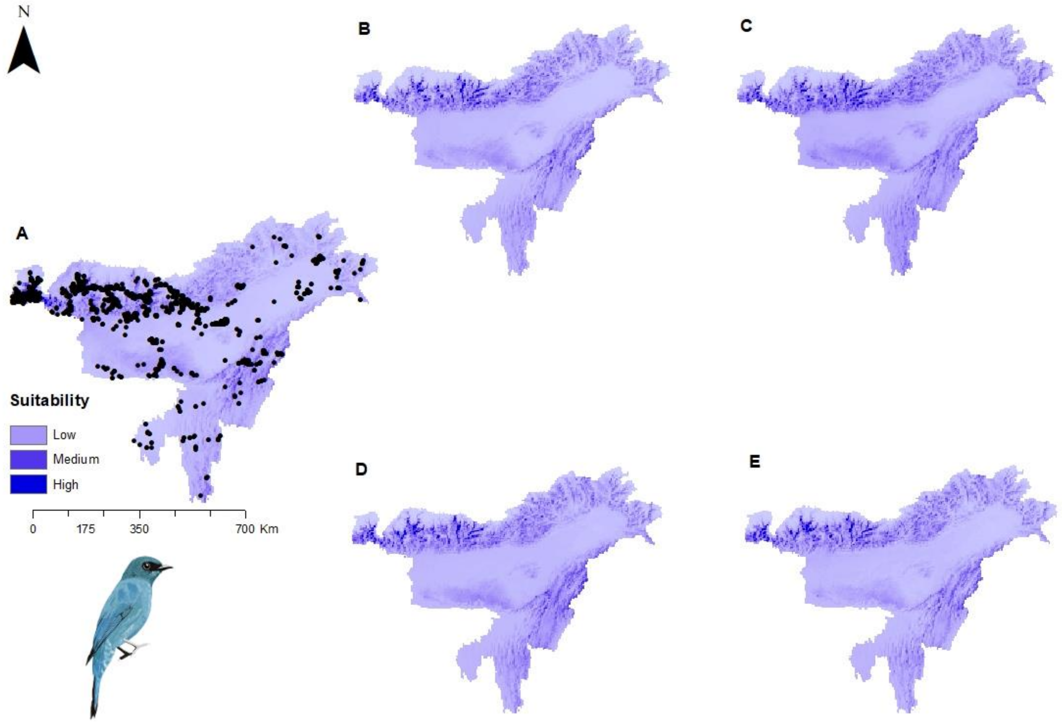


Figure S17: Potential distributions of *Eumyias thalassinus* (Verditer Flycatcher) under present climate with presence points (A) and under four future climatic scenarios- (B) SSP1-2.6 (low), (C) SSP2-4.5 (intermediate), (D) SSP 4-6.0 (medium) and (E) SSP5-8.5 (high)

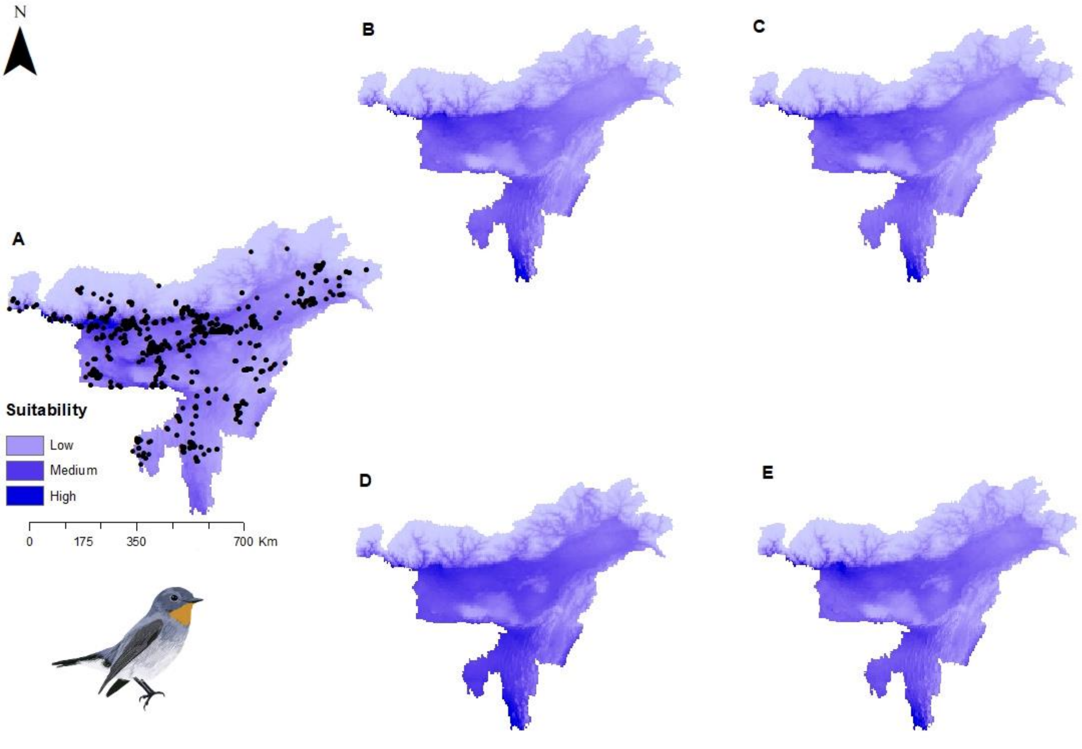


Figure S18: Potential distributions of *Ficedula albicilla* (Taiga Flycatcher) under present climate with presence points (A) and under four future climatic scenarios- (B) SSP1-2.6 (low), (C) SSP2-4.5 (intermediate), (D) SSP 4-6.0 (medium) and (E) SSP5-8.5 (high)

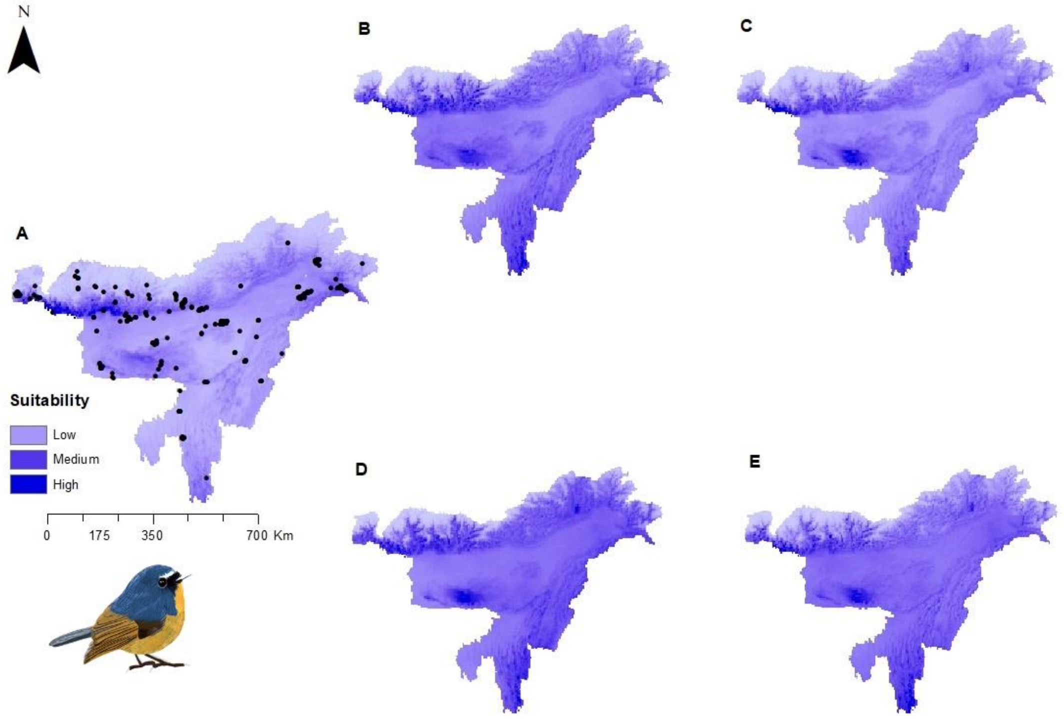


Figure S19: Potential distributions of *Ficedula hyperythra* (Snowy-browed Flycatcher) under present climate with presence points (A) and under four future climatic scenarios- (B) SSP1-2.6 (low), (C) SSP2-4.5 (intermediate), (D) SSP 4-6.0 (medium) and (E) SSP5-8.5 (high)

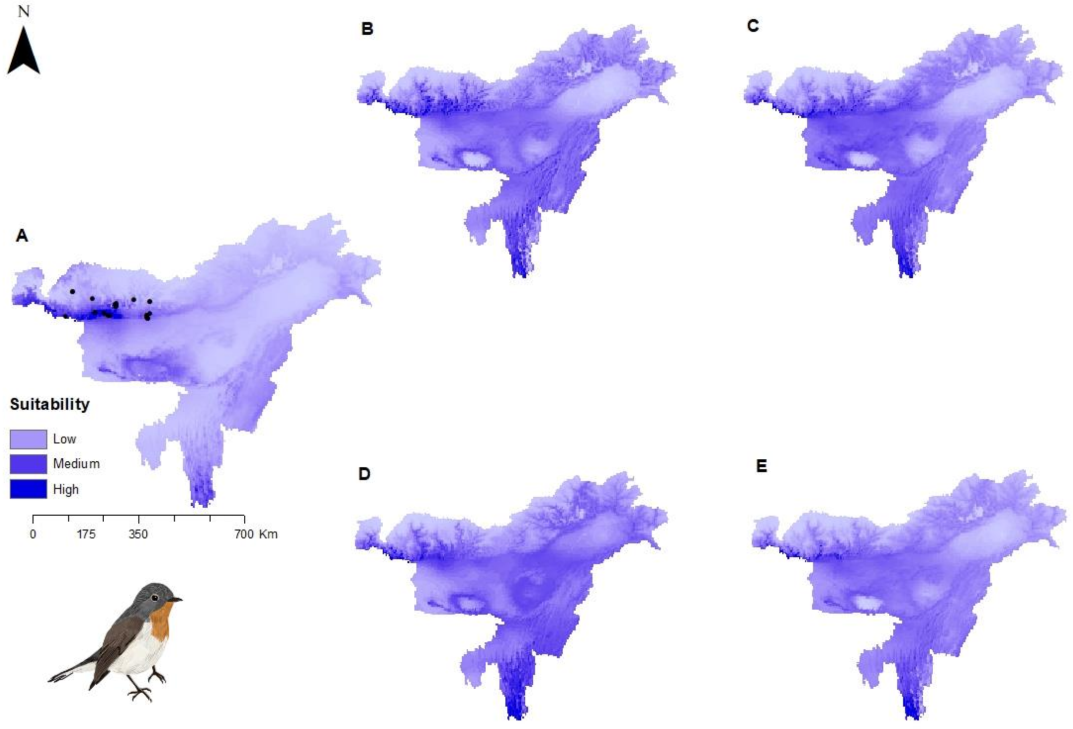


Figure S20: Potential distributions of *Ficedula parva* (Red-breasted Flycatcher) under present climate with presence points (A) and under four future climatic scenarios- (B) SSP1-2.6 (low), (C) SSP2-4.5 (intermediate), (D) SSP 4-6.0 (medium) and (E) SSP5-8.5 (high)

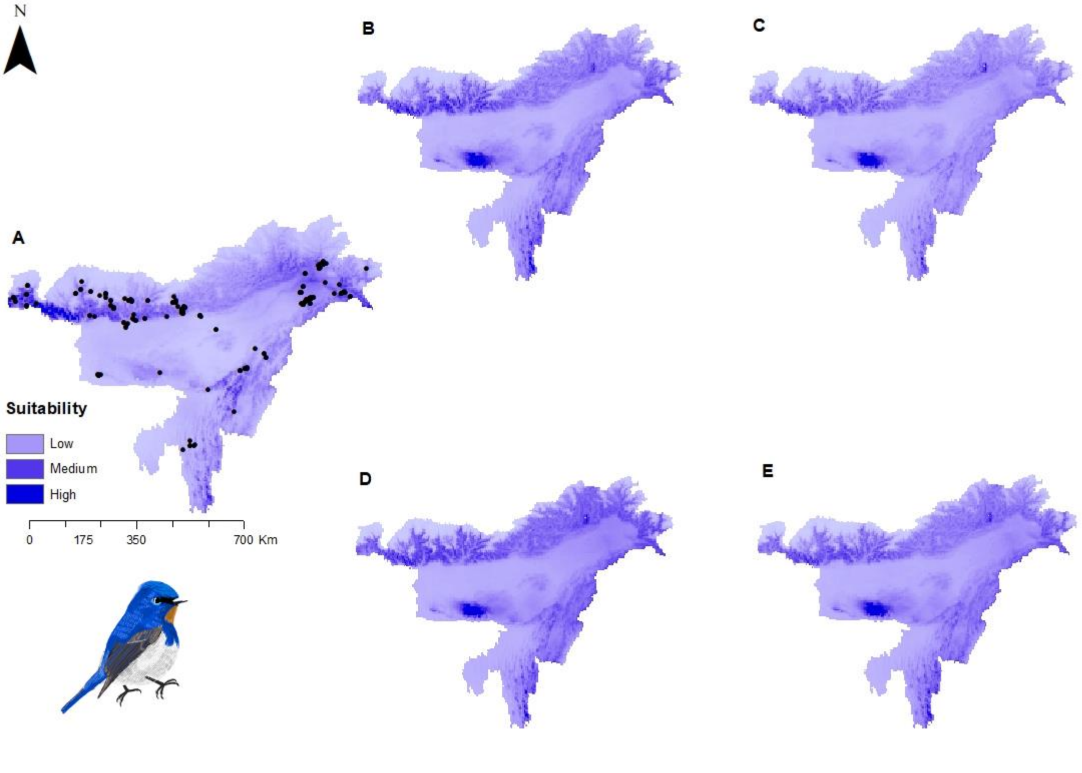


Figure S21: Potential distributions of *Ficedula sapphira* (Sapphire Flycatcher) under present climate with presence points (A) and under four future climatic scenarios- (B) SSP1-2.6 (low), (C) SSP2-4.5 (intermediate), (D) SSP 4-6.0 (medium) and (E) SSP5-8.5 (high)

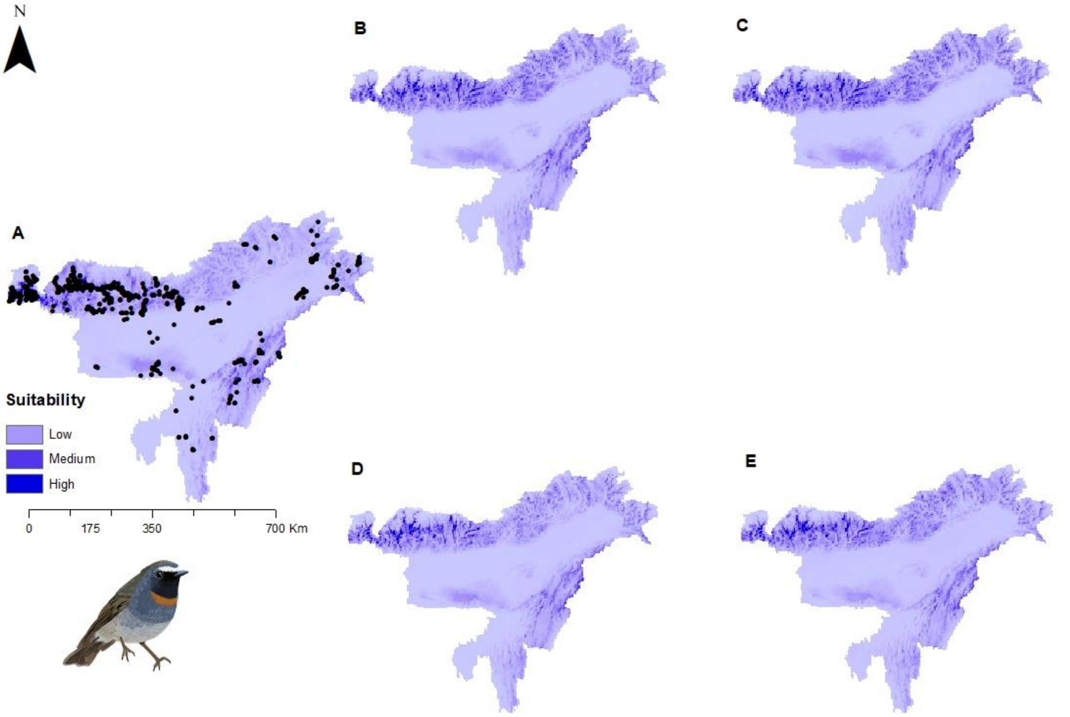


Figure S22: Potential distributions of *Ficedula strophiata* (Rufous-gorgeted Flycatcher) under present climate with presence points (A) and under four future climatic scenarios- (B) SSP1-2.6 (low), (C) SSP2-4.5 (intermediate), (D) SSP 4-6.0 (medium) and (E) SSP5-8.5 (high)

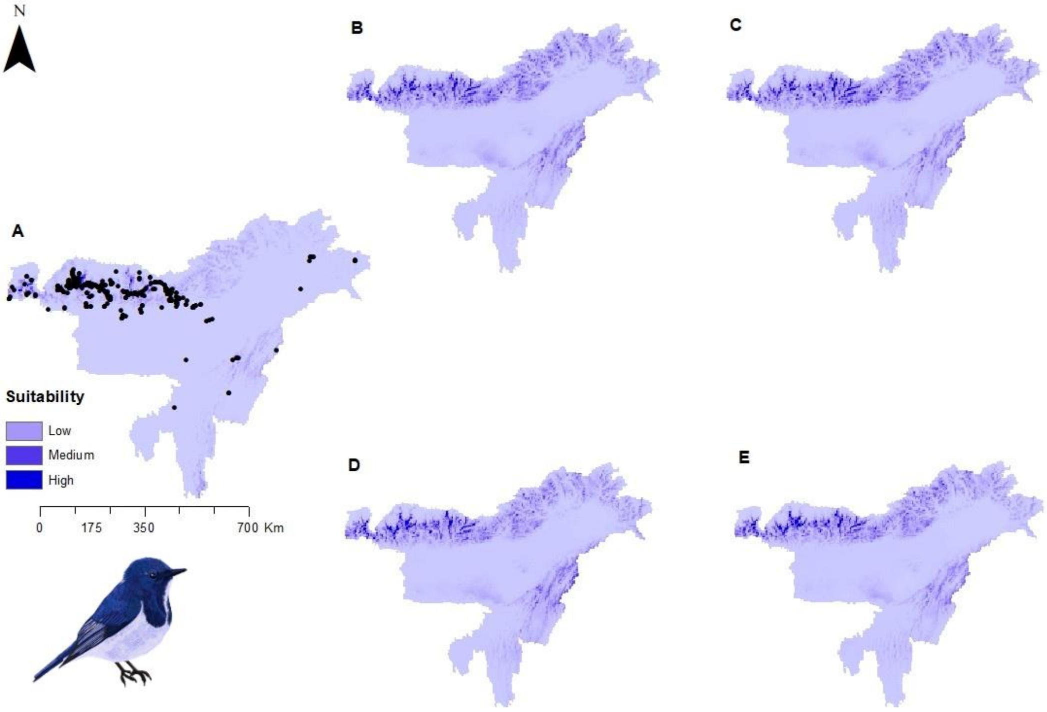


Figure S23: Potential distributions of *Ficedula superciliaris* (Ultramarine Flycatcher) under present climate with presence points (A) and under four future climatic scenarios- (B) SSP1-2.6 (low), (C) SSP2-4.5 (intermediate), (D) SSP 4-6.0 (medium) and (E) SSP5-8.5 (high)

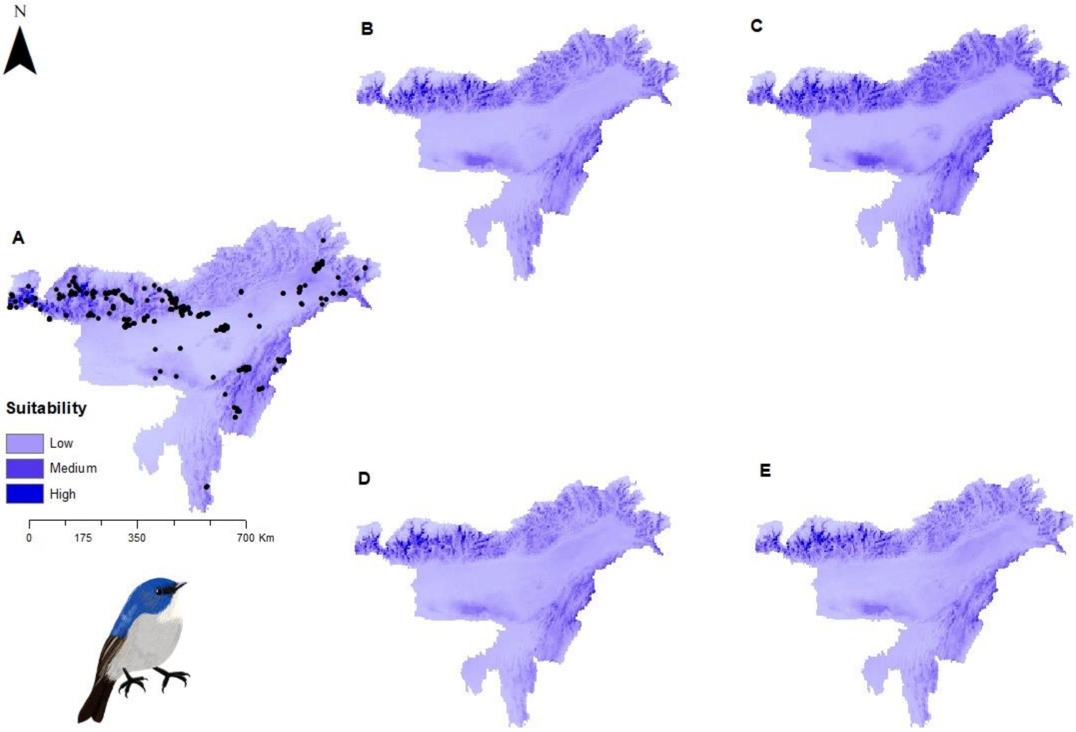


Figure S24: Potential distributions of *Ficedula tricolor* (Slaty-blue Flycatcher) under present climate with presence points (A) and under four future climatic scenarios- (B) SSP1-2.6 (low), (C) SSP2-4.5 (intermediate), (D) SSP 4-6.0 (medium) and (E) SSP5-8.5 (high)

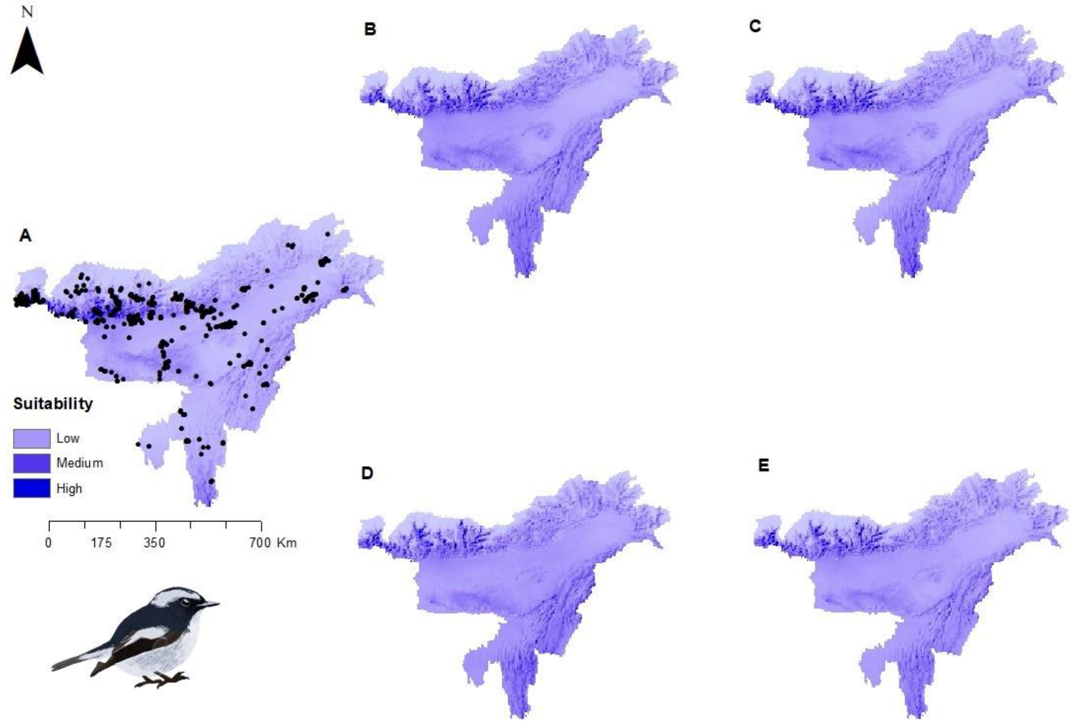


Figure S25: Potential distributions of *Ficedula westermanni* (Little Pied Flycatcher) under present climate with presence points (A) and under four future climatic scenarios- (B) SSP1-2.6 (low), (C) SSP2-4.5 (intermediate), (D) SSP 4-6.0 (medium) and (E) SSP5-8.5 (high)

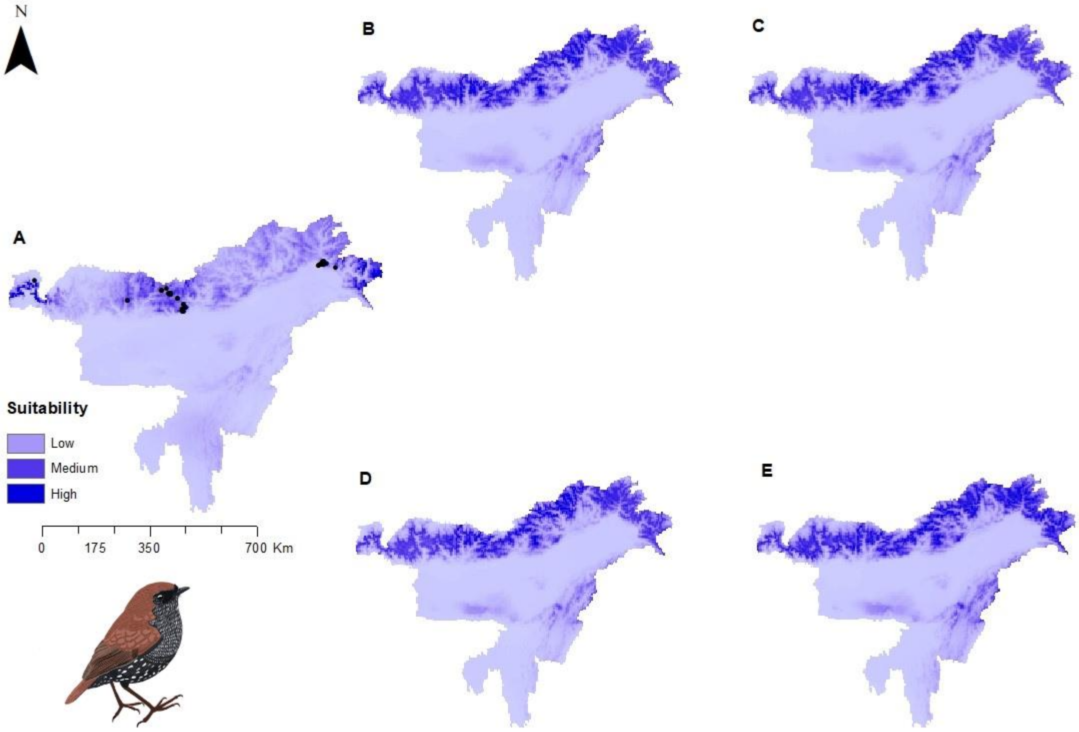


Figure S26: Potential distributions of *Heteroxenicus stellatus* (Gould's Shortwing) under present climate with presence points (A) and under four future climatic scenarios- (B) SSP1-2.6 (low), (C) SSP2-4.5 (intermediate), (D) SSP 4-6.0 (medium) and (E) SSP5-8.5 (high)

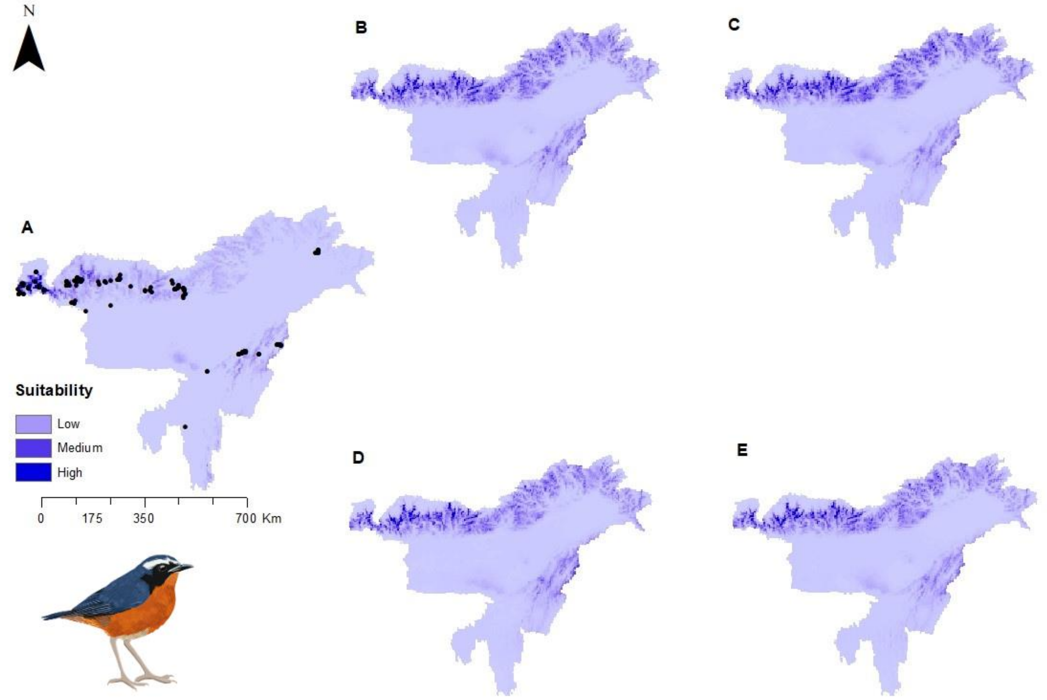


Figure S27: Potential distributions of *Larvivora brunnea* (Indian Blue Robin) under present climate with presence points (A) and under four future climatic scenarios- (B) SSP1-2.6 (low), (C) SSP2-4.5 (intermediate), (D) SSP 4-6.0 (medium) and (E) SSP5-8.5 (high)

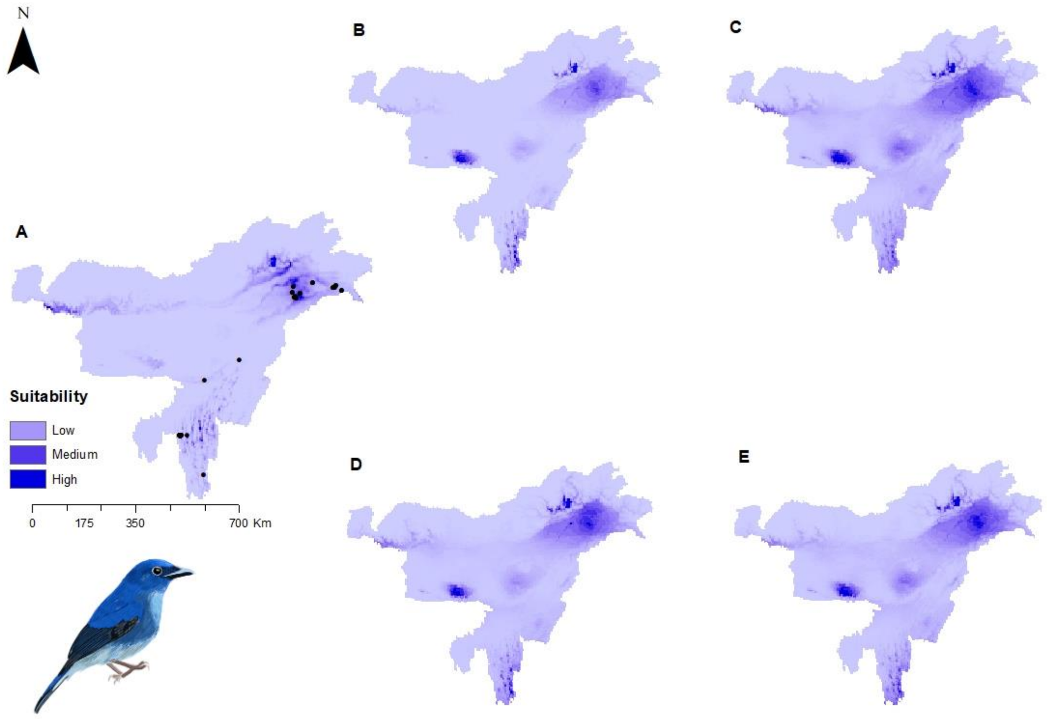


Figure S28: Potential distributions of *Leucoptilon concretum* (White-tailed Flycatcher) under present climate with presence points (A) and under four future climatic scenarios- (B) SSP1-2.6 (low), (C) SSP2-4.5 (intermediate), (D) SSP 4-6.0 (medium) and (E) SSP5-8.5 (high)

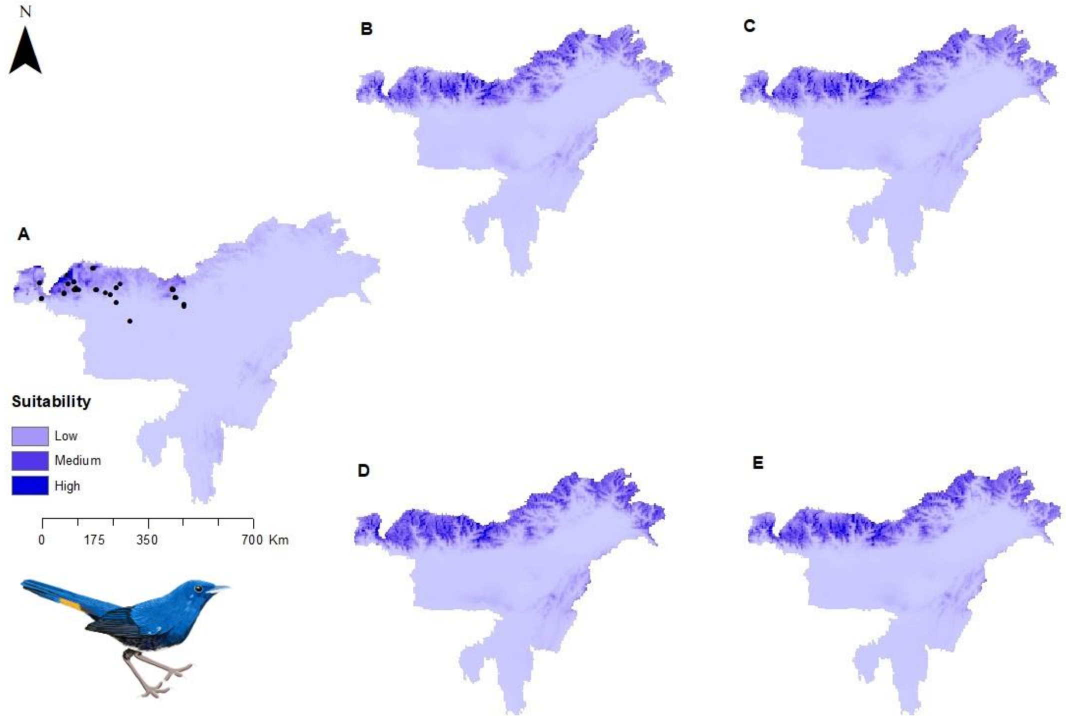


Figure S29: Potential distributions of *Luscinia phaenicuroides* (White-bellied Redstart) under present climate with presence points (A) and under four future climatic scenarios- (B) SSP1-2.6 (low), (C) SSP2-4.5 (intermediate), (D) SSP 4-6.0 (medium) and (E) SSP5-8.5 (high)

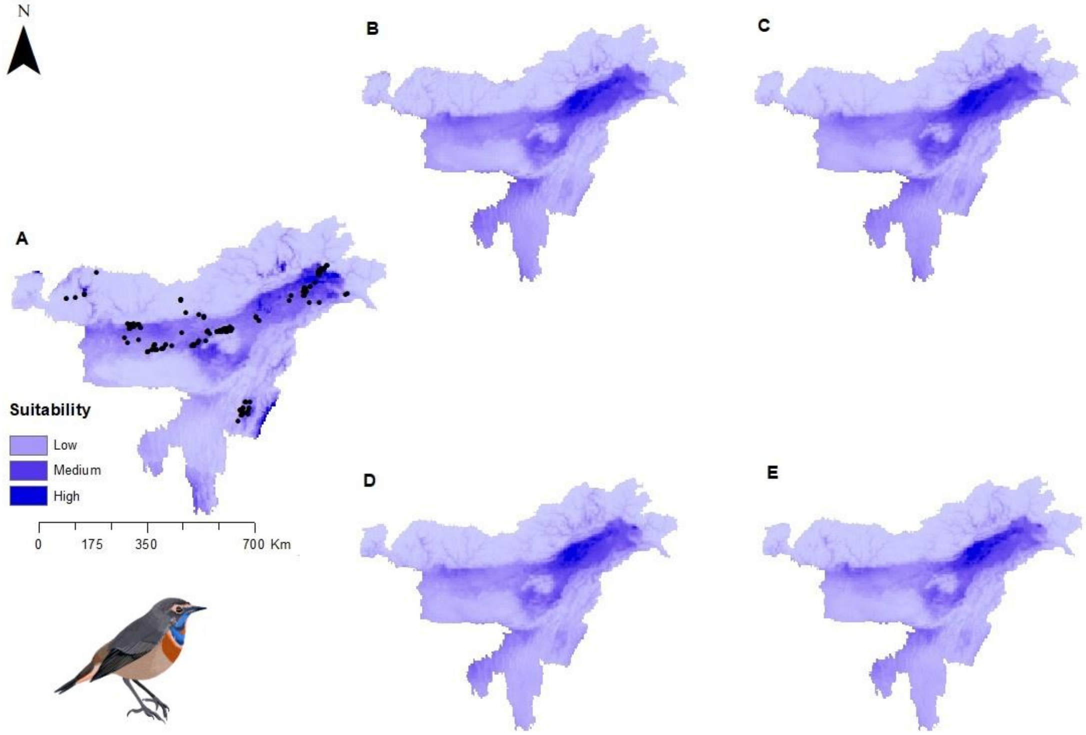


Figure S30: Potential distributions of *Luscinia svecica* (Bluethroat) under present climate with presence points (A) and under four future climatic scenarios- (B) SSP1-2.6 (low), (C) SSP2-4.5 (intermediate), (D) SSP 4-6.0 (medium) and (E) SSP5-8.5 (high)

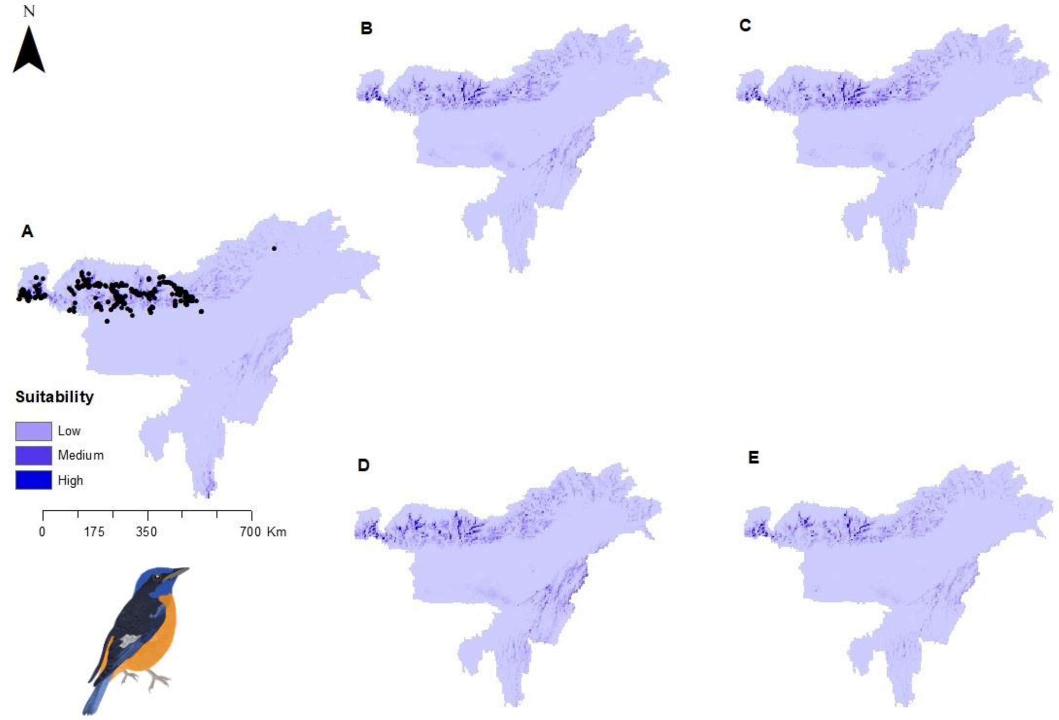


Figure S31: Potential distributions of *Monticola cinclorhyncha* (Blue-capped Rock Thrush) under present climate with presence points (A) and under four future climatic scenarios- (B) SSP1-2.6 (low), (C) SSP2-4.5 (intermediate), (D) SSP 4-6.0 (medium) and (E) SSP5-8.5 (high)

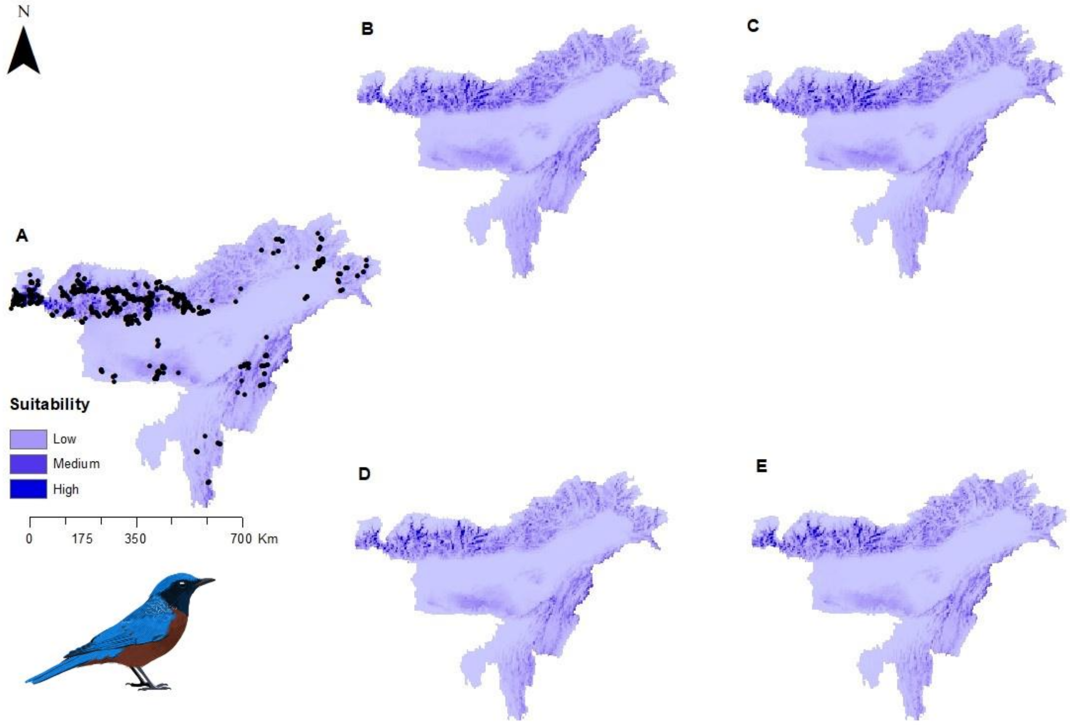


Figure S32: Potential distributions of *Monticola rufiventris* (Chestnut-bellied Rock Thrush) under present climate with presence points (A) and under four future climatic scenarios- (B) SSP1-2.6 (low), (C) SSP2-4.5 (intermediate), (D) SSP 4-6.0 (medium) and (E) SSP5-8.5 (high)

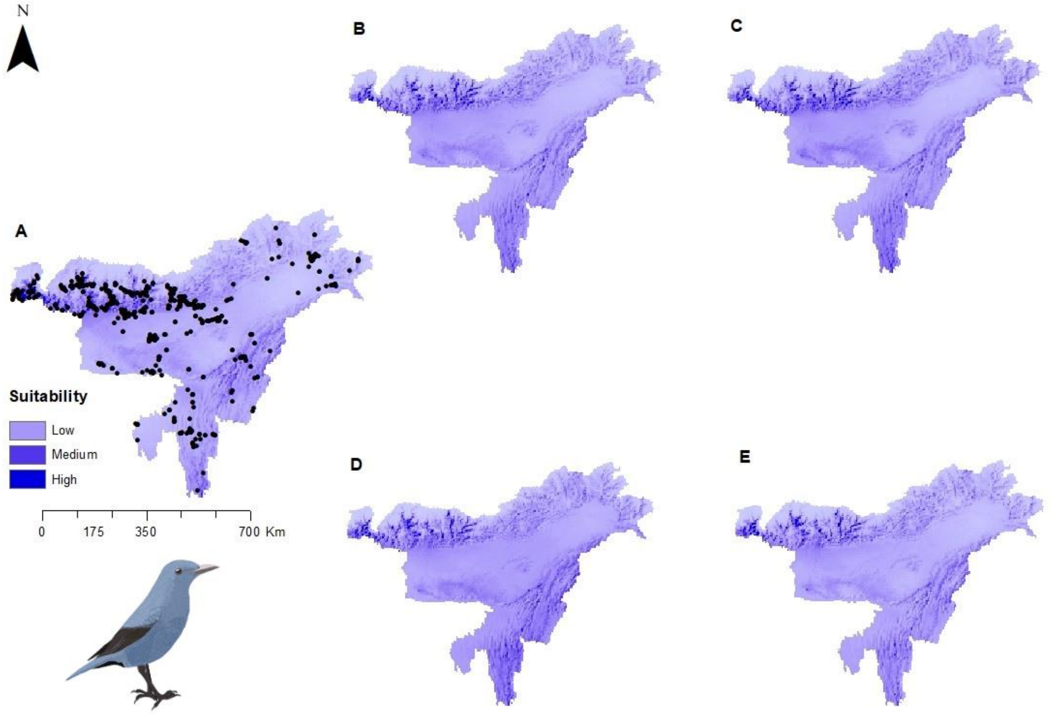


Figure S33: Potential distributions of *Monticola solitarius* (Blue Rock-Thrush) under present climate with presence points (A) and under four future climatic scenarios- (B) SSP1-2.6 (low), (C) SSP2-4.5 (intermediate), (D) SSP 4-6.0 (medium) and (E) SSP5-8.5 (high)

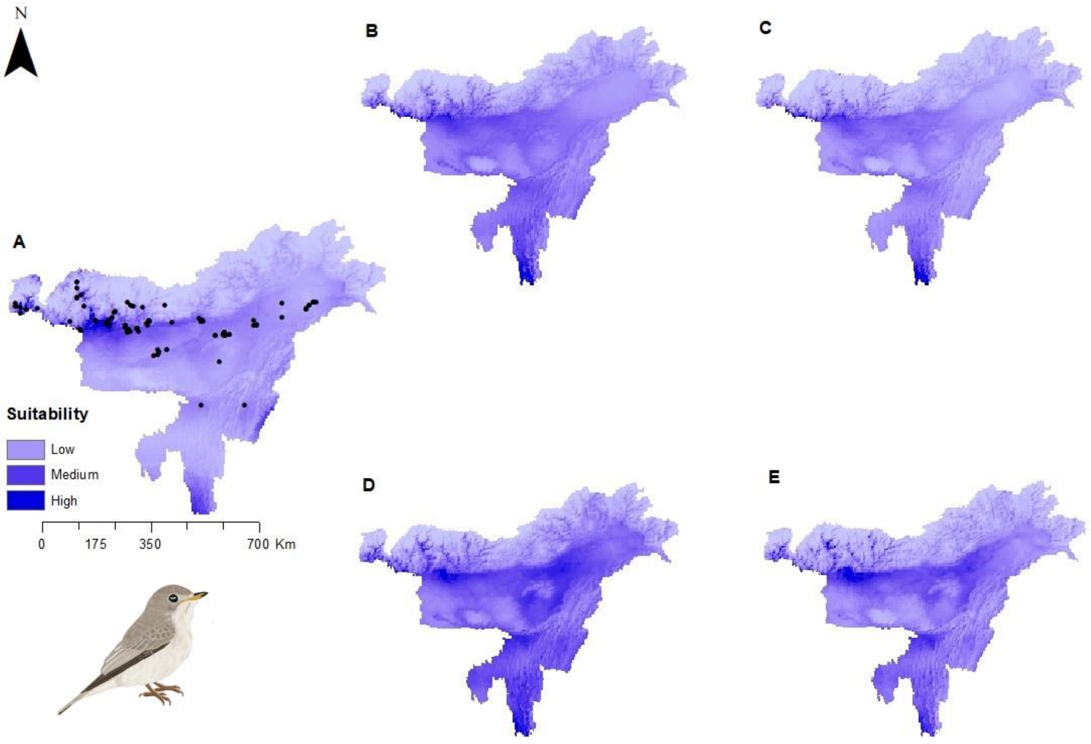


Figure S34: Potential distributions of *Muscicapa dauurica* (Asian Brown Flycatcher) under present climate with presence points (A) and under four future climatic scenarios- (B) SSP1-2.6 (low), (C) SSP2-4.5 (intermediate), (D) SSP 4-6.0 (medium) and (E) SSP5-8.5 (high)

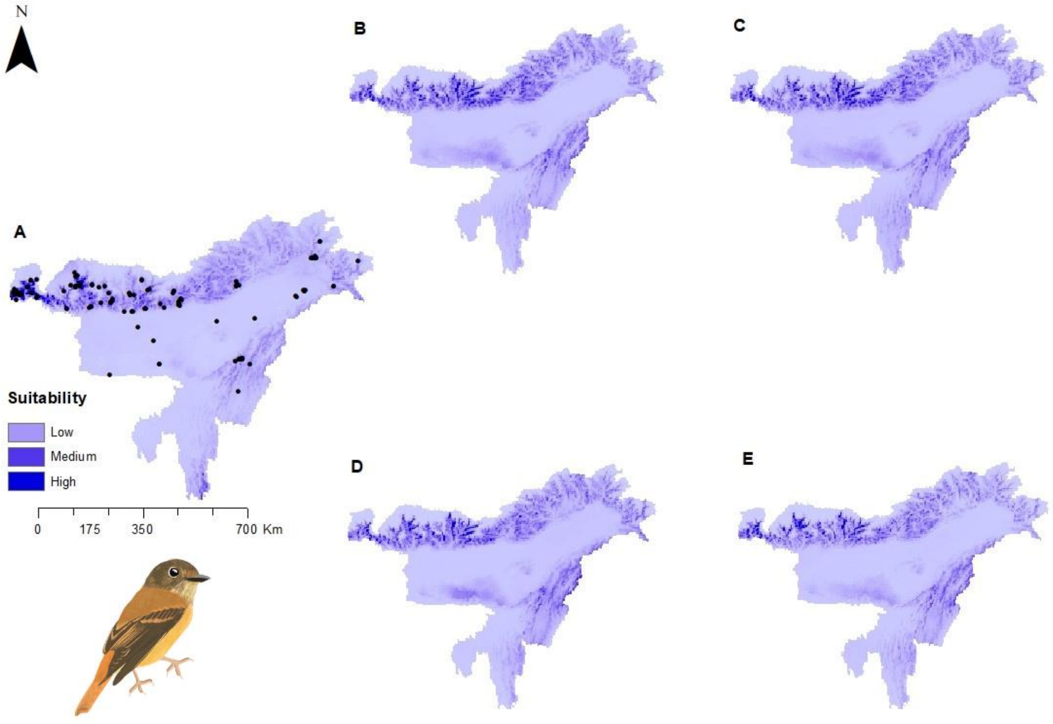


Figure S35: Potential distributions of *Muscicapa ferruginea* (Ferruginous Flycatcher) under present climate with presence points (A) and under four future climatic scenarios- (B) SSP1-2.6 (low), (C) SSP2-4.5 (intermediate), (D) SSP 4-6.0 (medium) and (E) SSP5-8.5 (high)

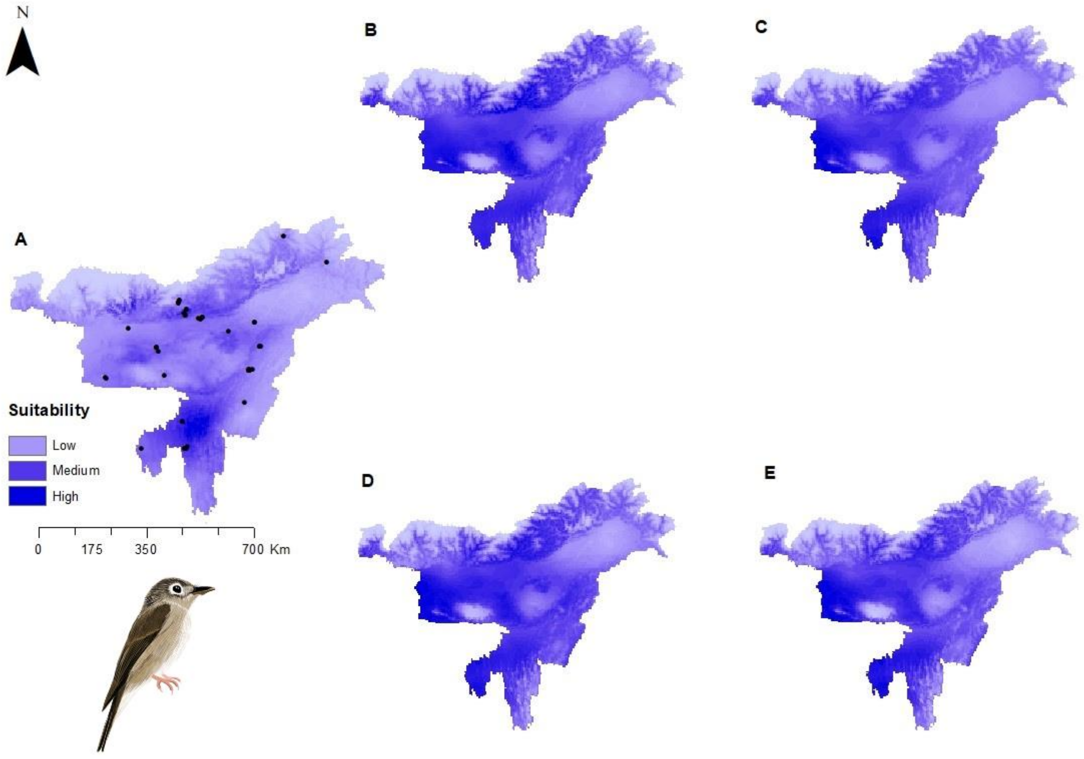


Figure S36: Potential distributions of *Muscicapa muttui* (Brown-breasted Flycatcher) under present climate with presence points (A) and under four future climatic scenarios- (B) SSP1-2.6 (low), (C) SSP2-4.5 (intermediate), (D) SSP 4-6.0 (medium) and (E) SSP5-8.5 (high)

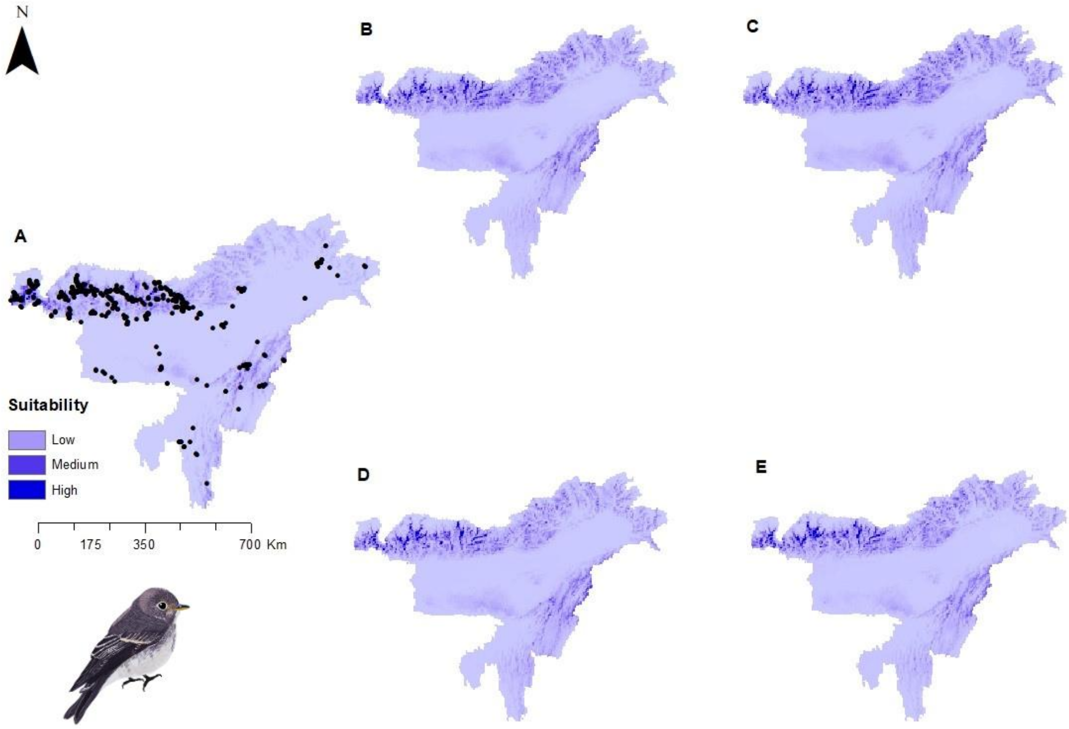


Figure S37: Potential distributions of *Muscicapa sibirica* (Dark-sided Flycatcher) under present climate with presence points (A) and under four future climatic scenarios- (B) SSP1-2.6 (low), (C) SSP2-4.5 (intermediate), (D) SSP 4-6.0 (medium) and (E) SSP5-8.5 (high)

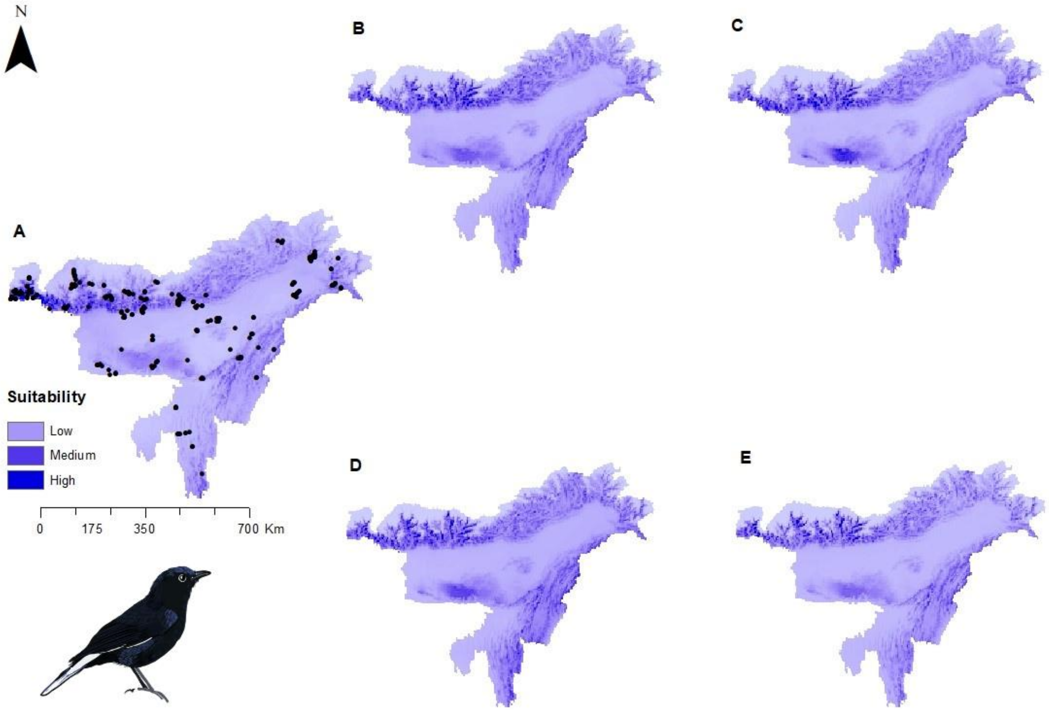


Figure S38: Potential distributions of *Myiomela leucura* (White-tailed Robin) under present climate with presence points (A) and under four future climatic scenarios- (B) SSP1-2.6 (low), (C) SSP2-4.5 (intermediate), (D) SSP 4-6.0 (medium) and (E) SSP5-8.5 (high)

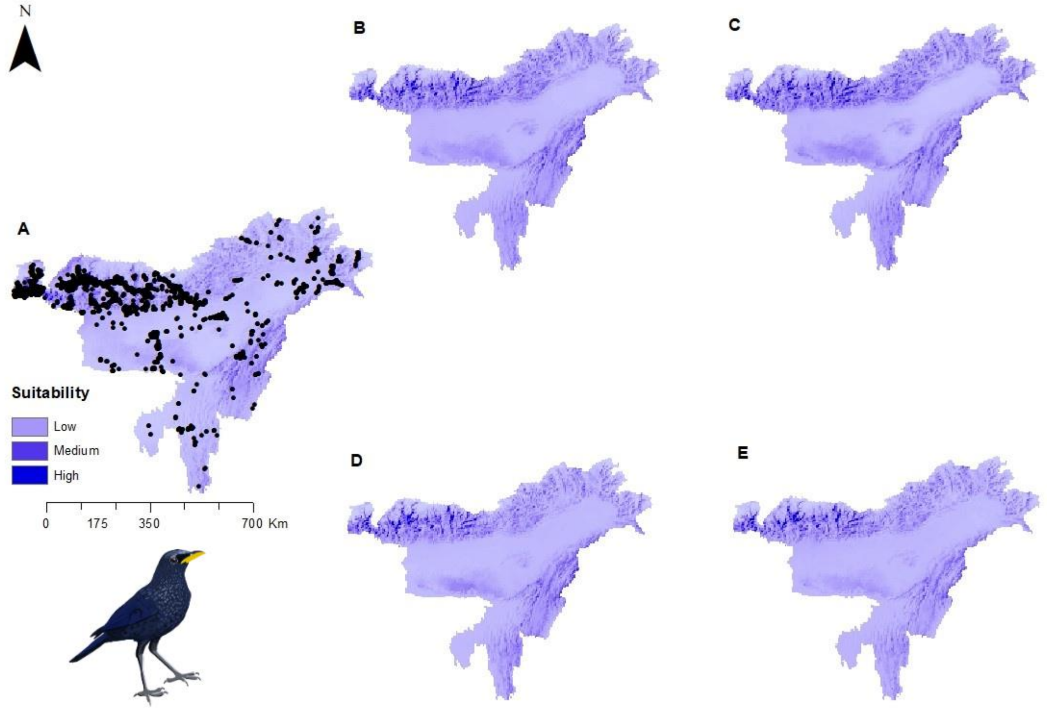


Figure S39: Potential distributions of *Myophonus caeruleus* (Blue Whistling-Thrush) under present climate with presence points (A) and under four future climatic scenarios- (B) SSP1-2.6 (low), (C) SSP2-4.5 (intermediate), (D) SSP 4-6.0 (medium) and (E) SSP5-8.5 (high)

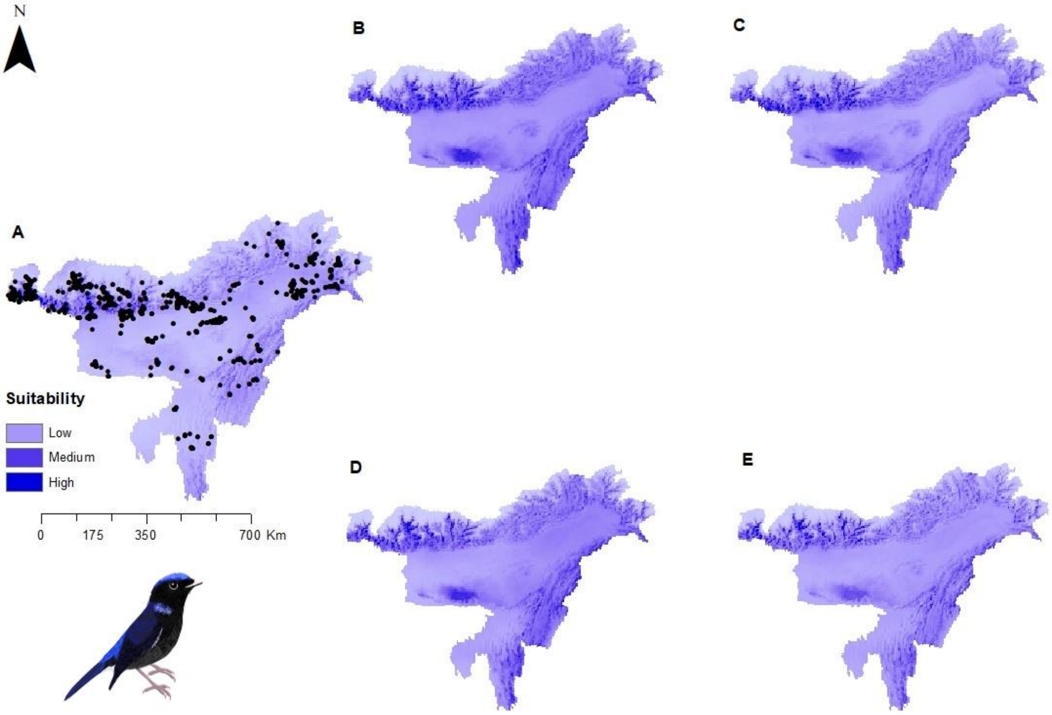


Figure S40: Potential distributions of *Niltava macgrigoriae* (Small Niltava) under present climate with presence points (A) and under four future climatic scenarios- (B) SSP1-2.6 (low), (C) SSP2-4.5 (intermediate), (D) SSP 4-6.0 (medium) and (E) SSP5-8.5 (high)

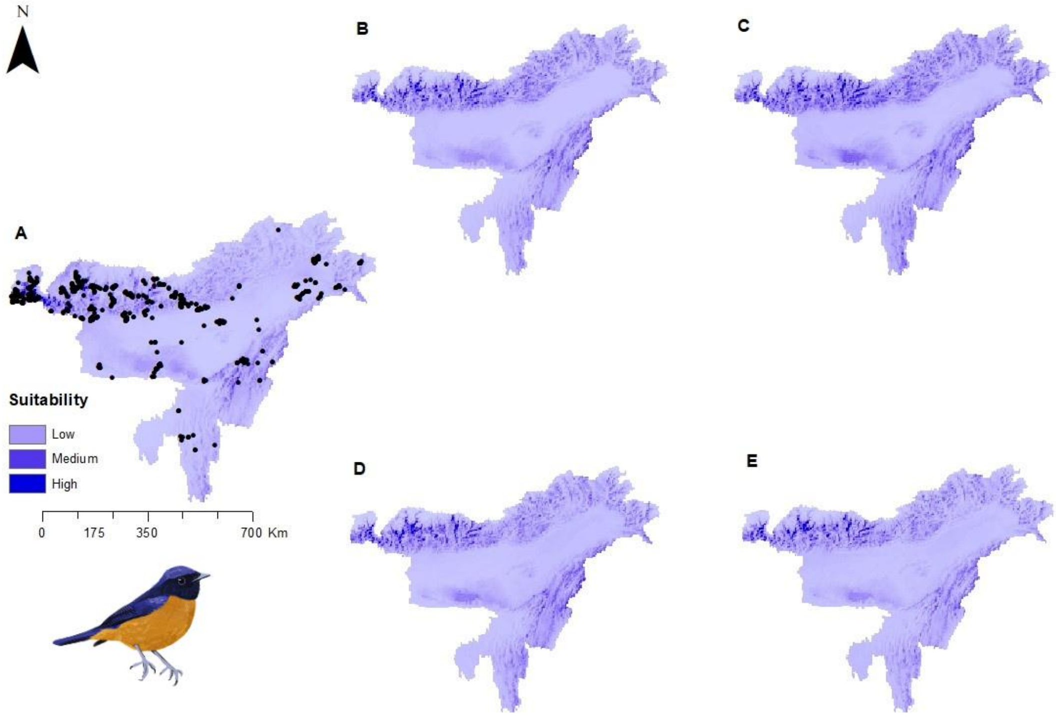


Figure S41: Potential distributions of *Niltava sundara* (Rufous-bellied Niltava) under present climate with presence points (A) and under four future climatic scenarios- (B) SSP1-2.6 (low), (C) SSP2-4.5 (intermediate), (D) SSP 4-6.0 (medium) and (E) SSP5-8.5 (high)

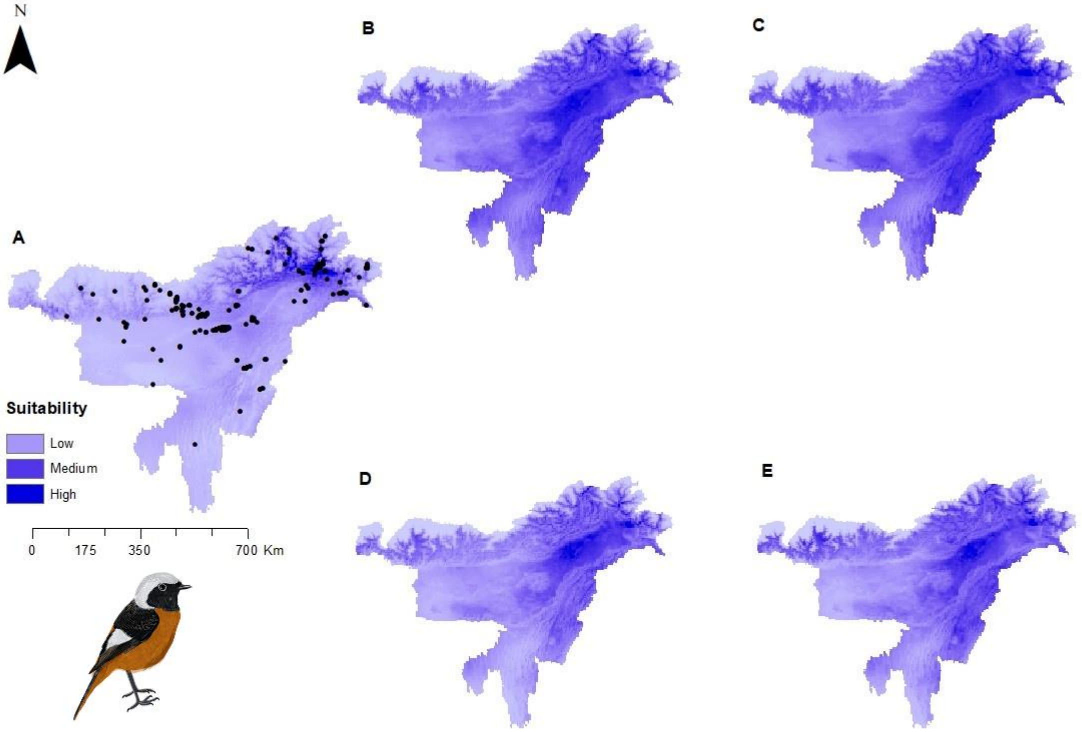


Figure S42: Potential distributions of *Phoenicurus auroreus* (Daurian Redstart) under present climate with presence points (A) and under four future climatic scenarios- (B) SSP1-2.6 (low), (C) SSP2-4.5 (intermediate), (D) SSP 4-6.0 (medium) and (E) SSP5-8.5 (high)

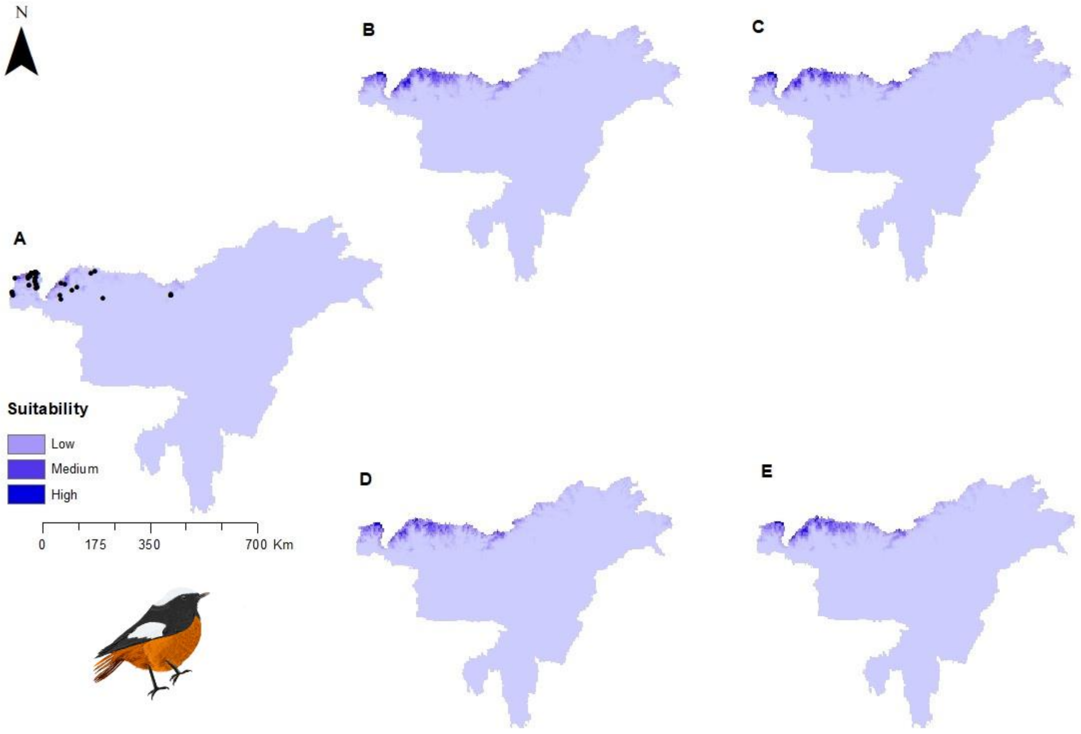


Figure S43: Potential distributions of *Phoenicurus erythrogastrus* (White-winged Redstart) under present climate with presence points (A) and under four future climatic scenarios- (B) SSP1-2.6 (low), (C) SSP2-4.5 (intermediate), (D) SSP 4-6.0 (medium) and (E) SSP5-8.5 (high)

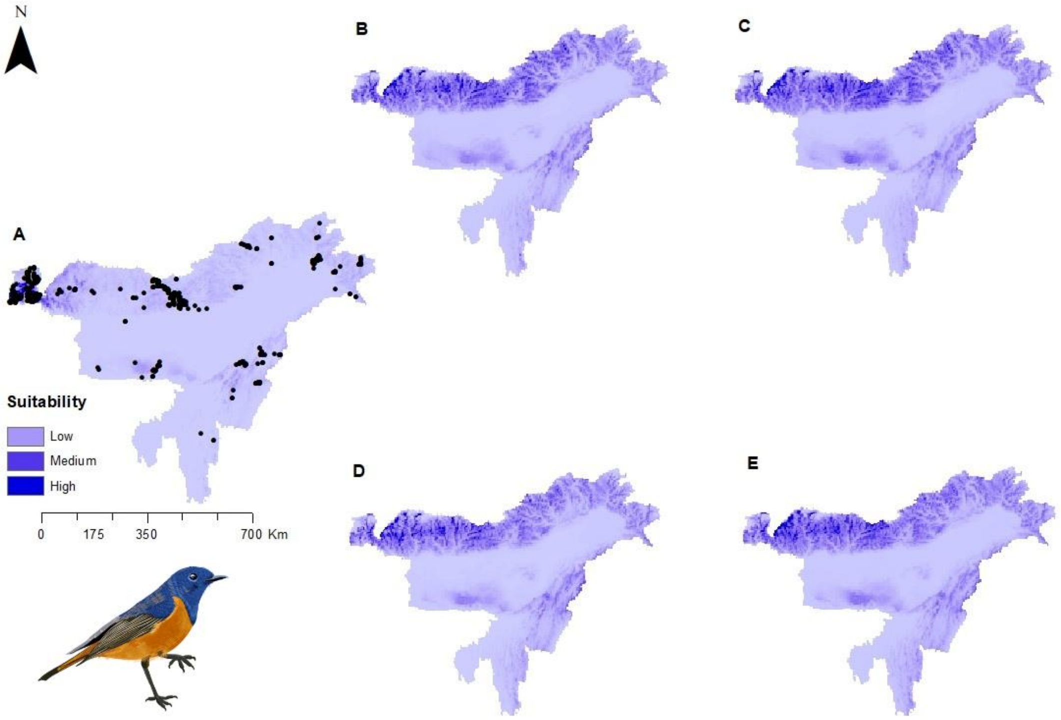


Figure S44: Potential distributions of *Phoenicurus frontalis* (Blue-fronted Redstart) under present climate with presence points (A) and under four future climatic scenarios- (B) SSP1-2.6 (low), (C) SSP2-4.5 (intermediate), (D) SSP 4-6.0 (medium) and (E) SSP5-8.5 (high)

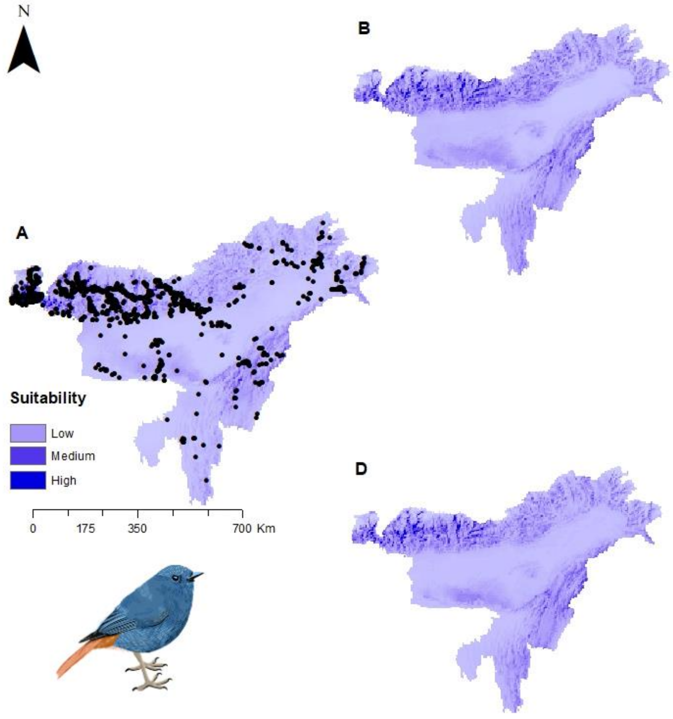


Figure S45: Potential distributions of *Phoenicurus fuliginosus* (Plumbeous Redstart) under present climate with presence points (A) and under four future climatic scenarios- (B) SSP1-2.6 (low), (C) SSP2-4.5 (intermediate), (D) SSP 4-6.0 (medium) and (E) SSP5-8.5 (high)

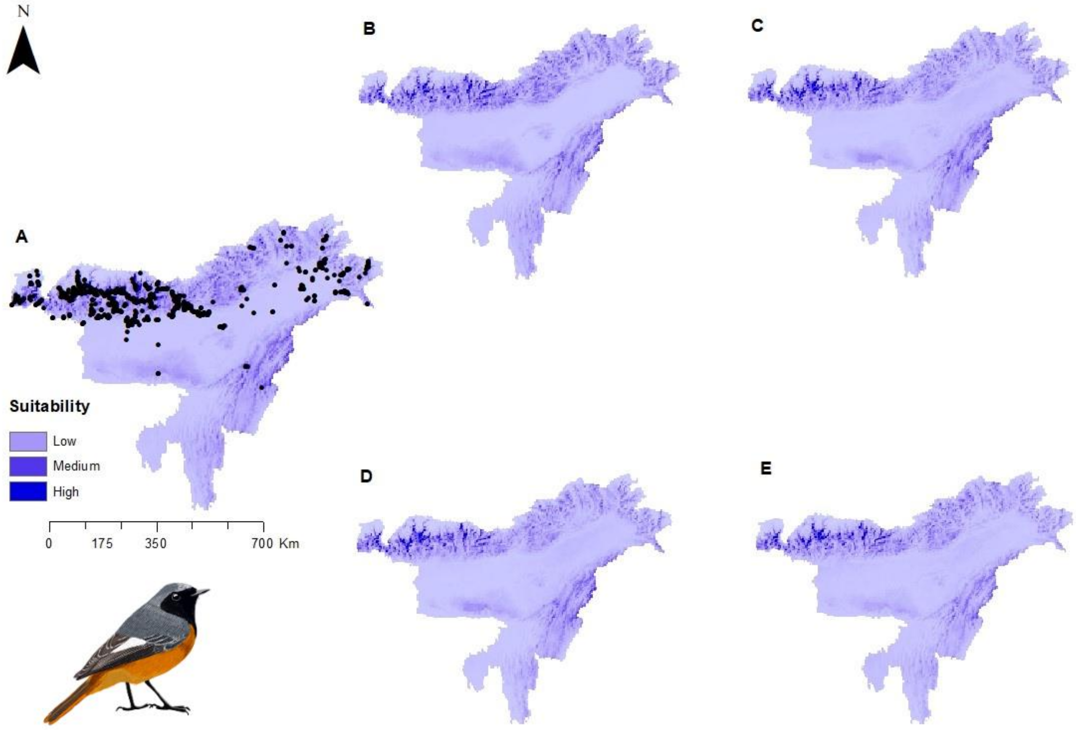


Figure S46: Potential distributions of *Phoenicurus hodgsoni* (Hodgson's Redstart) under present climate with presence points (A) and under four future climatic scenarios- (B) SSP1-2.6 (low), (C) SSP2-4.5 (intermediate), (D) SSP 4-6.0 (medium) and (E) SSP5-8.5 (high)

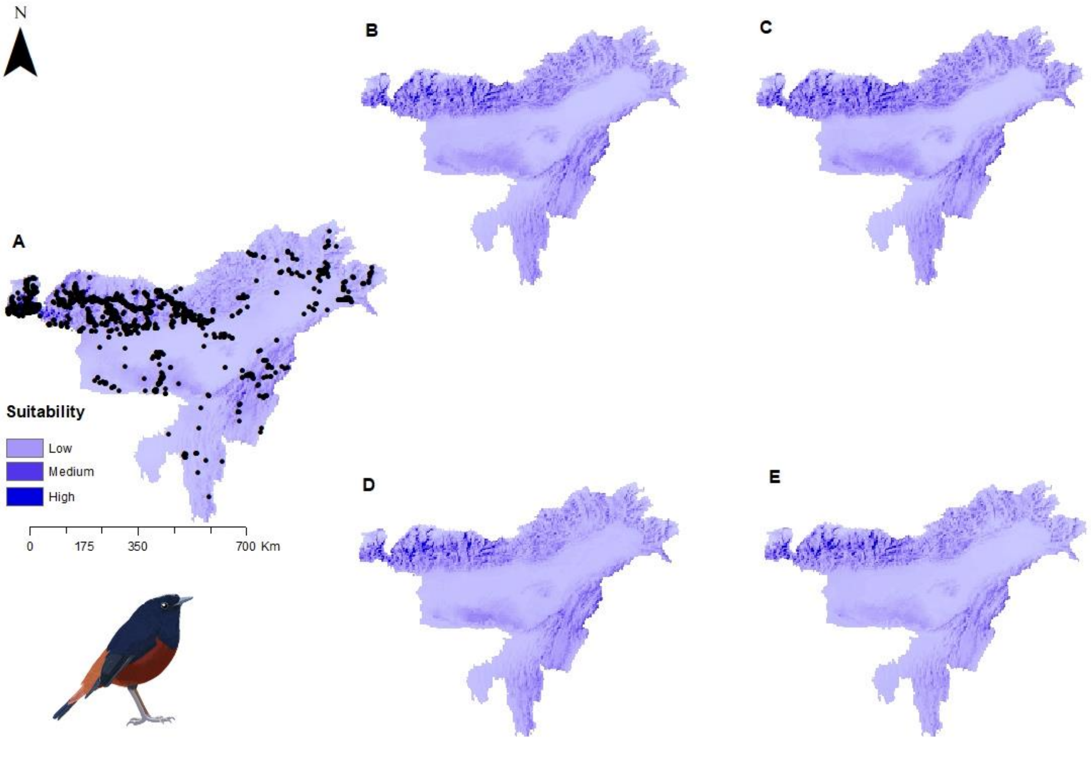


Figure S47: Potential distributions of *Phoenicurus leucocephalus* (White-capped Redstart) under present climate with presence points (A) and under four future climatic scenarios- (B) SSP1-2.6 (low), (C) SSP2-4.5 (intermediate), (D) SSP 4-6.0 (medium) and (E) SSP5-8.5 (high)

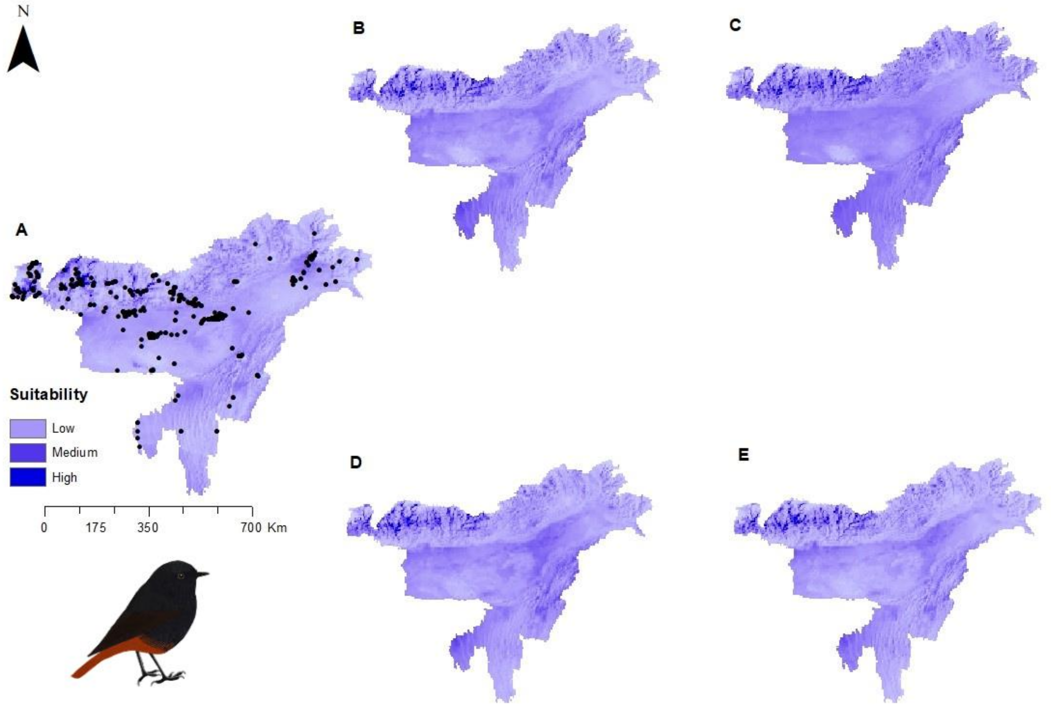


Figure S48: Potential distributions of *Phoenicurus ochruros* (Black Redstart) under present climate with presence points (A) and under four future climatic scenarios- (B) SSP1-2.6 (low), (C) SSP2-4.5 (intermediate), (D) SSP 4-6.0 (medium) and (E) SSP5-8.5 (high)

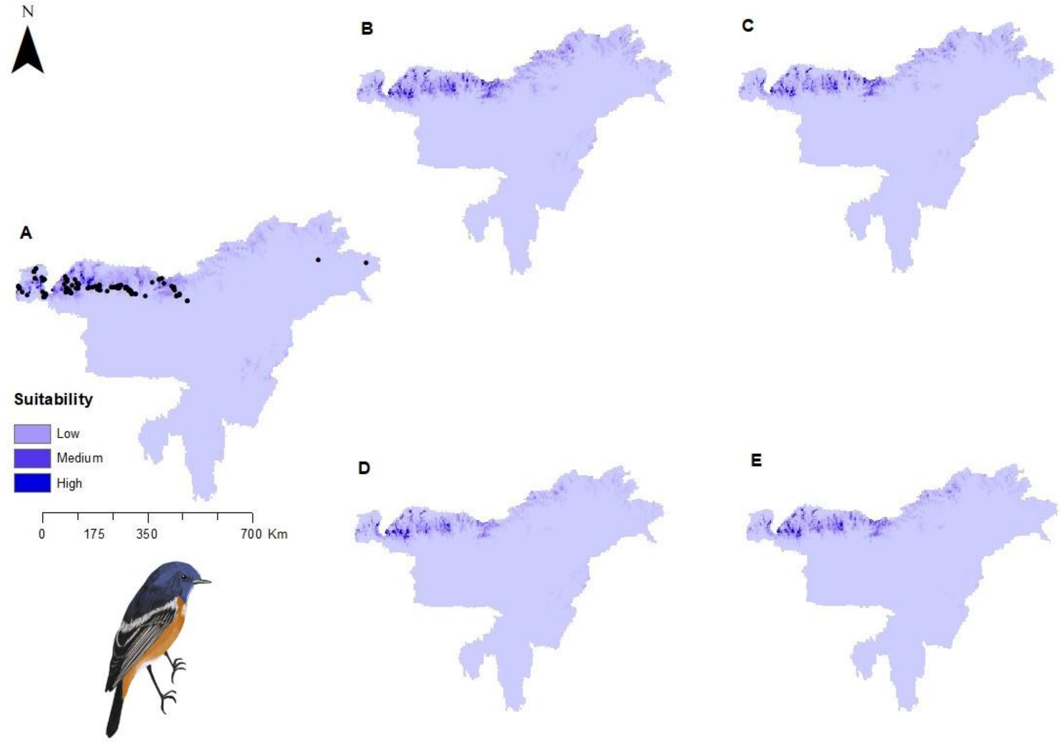


Figure S49: Potential distributions of *Phoenicurus schisticeps* (White-throated Redstart) under present climate with presence points (A) and under four future climatic scenarios- (B) SSP1-2.6 (low), (C) SSP2-4.5 (intermediate), (D) SSP 4-6.0 (medium) and (E) SSP5-8.5 (high)

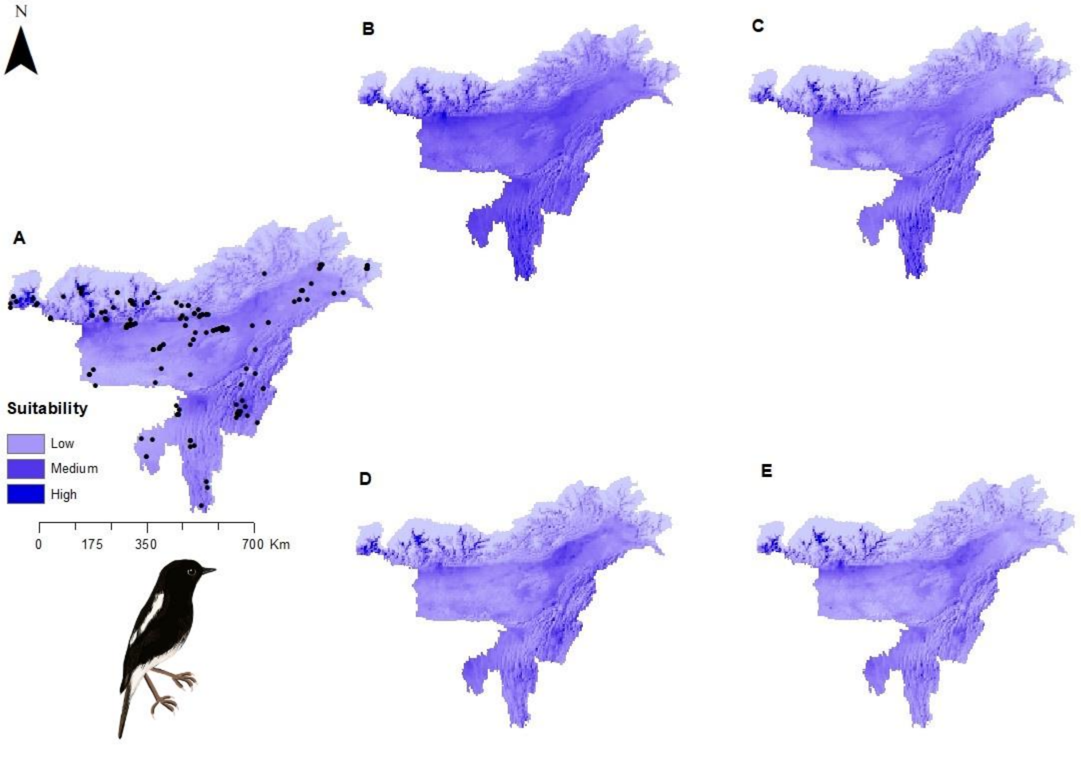


Figure S50: Potential distributions of *Saxicola caprata* (Pied Bushchat) under present climate with presence points (A) and under four future climatic scenarios- (B) SSP1-2.6 (low), (C) SSP2-4.5 (intermediate), (D) SSP 4-6.0 (medium) and (E) SSP5-8.5 (high)

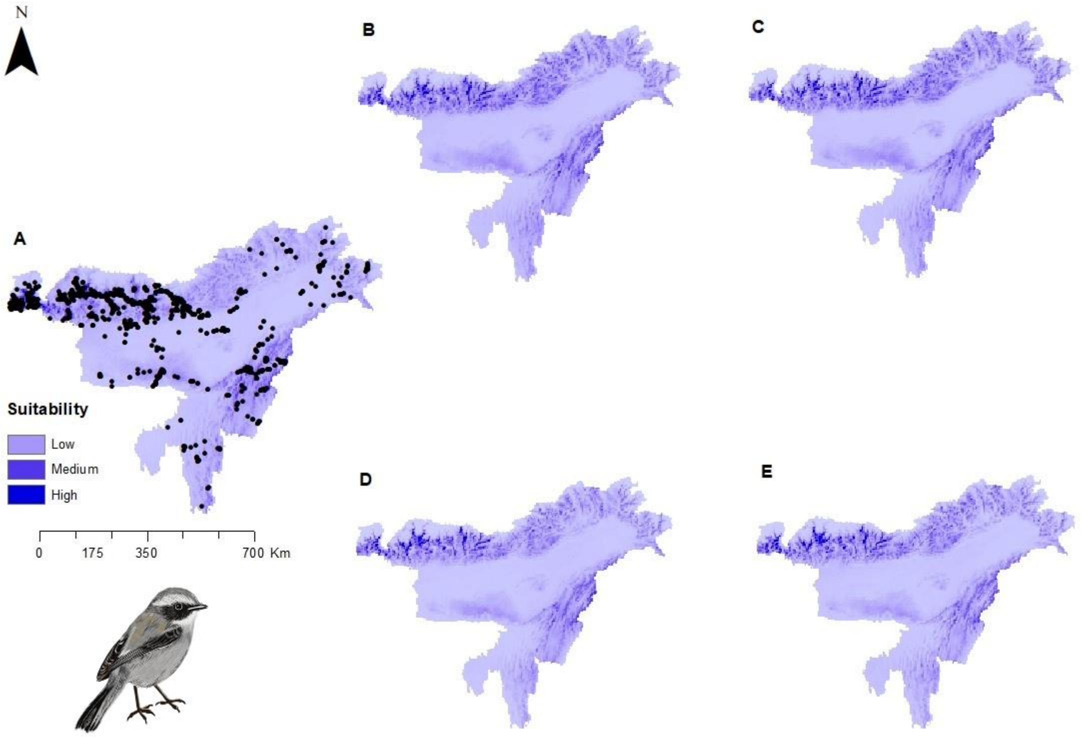


Figure S51: Potential distributions of *Saxicola ferreus* (Grey Bushchat) under present climate with presence points (A) and under four future climatic scenarios- (B) SSP1-2.6 (low), (C) SSP2-4.5 (intermediate), (D) SSP 4-6.0 (medium) and (E) SSP5-8.5 (high)

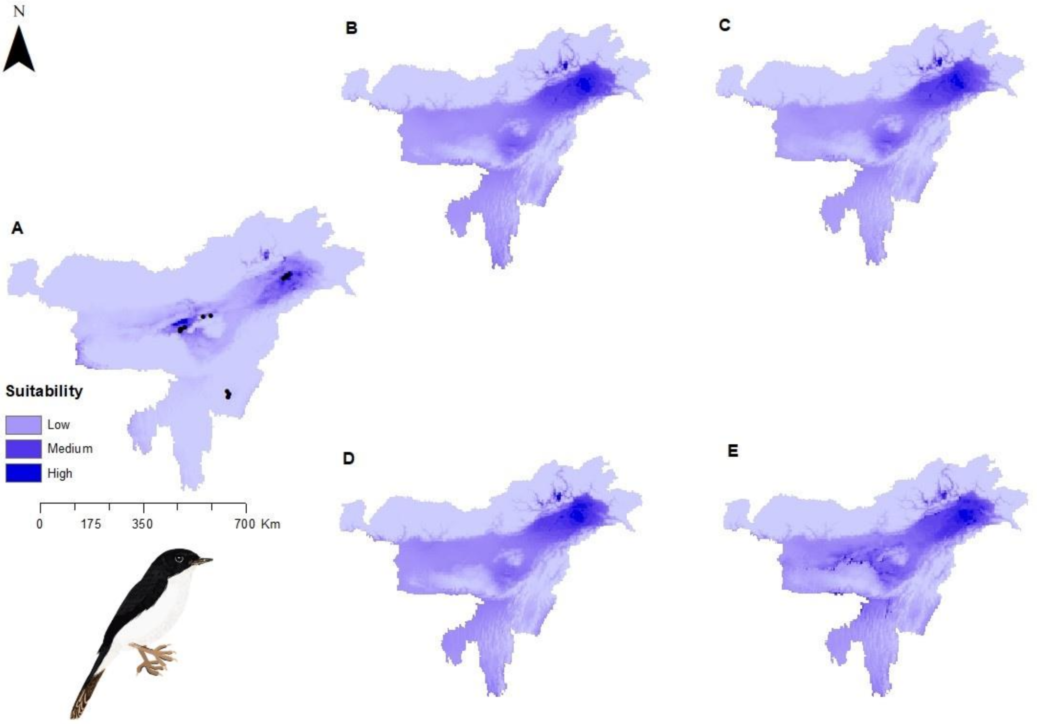


Figure S52: Potential distributions of *Saxicola jerdoni* (Jerdon's Bushchat) under present climate with presence points (A) and under four future climatic scenarios- (B) SSP1-2.6 (low), (C) SSP2-4.5 (intermediate), (D) SSP 4-6.0 (medium) and (E) SSP5-8.5 (high)

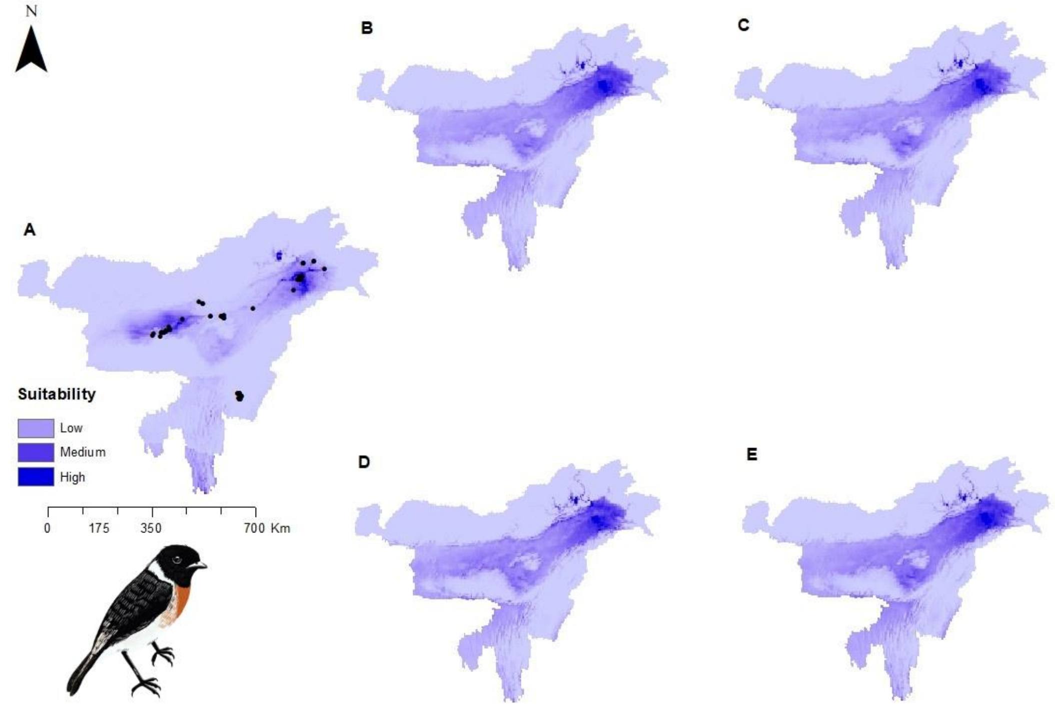


Figure S53: Potential distributions of *Saxicola leucurus* (White-tailed Stonechat) under present climate with presence points (A) and under four future climatic scenarios- (B) SSP1-2.6 (low), (C) SSP2-4.5 (intermediate), (D) SSP 4-6.0 (medium) and (E) SSP5-8.5 (high)

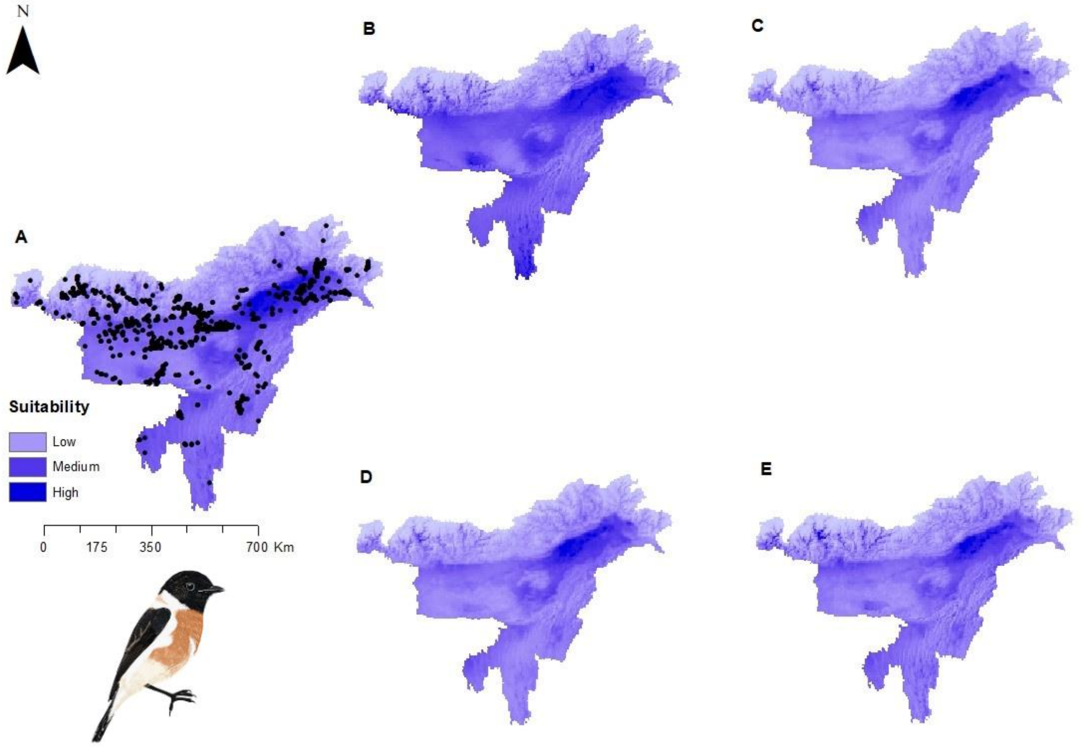


Figure S54: Potential distributions of *Saxicola maurus* (Siberian Stonechat) under present climate with presence points (A) and under four future climatic scenarios- (B) SSP1-2.6 (low), (C) SSP2-4.5 (intermediate), (D) SSP 4-6.0 (medium) and (E) SSP5-8.5 (high)

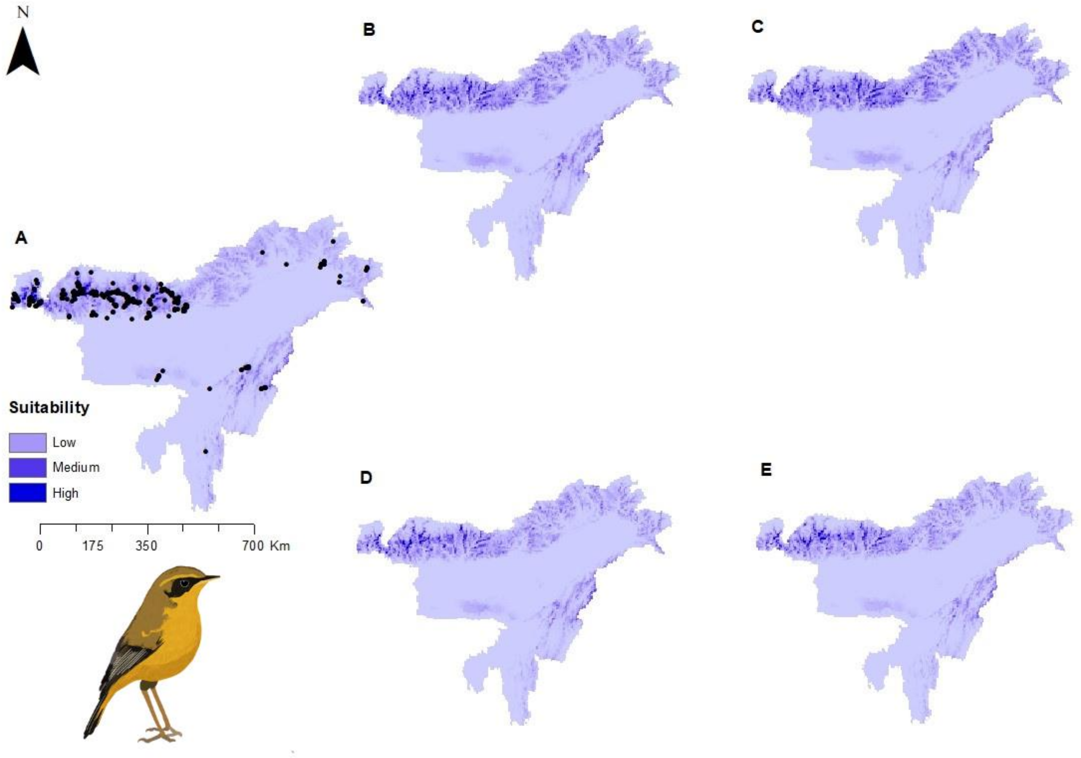


Figure S55: Potential distributions of *Tarsiger chrysaeus* (Golden Bush-Robin) under present climate with presence points (A) and under four future climatic scenarios- (B) SSP1-2.6 (low), (C) SSP2-4.5 (intermediate), (D) SSP 4-6.0 (medium) and (E) SSP5-8.5 (high)

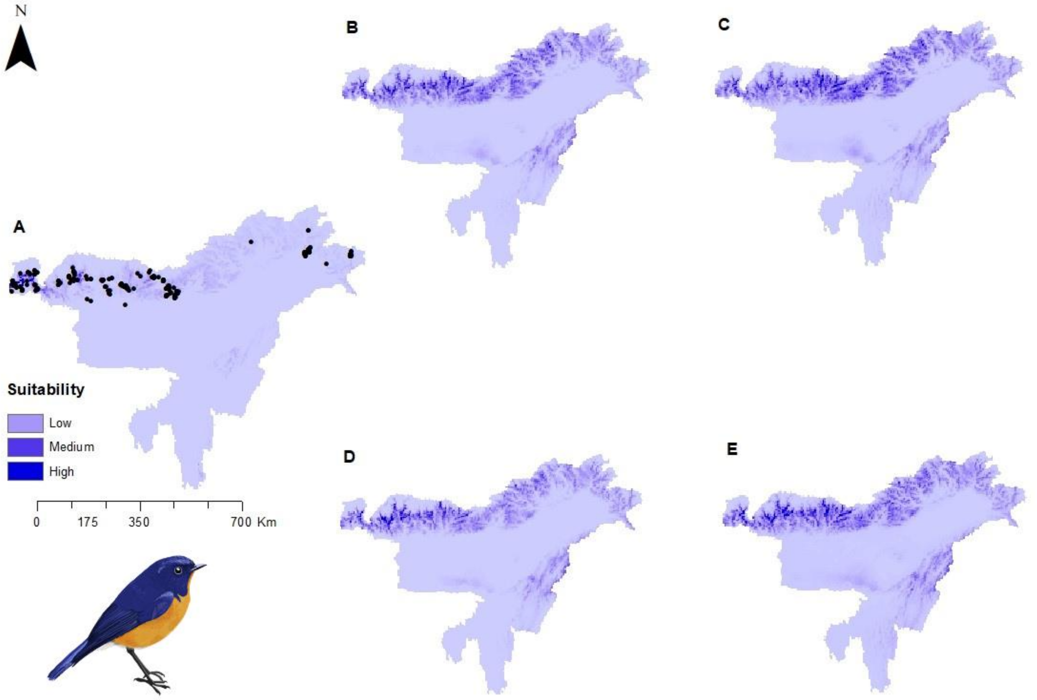


Figure S56: Potential distributions of *Tarsiger hyperythrus* (Rufous-breasted Bush-Robin) under present climate with presence points (A) and under four future climatic scenarios- (B) SSP1-2.6 (low), (C) SSP2-4.5 (intermediate), (D) SSP 4-6.0 (medium) and (E) SSP5-8.5 (high)

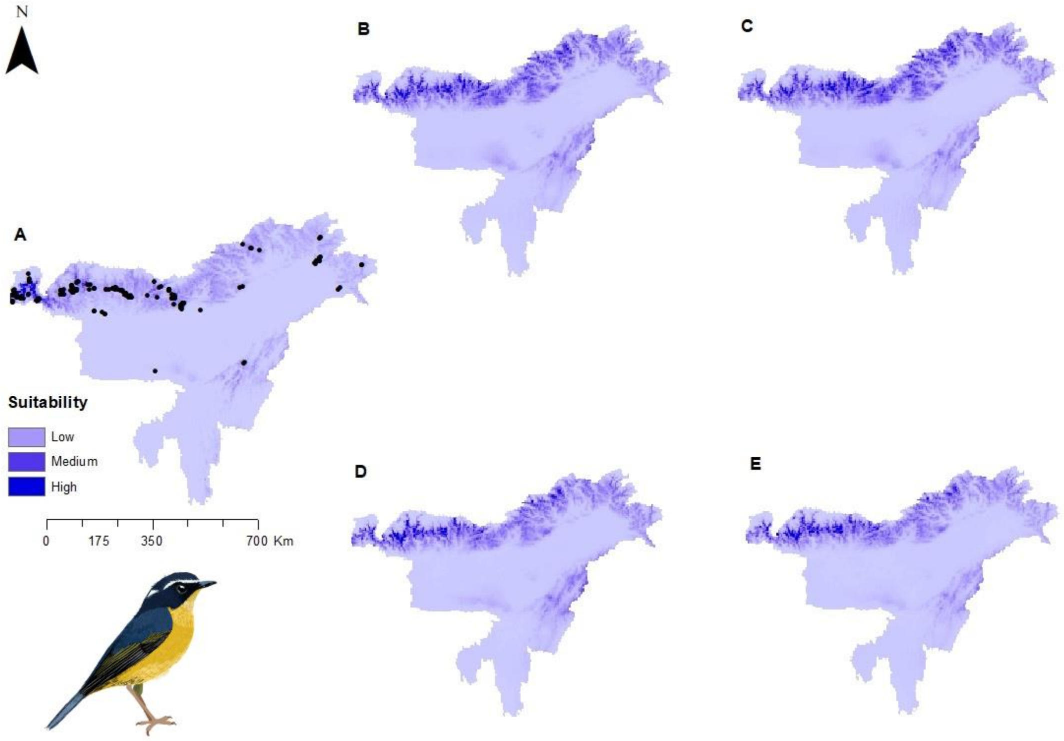


Figure S57: Potential distributions of *Tarsiger indicus* (White-browed Bush-Robin) under present climate with presence points (A) and under four future climatic scenarios- (B) SSP1-2.6 (low), (C) SSP2-4.5 (intermediate), (D) SSP 4-6.0 (medium) and (E) SSP5-8.5 (high)

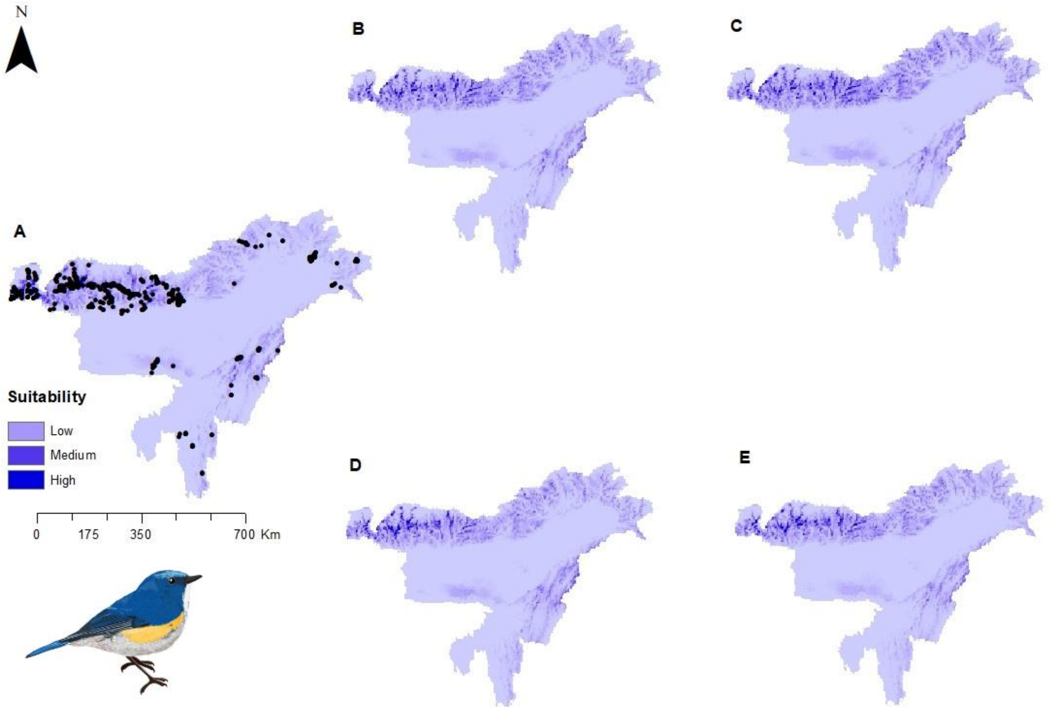


Figure S58: Potential distributions of *Tarsiger rufilatus* (Himalayan Bluetail) under present climate with presence points (A) and under four future climatic scenarios- (B) SSP1-2.6 (low), (C) SSP2-4.5 (intermediate), (D) SSP 4-6.0 (medium) and (E) SSP5-8.5 (high)

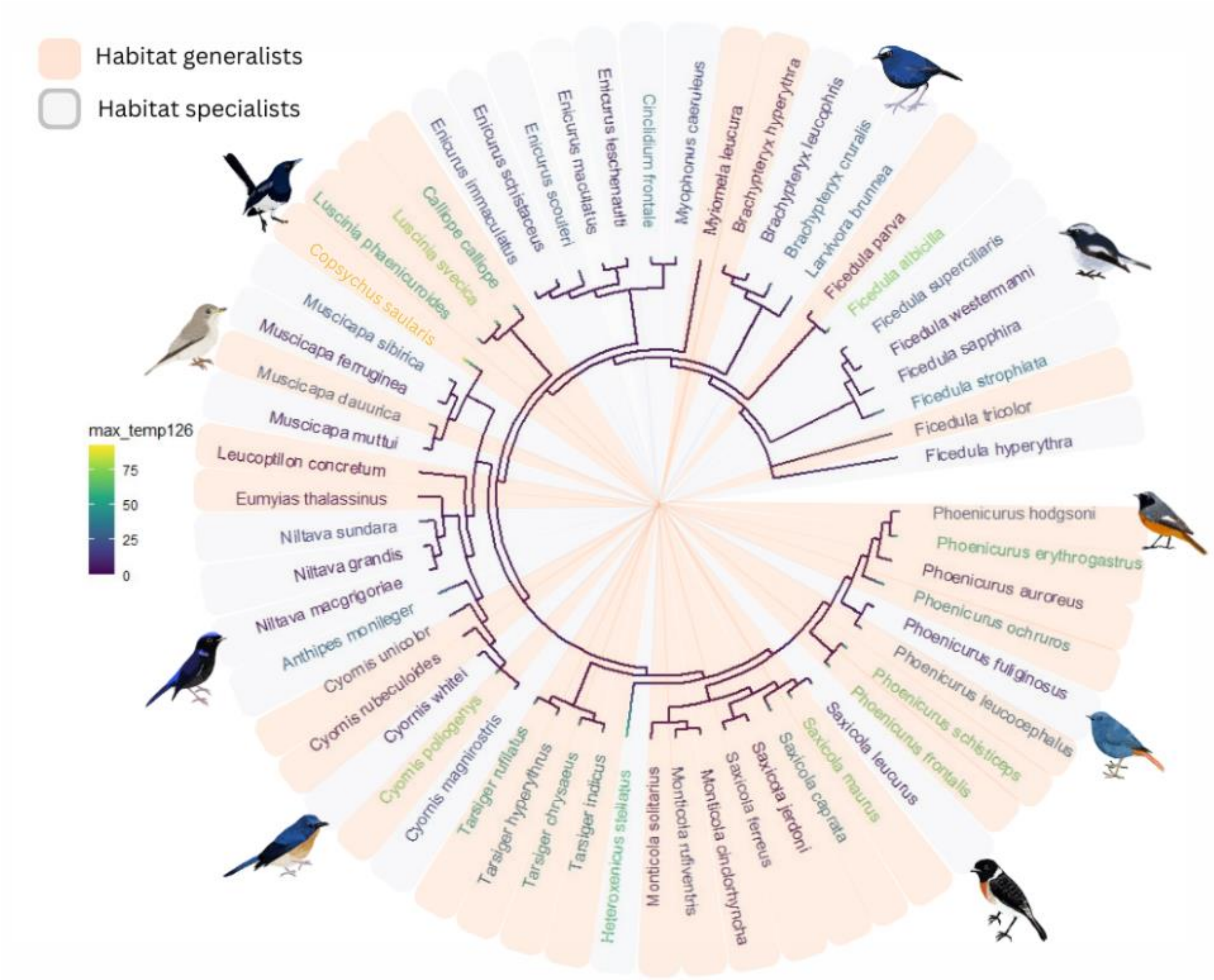


Figure S59: Percentage contribution of tmax (maximum temperature) under future climate scenario- SSP1-2.6 (low) to Ecological Niche Models of Muscicapidae (Flycatchers) species categorised as habitat specialists (light blue) or habitat generalists (light orange)

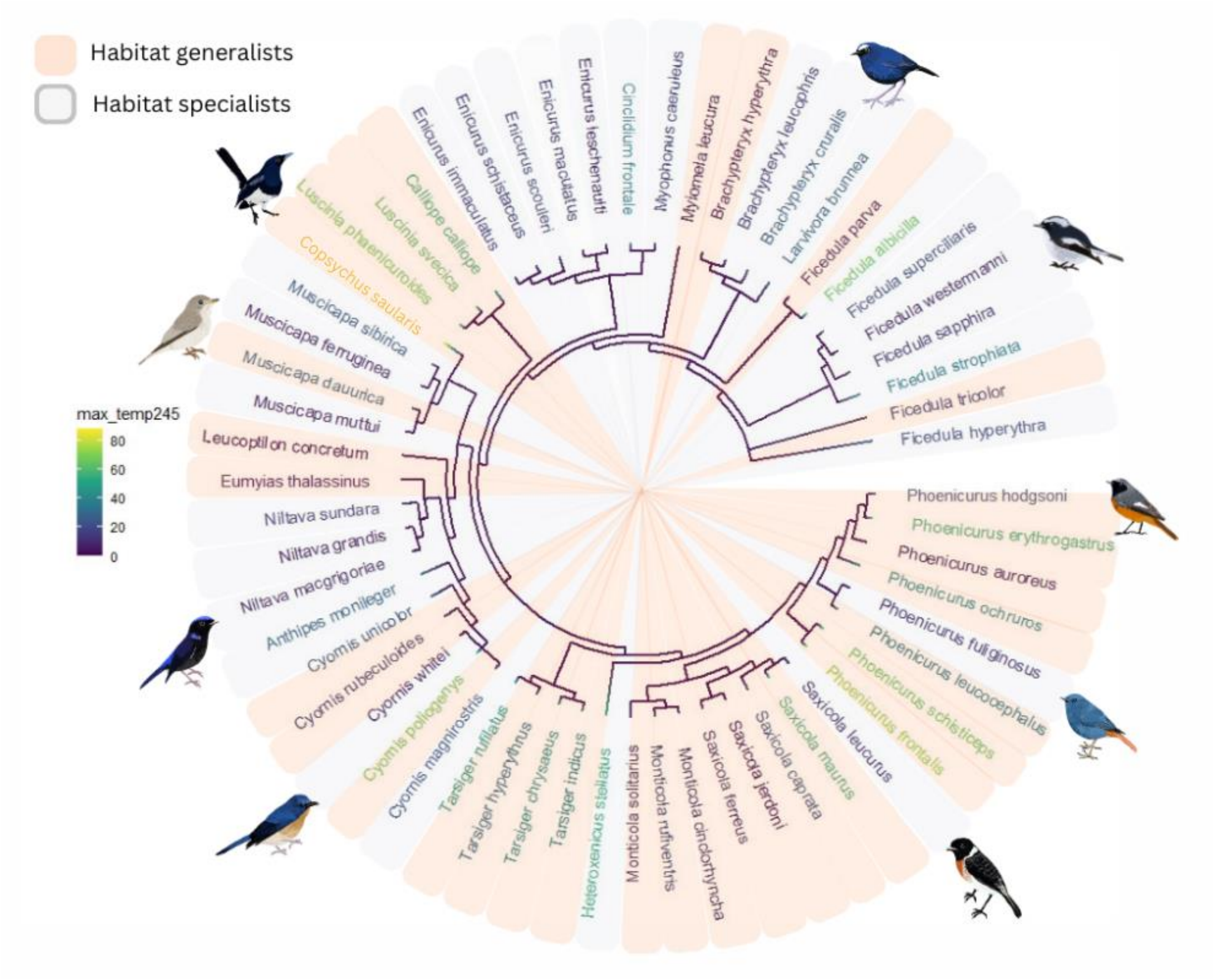


Figure S60: Percentage contribution of tmax (maximum temperature) under future climate scenario- SSP2-4.5 (intermediate) to Ecological Niche Models of Muscicapidae (Flycatchers) species categorised as habitat specialists (light blue) or habitat generalists (light orange)

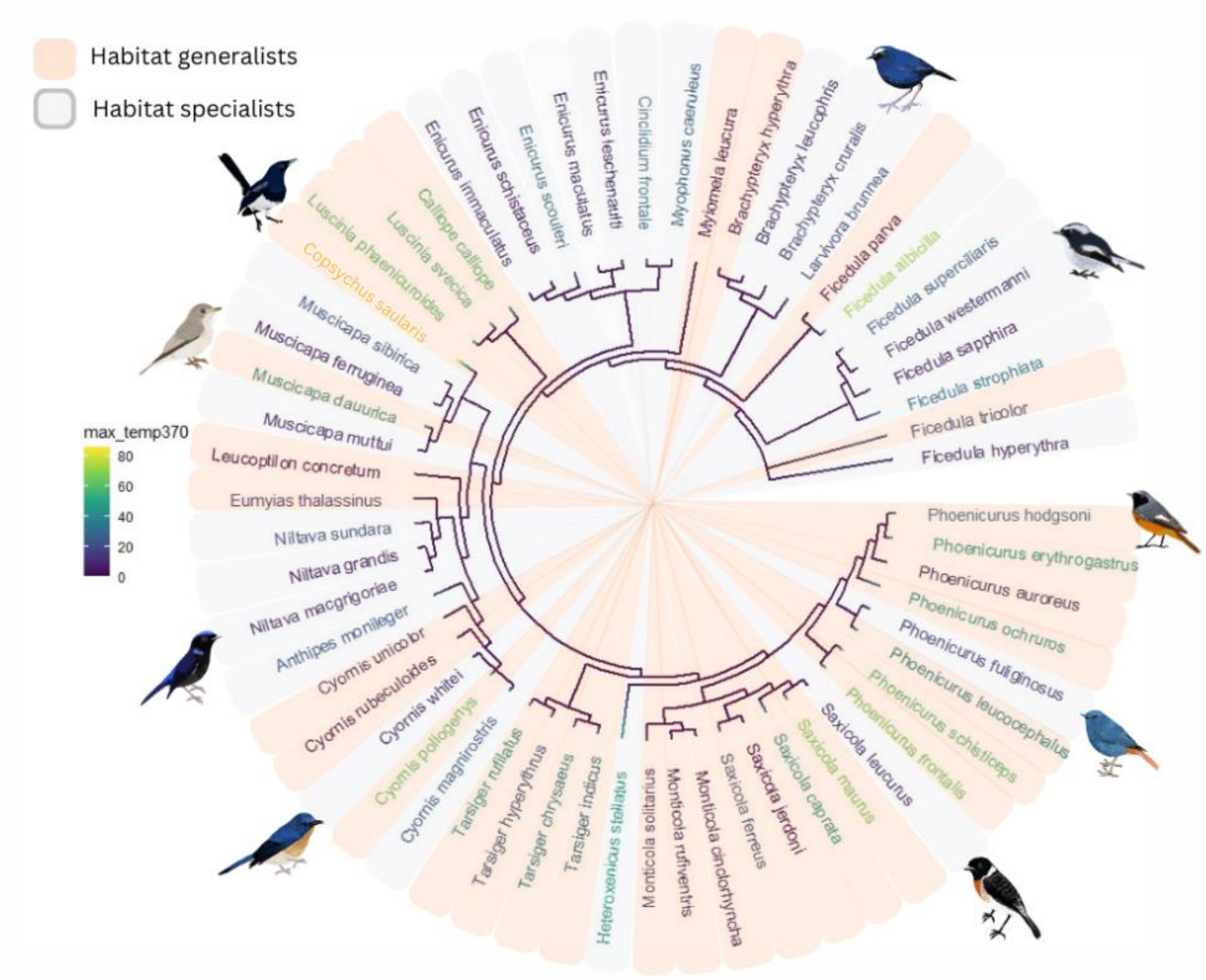


Figure S61: Percentage contribution of tmax (maximum temperature) under future climate scenario- SSP 4-6.0 (medium) to Ecological Niche Models of Muscicapidae (Flycatchers) species categorised as habitat specialists (light blue) or habitat generalists (light orange)

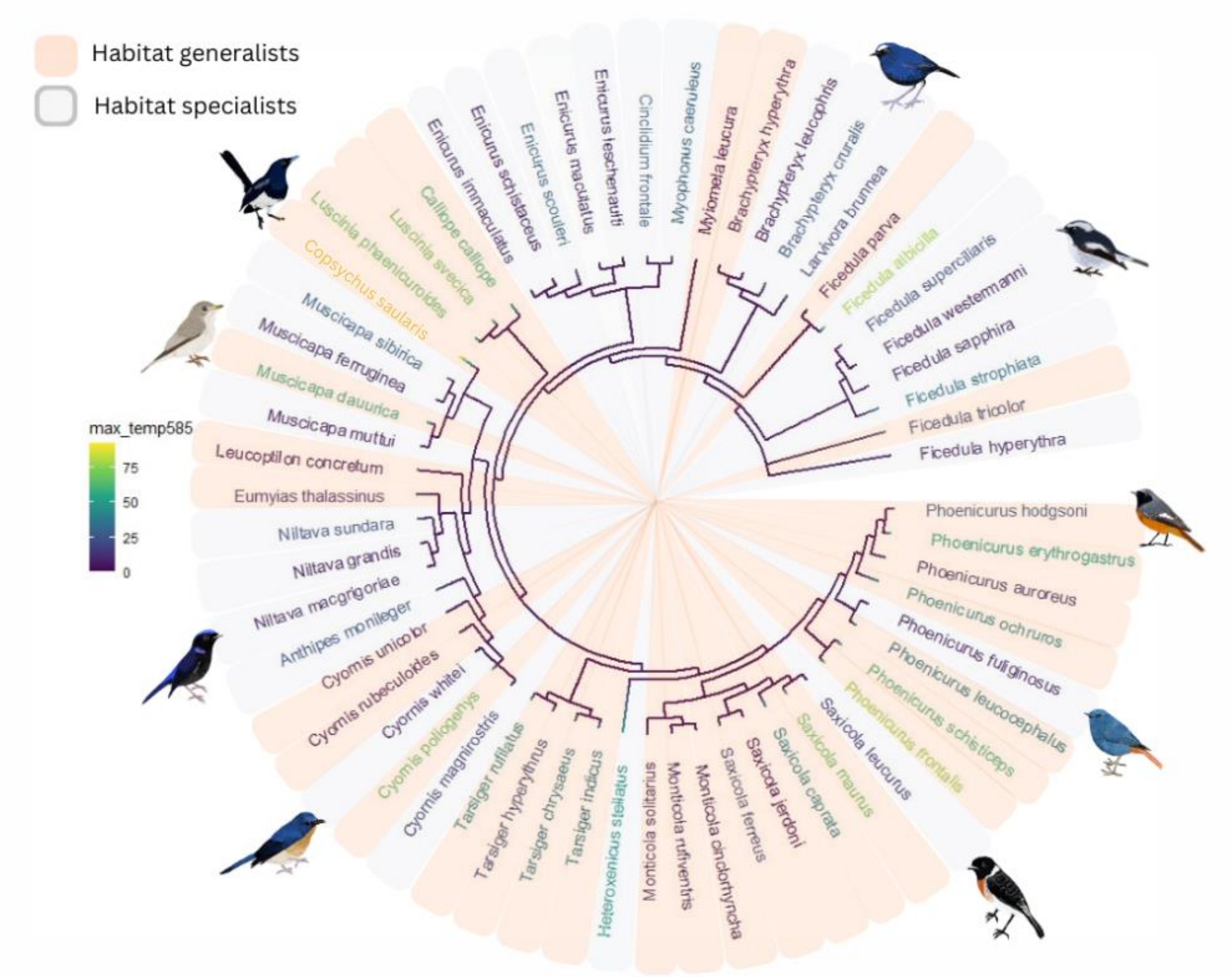


Figure S62: Percentage contribution of tmax (maximum temperature) under future climate scenario- SSP5-8.5 (high) to Ecological Niche Models of Muscicapidae (Flycatchers) species categorised as habitat specialists (light blue) or habitat generalists (light orange)

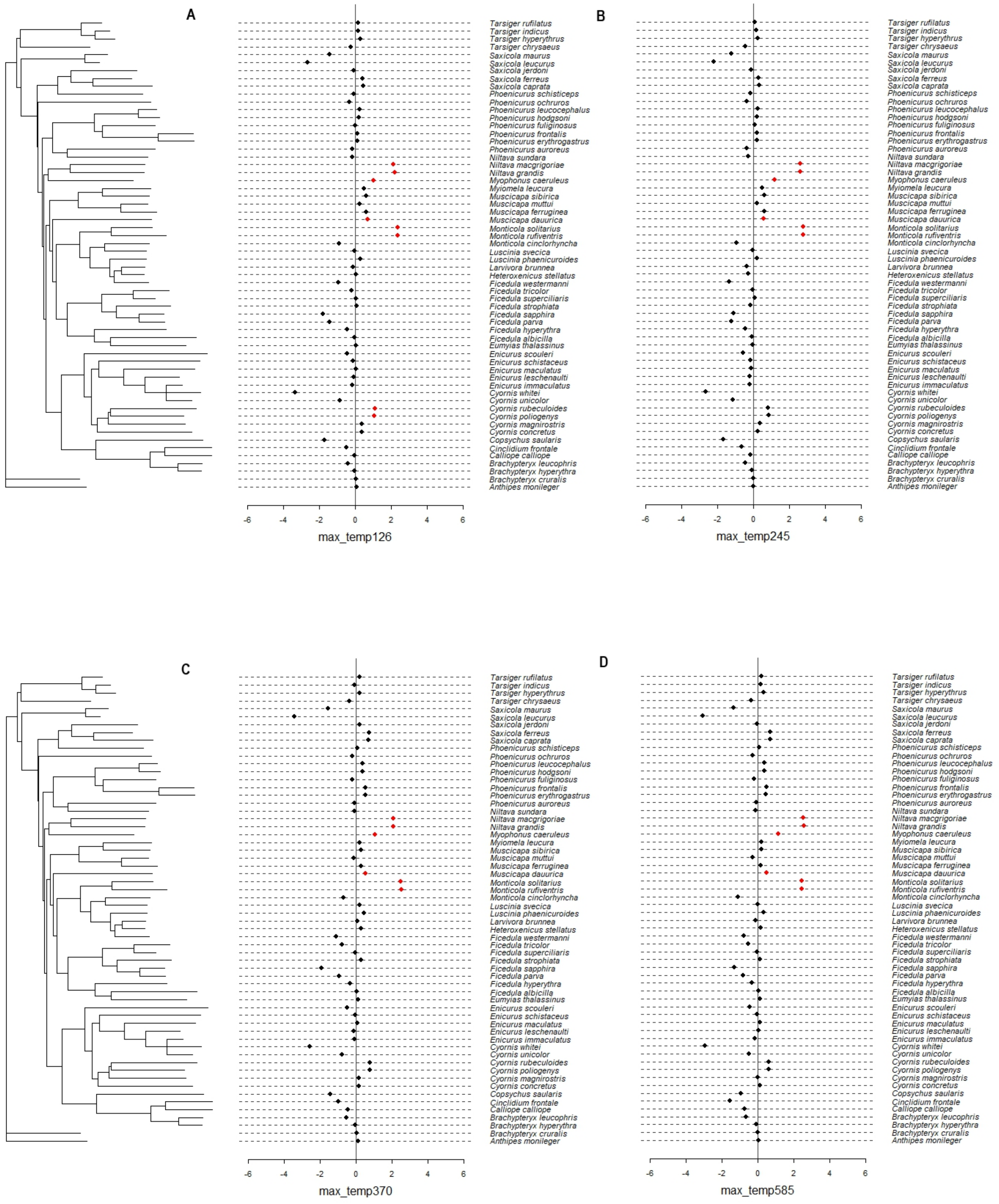


Figure S63: Local Moran's *I* plot of phylogenetic signal in percentage contribution of maximum temperature (tmax) under different emission scenarios- (A) SSP1-2.6 (low), (B) SSP2-4.5 (intermediate), (C) SSP 4-6.0 (medium) and (D) SSP5-8.5 (high). The red dots falling on and side of zero line correspond to significant relationships indicating positive correlation
